# Putting BASIL in a BLT: A Bayesian Filtering Method for Estimating the Fitness Effects of Nascent Adaptive Mutations

**DOI:** 10.1101/2025.03.29.646120

**Authors:** Huan-Yu Kuo, Sergey Kryazhimskiy

## Abstract

The distribution of fitness effects (DFE) of new beneficial mutations is a key quantity that dictates the dynamics of adaptation. The barcode lineage tracking (BLT) approach is an important advance toward measuring DFEs. BLT experiments enable researchers to track the frequencies of ∼10^5^ of barcoded lineages in large microbial populations and detect up to thousands of nascent beneficial mutations in a single experiment. However, reliably identifying adapted lineages and estimating the fitness effects of driver mutations remains a challenge because lineage dynamics are subject to demographic and measurement noise and competition with other lineages. We show that the commonly used Levy-Blundell method for analyzing BLT data and its improved version FitMut2 can produce biased fitness estimates, particularly if selection is strong. To address this problem, we develop a new method called BASIL (<u>BA</u>yesian <u>S</u>election <u>I</u>nference for <u>L</u>ineage tracking data), which dynamically updates the belief distribution of each lineage’s fitness and size based on the number of barcode reads. We calibrate BASIL’s model of noise with new experimental data and find that noise variance scales non-linearly with lineage abundance. We test how BASIL and FitMut2 perform on simulated data and on down-sampled data from the original BLT data by Levy et al and find that BASIL is both more robust and more accurate than FitMut2. Our work paves the way for a systematic inference of the distribution of fitness effects of new beneficial mutations from BLT experiments in a variety of scenarios.

**AUTHOR SUMMARY:** Beneficial mutations are rare but they are the ultimate drivers of evolution by natural selection. Evolutionary biologists seek to understand how many beneficial mutations an organism has access to in different environments and how these mutations affect fitness. Barcode lineage tracking (BLT) is a powerful experimental approach that tracks the frequencies of hundreds of thousands of subpopulations labeled with unique DNA barcodes and provides data that potentially enables researchers to identify and isolate many beneficial mutations arising in experimental microbial populations. However, analyzing these data is challenging because of the randomness of evolution and measurement noise. We found that existing methods for analyzing BLT data can lead to biased estimates of the fitness effects of beneficial mutations, especially when selection is strong. To overcome this issue, we developed a new method called BASIL, which uses a Bayesian approach that updates the estimated fitness and size of each lineage based on the measured barcode counts. We show that BASIL provides more accurate and robust estimates of the fitness effects of beneficial mutations in both simulated and real datasets than the existing alternatives. Thus, BASIL will facilitate a better understanding of beneficial mutations and adaptation more generally.

## INTRODUCTION

Every new adaptation originates in a single individual. Most vanish soon thereafter. The few that survive may encounter fierce competition with other contending adapted lineages [1–5]. This competition—termed “clonal interference”—is particularly severe in organisms that have limited recombination and also have a large supply of beneficial mutations, such as many bacteria [6–8], cancers [9,10], and viruses [11–13]. In the clonal interference regime, the chance of fixation of a new allele depends not only on its own fitness benefit but also on the benefits provided by competing alleles [14–16]. Thus, to understand the dynamics of rapid adaptation, we must know the distribution of fitness effects (DFE) of adaptive mutations [16,17]. However, this empirical knowledge is not yet readily available.

Measuring the effects of new adaptive mutations is challenging. The most statistically sound strategy would be to isolate them from random samples of spontaneous mutations. However, in most organisms, mutations are rare [18], and beneficial mutations typically constitute a relatively small fraction of all mutations [3,19,20] (but see [21,22]). As a result, obtaining accurate statistics of beneficial mutations from samples of random mutations is generally extremely inefficient. Instead, to sample adaptive mutations, researchers leverage the power of natural selection, which elevates the frequencies of beneficial mutations in a population. It is relatively easy to find and isolate beneficial mutations from recently adapted populations, particularly those evolved in the lab [23–26]. However, this strategy preferentially captures mutations with the strongest fitness benefits because they are least likely to be lost by genetic drift or clonal interference. Therefore, this approach may provide an incomplete view of the diversity of adaptive mutations [3]. Moreover, this view may also be biased because which mutations survive and fix depends in a complex way on the DFE itself and on population size and structure [16,27,28].

The barcode lineage tracking (BLT) method is a powerful strategy for estimating the fitness effects (also referred to as selection coefficients) of many nascent beneficial mutations in asexual genetically tractable microbes [3–5,29–36]. The key idea underlying BLT is to introduce many (typically ∼10^5^) random neutral DNA barcodes into an otherwise isogenic population and track barcode frequencies over time using deep sequencing [3,37]. Each new adaptive mutation that arises in the population is permanently linked to a unique DNA barcode. As an adaptive mutation spreads in the population, the corresponding increase in the frequency of the linked barcode can be detected in the sequencing data. Moreover, since the sizes of most barcoded subpopulations are initially small (typically ∼100 individuals), the expansion of each adapted subpopulation is initially driven by a single adaptive mutation. In other words, for a period of time, each barcode with increasing frequency reports on a single adaptive driver mutation. As a result, this approach allows one to measure the fitness benefits of many simultaneously segregating driver mutations, including weak ones, in a single relatively short evolution experiment, while avoiding laborious genetic reconstructions and screens [3,38]. Although the BLT approach does not entirely eliminate biases introduced by natural selection, it significantly mitigates them, and overall provides a much more accurate picture of the diversity of adaptive mutations than other existing methods [3].

While the BLT approach significantly advances our ability to capture and study adaptive mutations, analyzing BLT data remains difficult. The main challenge is to distinguish lineages that have acquired a single beneficial mutation (“adapted lineages”) from those that have not (“neutral lineages”) based on noisy dynamics of low-abundance barcodes. The core idea is the same as in classical competition assays [39–42], i.e., to determine whether a lineage systematically increases in frequency relative to neutral reference lineages (i.e., those with wildtype fitness) and then to infer the fitness effect of the underlying driver mutation from the rate of this increase. However, despite many similarities, the BLT setup differs in three important ways from the canonical competition assay. First, in competition assays, lineages are usually represented by thousands of cells at the beginning of the experiment, whereas in the BLT experiments all lineages are by design present at low abundances (∼100 cells), which makes lineage extinctions much more likely. Second, competition assays are typically short (≲ 50 generations) to prevent new mutations from significantly affecting lineage dynamics, whereas BLT experiments are longer (> 100 generations) precisely to allow for new adaptive mutations to arise and reach sufficiently high frequencies. Third, in competition assays, designated neutral reference lineages are deliberately added to the population. Their frequency dynamics reports on population’s mean fitness which allows one to estimate the fitness of all other lineages relative to the reference [22,41]. In contrast, in the BLT setup, all lineages are initially equivalent, and since adaptive mutations can arise during the experiment (or shortly before), which lineages become adapted and which ones remain neutral is a priori unknown.

The absence of defined neutral reference lineages is perhaps the biggest challenge of analyzing BLT data because such lineages provide the most obvious and robust way to readout the mean fitness of the population, which in turn is required for the inference of fitness of all other lineages. Two main approaches addressing this problem have been proposed so far. Levy et al developed the original approach, which we refer to as the “Levy-Blundell” or the “LB” method for short [3]. They chose low-abundance lineages as the neutral reference, reasoning that such lineages are unlikely to acquire adaptive mutations during the BLT experiment, and inferred mean fitness based on the rate of decline in the frequency of such lineages. However, low-abundance lineages are likely to go extinct, especially if selection is strong. One could choose more abundant lineages as the neutral reference, but such lineages are more likely to carry adaptive mutations, especially later in the experiment. Both outcomes are undesirable as they can bias the estimates of fitness effects of all adaptive mutations. More recently, Li et al developed an approach called FitMut2, which retains many features of the original LB method but instead of relying on designated neutral reference lineages it iteratively switches between identifying adapted lineages and estimating mean fitness on their basis until the process converges [43]. They have shown that FitMut2 outperforms the original LB approach on their simulated data. However, FitMut2 could also be prone to biases because the solution to which it converges might be sensitive to initial conditions and/or noise in the data.

Given these potential theoretical concerns, it is important to assess the performance of existing approaches empirically on simulated data where the ground truth is known, and to understand when and why they fail. In particular, the ability to accurately infer fitness effects will likely depend on the strength of selection because stronger selection accelerates evolutionary dynamics and reduces the amount time available for sampling lineage trajectories. To this end, in the first part of this paper, we assess the performance of the LB and FitMut2 approaches on BLT data simulated under weak and strong selection. We find that both approaches perform well under weak selection but produce substantially biased fitness estimates under strong selection. Since the LB method has been more widely used [3,4,29,31,32,35], we carry out a deeper investigation of the underlying reasons for its poor performance.

In the latter part of the paper, we develop a new BLT analysis method termed BASIL (<u>BA</u>yesian <u>S</u>election <u>I</u>nference for <u>L</u>ineage tracking data), which keeps track of belief distributions for the fitness of all lineages under the assumption that most lineages are initially neutral. It updates these distributions based on barcode read counts, and uses random lineages (both putatively adapted and neutral) to estimate mean fitness. Our approach combines several techniques developed previously for the analyses of competition assays [42,44–46] and BLT experiments [3,5,43]. In particular, similar to previous methods, BASIL models the evolutionary dynamics of lineages between sampling time points as well as the measurement process, whose properties we measure using a new calibration experiment. In contrast to most other approaches and similar to Ref. [45], BASIL treats unobserved lineage sizes as hidden variables, which is important for obtaining accurate estimates of lineage fitness, particularly under strong selection. Similar to Ref. [42], we cast our model in the Bayesian framework, which allows us to keep track of uncertainties in our fitness and lineage size estimates and dynamically update our belief distributions as the data arrives. We then directly compare the performance of BASIL on published BLT datasets to the performance of FitMut2. Finally, we use BASIL to characterize adaptation of several strains of yeast *Saccharomyces cerevisie* to various environments.

## RESULTS

### Existing BLT analysis methods can produce biased fitness estimates

To understand how accurately the LB and FitMut2 methods infer the fitness effects of nascent adaptive mutations, we simulate a batch-culture BLT experiment with 10^5^ barcoded lineages, mimicking the setup of Levy et al [3]. The details of our simulations are provided in Materials and Methods. Briefly, each batch-culture cycle begins with dilution, which we simulate by down-sampling each lineage to 1/*D* = 1/256 of its size using the Poisson distribution. Some lineages may go extinct at this step. Then, all surviving lineages deterministically expand by a factor proportional to the difference between their fitness and the population’s mean fitness, so that the population reaches approximately its pre-dilution size. Our simulations continue for 20 cycles, corresponding to 160 generations. Every other cycle, we simulate barcode sequencing by randomly sampling a certain number of individuals from our population and recording their lineage identities.

In our simulations, 3,000 lineages are adapted, i.e., all individuals within these lineages have the same fitness advantage compared to the rest of the population. This scenario where all beneficial mutations are pre-existing is less complex than many real BLT experiments, and inference methods should perform best in this case. Our reasoning for using these relatively simple simulations was twofold. First, many if not most adaptive mutations detected in BLT experiments arise prior to the beginning of the experiment [3,4,33]. Second, we reasoned that if we find that a method fails on these relatively simple simulated data, it is very unlikely to perform well on real BLT datasets. Since there are no new mutations, our simulations are similar to competition assays, with the exception that the identity of neutral and adapted lineages is unknown to the inference algorithm. We simulate two conditions that we refer to as “strong selection” and “weak selection” (Figure S1 and Table S3). In the weak selection regime, which resembles the original BLT experiment carried out by Levy et al [3], the average fitness of an adapted lineage is 3%. As a result, the mean fitness of the population increases by about 4% in 150 generations. In the strong selection regime, the average fitness of an adapted lineage is 8%, and the population’s mean fitness increases by about 12% in 150 generations.

Our first goal is to assess how the LB and FitMut2 methods perform on these simulated data. If the methods underperform, we would like to understand why, focusing specifically on the better-established LB method. To enable a potential in-depth investigation, we simplify the original LB approach by stripping away much of the complexity required for analyzing real BLT datasets but unnecessary for the analysis of our simulated data, while retaining its core assumptions (see Materials and Methods and Section 2.3 of the Supplementary Information). We refer to this simplified version as the “neutral decline” method to distinguish it from the original. As the original LB approach, the neutral decline method requires us to specify which lineages we select as the neutral reference. To determine how this choice might affect inference, we select either low- or high-abundance lineages, with low-abundance lineages being defined as those represented by 20 to 40 reads (corresponding to 20 to 40 cells at the bottleneck in our simulations) and high-abundance lineages being defined as those represented by 80–100 reads (corresponding to 80 to 100 cells at the bottleneck in our simulations).

The performance of both the neutral decline and FitMut2 methods on our BLT simulations are illustrated in Figure 1. We find that in the weak selection regime, both FitMut2 and the neutral decline method with high-abundance reference lineages accurately infer the mean fitness trajectory and the fitness effects of individual adaptive mutations (Figure 1E,F,I). When low-abundance lineages are used as reference, the neutral decline approach underestimates both the mean fitness of the population and the selection coefficients of individual adaptive mutations even in this favorable regime (Figure 1A,B). Both methods perform significantly worse in the strong selection regime (Figure 1C,D,G,H,K,L). FitMut2 and the neutral decline approach with high-abundance reference lineages severely underestimate both the mean fitness and the effects of individual mutations (Figure 1G,H). If low-abundance lineages are used as reference, the neutral decline method estimates the mean fitness accurately, but only for the first ∼50 generations (Figure 1C), which results in noisy estimates of selection coefficients of individual adaptive mutations (Figure 1D).

**Figure 1.**
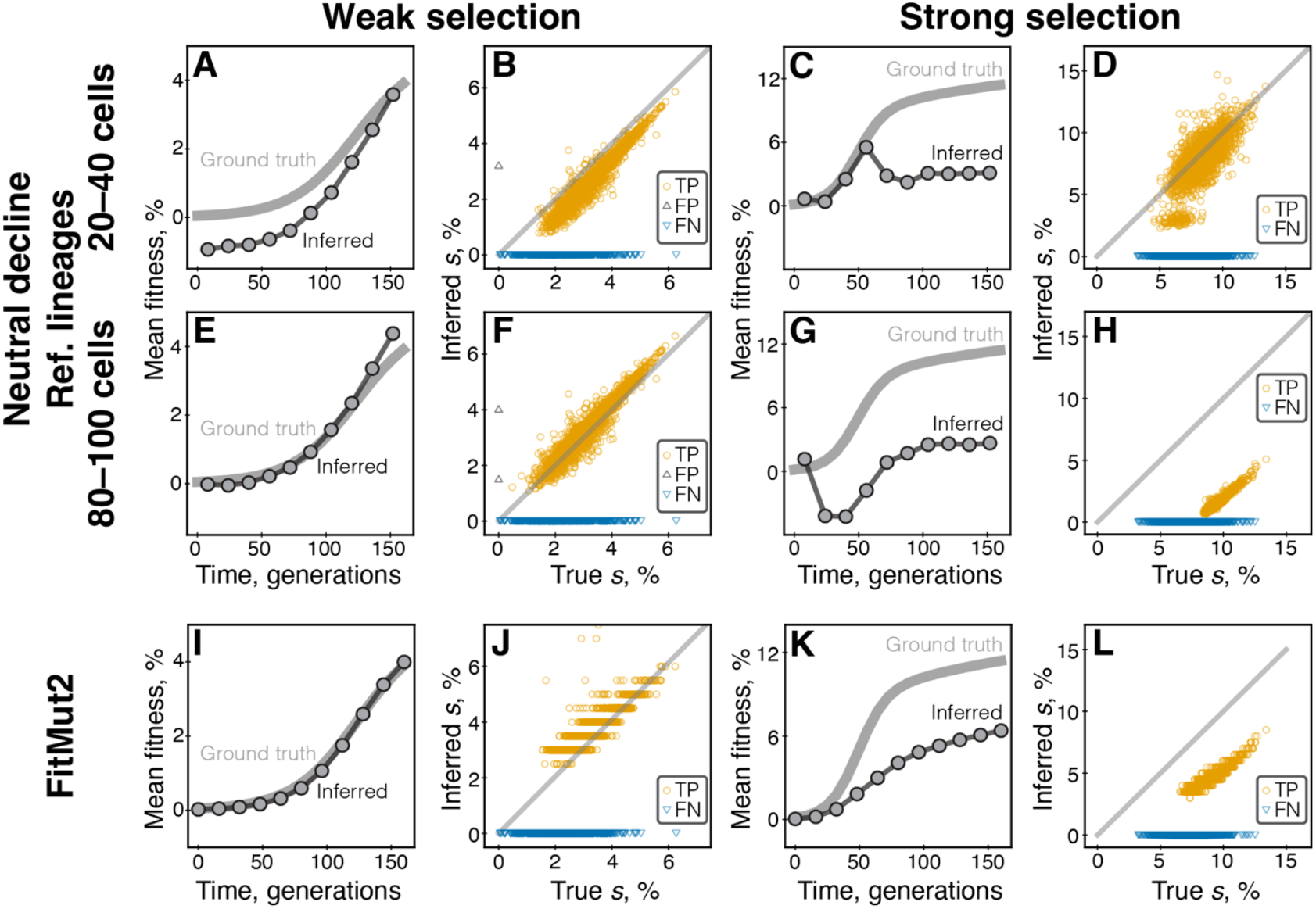
Performance of the neutral decline and FitMut2 methods on simulated BLT data depends on the strength of selection. Panels A–H show the results for the neutral decline method, and panels I– L show the FitMut2 results. Left panels A, B, E, F, I, J show simulations in the weak selection regime; right panels C, D, G, H, K, L show simulations in the strong selection regime. **A, C.** Mean fitness trajectories, true and inferred by the neutral decline method with low-abundance reference lineages. **B, D**. Selection coefficients of adapted lineages, true versus inferred by the neutral decline method with low-abundance reference lineages. Each symbol corresponds to a lineage; orange circles, black upward triangles and blue downward triangles represent true positives, false positives and false negatives, respectively; true negatives are not shown. **E–H**. Same as A–D but for high-abundance lineages. **I–L**. Same as A–D but for FitMut2.

Thus, the performance of existing BLT analysis methods depends on the conditions of the BLT experiment, in particular on the strength of selection. Furthermore, the accuracy of the neutral decline method—and by extension that of the LB method—hinges on the choice of reference lineages.

### Statistical causes for bias of the LB method

To understand why the neutral decline method (and, consequently, the LB method) sometimes produces biased estimates of lineage fitness, we first summarize the central assumption underlying this approach (see Materials and Methods and Table S4 for more details). To infer the population mean fitness 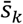 at the sampling interval (*t*_*k*−1_, *t*_*k*_), the neutral decline method groups together reference lineages with each reads count *r*_*k*−1_ observed at *t*_*k*−1_. For each such group, it computes the average read count 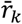 at the next sampling time *t*_*k*_ and infers mean fitness 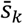 from the equation

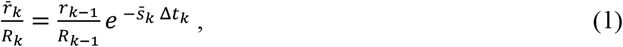

where *R*_*k*_ is the total read depth at time *t*_*k*_ and Δ*t*_*k*_ = *t*_*k*_ − *t*_*k*−1_. Equation (1) can be derived from the standard population genetics theory under the crucial assumption that all lineages with the same number of reads *r*_*k*−1_ are present in the population at frequency *r*_*k*−1_/*R*_*k*−1_ at *t*_*k*−1_ [47]. We refer to equation (1) as the “neutral decline” equation. Once the mean fitness trajectory 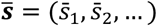 is known, the neutral decline method infers the selection coefficients of all non-reference lineages by maximizing the likelihood of their frequency trajectories (for details, see Materials and Methods). Importantly, even though the original LB method is more sophisticated in that it utilizes the entire conditional distribution of lineage read counts at the next sampling time given their read counts at the previous time point (see equation (45) in the Supplementary material to Ref. [3] or equation (S8) in the Supplementary Information), the mean of that distribution is given by equation (1). Therefore, a failure of equation (1) to capture the relationship between *r*_*k*–1_ and 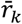 would imply a failure of the original LB model.

To test the validity of equation (1), we plot the rescaled average read count 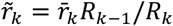 at time point *t*_*k*_ against the corresponding observed read count *r*_*k*–1_ at the previous time point. According to equation (1), 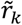 must be proportional to *r*_*k*−1_ with a zero *y*-intercept. Instead, we find that this relationship is more complex, with its shape being dependent on the selection regime and on time (Figure 2A,C). Specifically, when selection is weak, 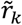 relates linearly to *r*_*k*−1_ during the entire BLT experiment, but with a non-zero *y*-intercept (Figure 2A). Despite its relatively small value (e.g., 6.60 at generation 16 and 2.89 at generation 160), a non-zero *y*-intercept can cause significant underestimates of mean fitness when low-abundance lineages are chosen as reference (Figure 2B), including negative estimates at early time points (see Figure 1A). Since the estimates of selection coefficients of all lineages depend on the estimate of mean fitness, the bias in the latter propagates to a bias in the former, as seen in Figure 1B. Choosing initially more abundant lineages as reference produces a more accurate estimate of initial mean fitness (Figures 1E and 2B) because such estimates are less sensitive to the value of the *y*-intercept (Figure 2A). However, choosing more abundant lineages is not universally better. For example, under strong selection, the relationship be-tween 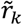 and *r*_*k*−1_ becomes non-linear for higher-abundance lineages, particularly later in the experiment (Figure 2C), which also biases the estimates of mean fitness (Figures 1C,G and 2D).

**Figure 2.**
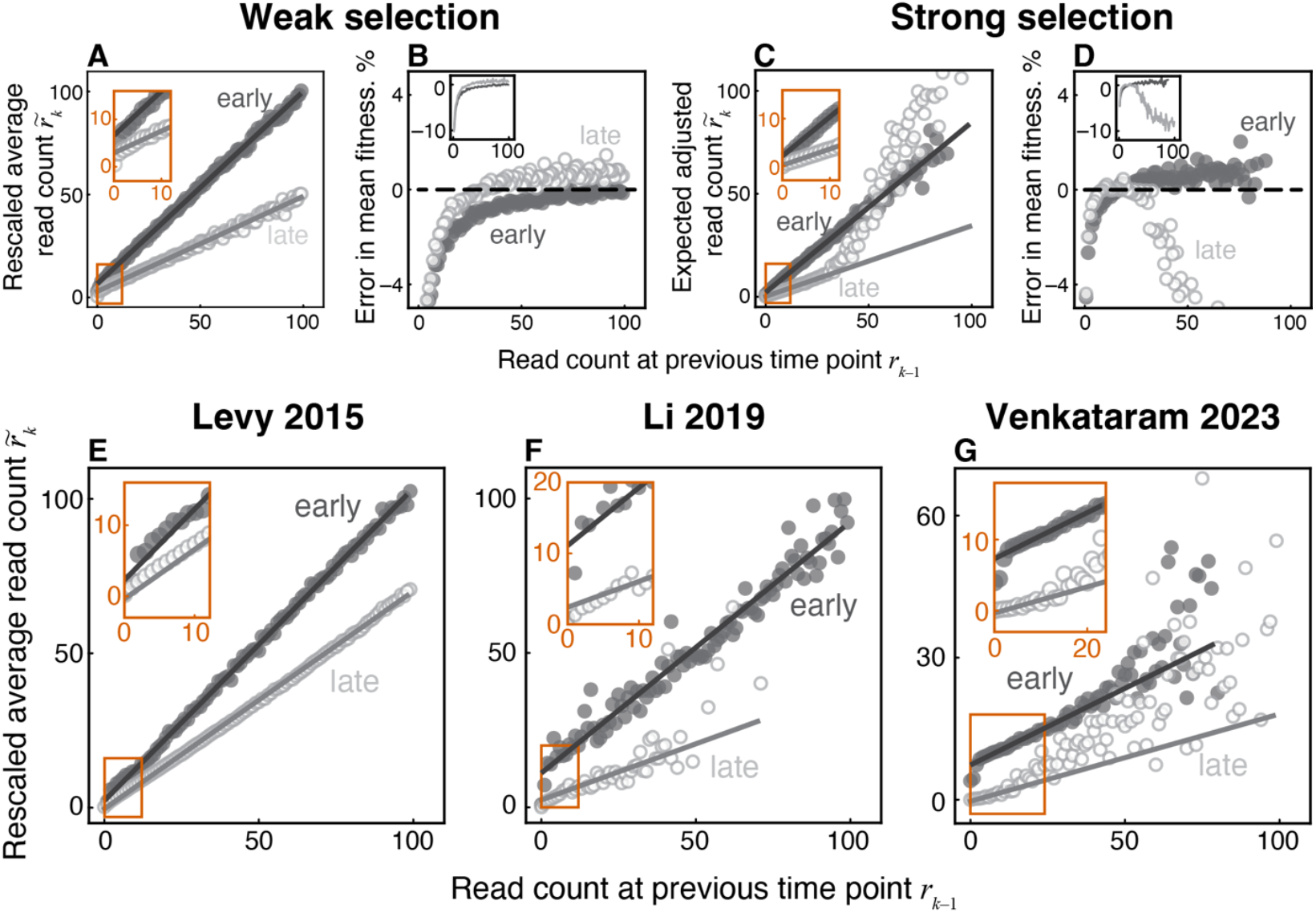
Observed lineage read counts deviate from those predicted by equation (1) both in simulated and real data. A. Rescaled average read count 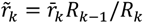 at the next time point (ordinate) plotted against the read count observed at the previous time point *r*_*k*−1_ (abscissa) for the weak-selection simulation. Light and dark gray points represent early (*t*_*k*_ = 16 generations) and late (*t*_*k*_ = 160 generations) time intervals. Lines represent least-squares best fits within the linear regime (see Materials and Methods). **B.** Error in the inferred mean fitness (inferred minus true) as a function of the read count of lineages that are used as the neutral reference. Shades are the same as in panel A. **C**. Same as panel A but for the strong selection regime. Early time interval is at *t*_*k*_ = 16 generations and late time interval is at *t*_*k*_ = 64 generations. **D**. Same as panel B but for the strong selection regime. **E–G**. Same as panel A but for three real BLT datasets: Levy 2015 R1 (panel E), Li 2019 Evo1D R2 (panel F) and Venkataram 2023 Co-evolution R5 (panel G). Early time interval is at *t*_*k*_ = 16, 7, 20 generations in panels E, F, G, respectively and late time interval is at *t*_*k*_ = 112, 133, 86 generations.

These observations show that the relationship between lineage read counts at successive time points can be used to diagnose potential inference problems in the absence of the ground truth. Thus, we examined these relationships in real BLT data, asking whether they exhibit similar deviations from equation (1) as we observed in our simulations. To this end, we reanalyzed data from three published BLT studies, Levy 2015 [3], Li 2019 [29] and Venkataram 2023 [33]. We find strong deviations from linearity in the Li 2019 and Venkataram 2023 data for higher-abundance lineages at later time points (Figures 2 F,G). Even at the initial time point where we expect equation (1) to be most accurate, we find that, even though 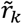 depends on *r*_*k*−1_ linearly, there is a statistically significant positive *y*-intercept (2.26 for Levy 2015, 11.10 for Li 2019, and 7.32 for Venkataram 2023, all *P*-values < 10^−12^, F-test; Figure 2E–G). As a result, the estimates of mean fitness and selection coefficients of adaptive mutations in real data produced by the LB method are likely subject to the same biases as observed in simulated data sets.

Finally, we asked why the observed relationship between *r*_*k*–1_ and 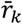 deviates from the prediction given by equation (1). First, the deviation from linearity at high *r*_*k*–1_, especially later in the experiment (Figure 2C,F,G), can be explained if reference lineages are in fact not neutral. Indeed, after selection had time to act, lineages that carry stronger beneficial mutations are typically present at higher frequencies in the population and hence have higher values of *r*_*k*–1_ than neutral lineages. As a result, lineages with higher *r*_*k*–1_ increase disproportionately more or decrease disproportionately less than lineages with lower *r*_*k*–1_. Second, to understand why the relationship between *r*_*k*–1_ and 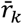 has a non-zero *y*-intercept, consider a hypothetical example where all lineages are neutral and equally abundant in the population and the coverage at both time points *t*_*k*−1_ and *t*_*k*_ is such that a typical lineage is represented by 10 reads. Then, at *t*_*k*−1_, while most lineages are represented by *r*_*k*− 1_ = 10 reads, measurement noise will result in many lineages with *r*_*k*−1_ = 9, *r*_*k*−1_ = 11, *r*_*k*−1_ = 8, *r*_*k*−1_ = 12, etc. Since all lineages are neutral, the expected read count for any lineage at the next time point is 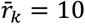, including those with *r*_*k*−1_ ≠ 10, whereas the neutral decline equation (1) predicts 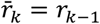. This bias arises as a result of the regression to the mean, whereby lineages whose read counts are by chance abnormally low at one time point are represented by a more typical (larger) number of reads at the next time point, causing the *y*-intercept to be non-zero (see Supplementary Information, Section 2.3.2 for an extended discussion). The specific value of the intercept depends, among other things, on the measurement noise and on the distribution of lineage frequencies in the population, quantities that are a priori unknown. Therefore, correcting for this intercept appears difficult.

Overall, this investigation demonstrates that the neutral decline approach (and, consequently, the LB method) can lead to biased inferences of the fitness effects of nascent adaptive mutations because equation (1) (or the corresponding equation (S8) for the full distribution of reads) is in general incorrect, particularly for low-abundance lineages or under strong selection.

### BASIL: Bayesian selection inference for lineage tracking data

To overcome challenges of the existing approaches, we developed BASIL (<u>Ba</u>yesian <u>S</u>election <u>I</u>nference for <u>L</u>ineage tracking data), a robust statistical method for identifying adapted lineages and inferring their fitness and implemented it in a software package [48]. The key hidden variables in BASIL are the sizes *n* and selection coefficients *s* (fitness) of individual lineages. Population’s mean fitness 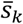 and the measurement noise parameter *ϵ*_*k*_ defined below are time-dependent global variables. At any given sampling time *t*_*k*_, each lineage *i* is characterized by the joint belief distribution 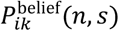 for its size and fitness, which we model parametrically as a product of a normal and gamma distributions (see Materials and Methods and equations (S31)−(S33) in the Supplementary Information). BASIL is a Bayesian filtering model (Figure 3B, [49]), in which the belief distributions 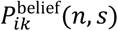 are based on the past data, i.e., data accumulated up to and including time *t*_*k*_.

**Figure 3.**
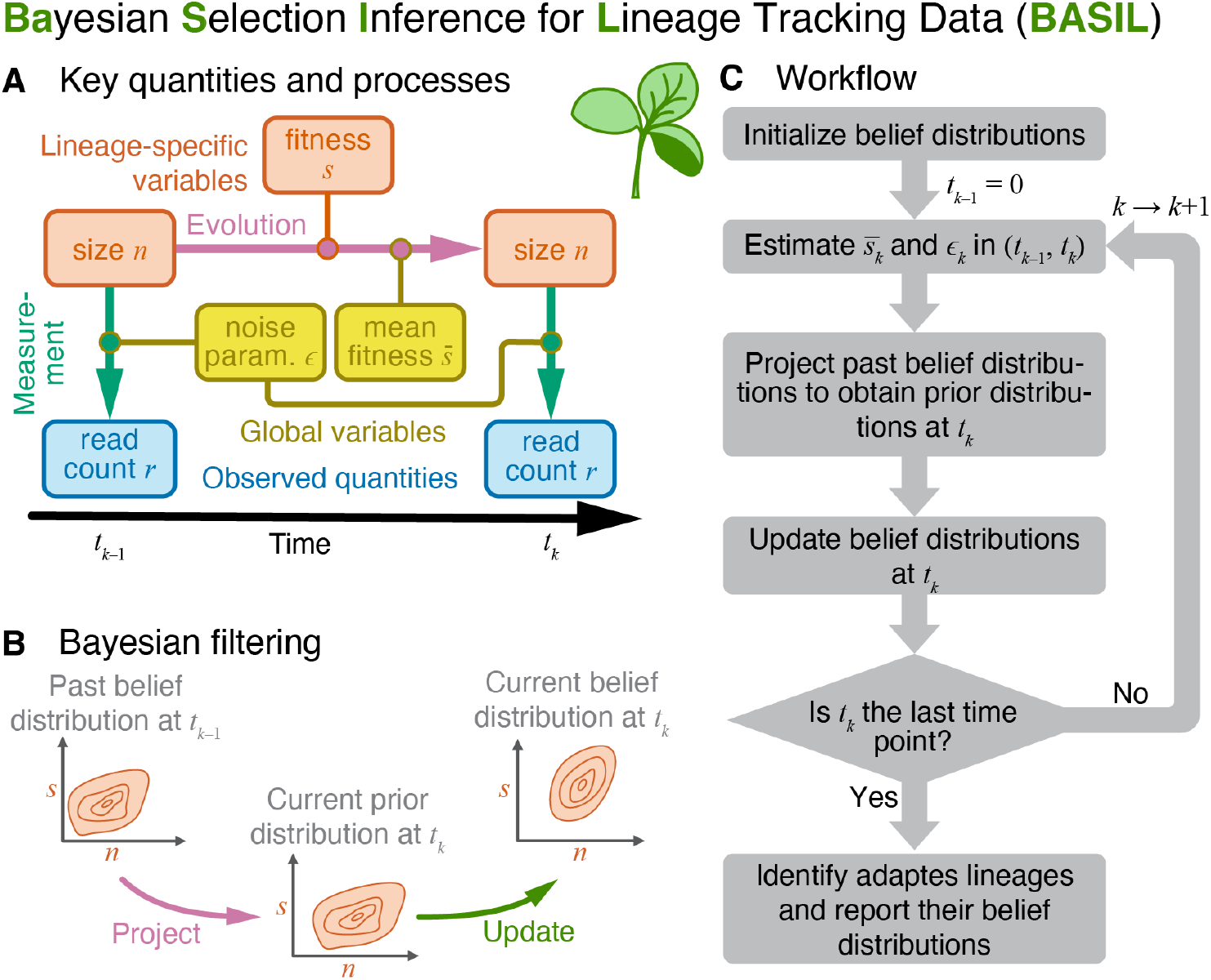
BASIL schematic. **A.** Key quantities and processes. Each barcoded lineage *i* is characterized at the current time *t*_*k*_ by its size *n*_*ik*_ and fitness (selection coefficient) *s*_*ik*_, which are unknown. Lineage size changes over time depending on the lineage fitness and the mean fitness of the population. At each sampling time point, we measure relative lineage sizes by counting sequencing reads with the corresponding barcode. **B**. Bayesian filtering consists of projecting the past belief distribution over the hidden variables *n*_*k*–1_ and *s*_*k*–1_ using the model of evolution to obtain the prior distribution for *n*_*k*_ and *s*_*k*_ for the current sampling time point *t*_*k*_ and then updating this distribution using read count *r*_*k*_ to obtain the new belief distribution. **C**. BASIL workflow.

At BASIL’s core is the model of evolutionary dynamics that computes the prior distribution 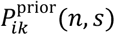 for the size and selection coefficient of each lineage *i* at the current sampling time *t*_*k*_ based on the past belief distribution 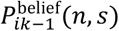 and the population’s mean fitness 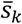 in the time interval (*t*_*k*−1_, *t*_*k*_) (Figure 3A). We also develop a model of measurement *P*^meas^(*r*|*n*: *ϵ*), which probabilistically relates the true lineage size *n* to the corresponding barcode read count *r* observed in the sequencing data. We model *P*^meas^(*r*|*n*: *ϵ*) as a negative binomial distribution that depends on the current value of the noise parameter *ϵ*.

Estimation of 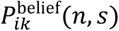 at *t*_*k*_ occurs in three steps (Figure 3C). First, we estimate the mean fitness of the population 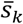 and the noise parameter *ϵ*_*k*_ in the current time interval (*t*_*k*−1_, *t*_*k*_) by sampling thousands of random lineages, computing their prior distributions 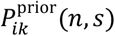 and then maximizing the likelihood of their observed read counts *r*_*ik*_ at *t*_*k*_. Second, we compute the prior distributions 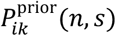 for all lineages, given the estimated population’s mean fitness 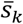. We then apply the Bayes theorem to obtain the updated lineage belief distributions 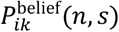, given the read counts *r*_*ik*_ and the estimated noise parameter *ϵ*_*k*_. In the rest of the section, we flesh out the main ideas and expressions behind our method. The full mathematical details are provided in Section 3 of the Supplementary Information.

### Model of evolution

Consider a barcoded lineage with fitness *s* relative to the ancestor and suppose that this lineage is represented by *n* _*k*−1_= *n*(*t* _*k*−1_) individuals in the population at the previous sampling time *t*_*k*−1_. On average, such lineage will expand if *s* is larger than the population’s current mean 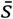 and it will contract if 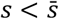. Demographic noise will also cause its size to randomly fluctuate around this expectation, such that the lineage size *n*_*k*_ at the current sampling time *t*_*k*_ is a random variable. To model it, we assume that our BLT population grows in the batch-culture regime [25] and sampling happens immediately prior to a dilution step. Then, during dilution, a fraction 1/*D* of the population is transferred into the fresh medium and the rest is discarded, where *D* > 1 is the dilution factor. Immediately after dilution, the size of the focal lineage becomes 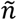, which we draw from the Poisson distribution with mean *n* _*k*−1_/*D*. Then, we assume that the lineage grows or shrinks exponentially and deterministically for the batch-culture cycle duration Δ*t*_*c*_, such that by the end of the cycle its size becomes 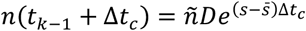. If the next sampling point is the end of this cycle, then *n*_*k*_ = *n*(*t* _*k*−1_ + Δ*t*_*c*_). Otherwise, we repeat the dilution and growth phases until the sampling time *t*_*k*_ is reached. Note that this model allows for the possibility of lineage extinction during dilution.

### Model of measurement

The actual lineage sizes are not observable. Instead, the population is sampled at time points *t*_0_, *t*_1_, … and sequenced at the barcode locus, which involves a series of processing steps [50]. Then the sequenced reads containing each barcode are counted. This measurement process can be described by a probability distribution *P*^meas^(*r*|*n*) of observing *r* reads for a barcode lineage with *n* cells. The simplest model for this process is the Poisson distribution with mean *nR/N*, where *R* is the read depth (i.e., the total number of reads obtained for the sample) and *N* is the total population size at the sampling point. However, sequencing read counts are often overdispersed with respect to the Poisson distribution [46,51]. Previously, Levy et al modeled measurement noise with a distribution that allows for overdispersion and in which variance scales linearly with the mean, similar to the Poisson distribution [3]. Other studies modeled read counts with a negative binomial distribution which permits an arbitrary scaling between mean and variance [52−54].

To determine how measurement noise variance scales with the mean read count, we performed the following calibration experiment. We assembled a barcoded population of *Saccharomyces cerevisiae* with a total size of 1.6×10^7^ individuals that consisted of 26 subpopulations present at six different frequencies (from 10^−5^ to 0.40) and sequenced it with 9-fold replication to an average depth of about 2×10^5^ reads per replicate (see Materials and Methods for details).

As expected, we found that the mean of the barcode frequency estimated from sequencing data is very close to the input frequency of each lineage (Figure 4A). To test the extent of overdispersion, we plotted the variance of the measured read counts 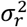 against the mean ⟨*r*⟩. If the measurement process was adequately described by the Poisson distribution, we would expect to observe a linear relationship with slope 1, whereas the noise model of Levy et al predicts linear scaling with a slope greater than 1 [3]. We find that the Poisson model fits our data quite well for read counts ≲100 but fails for more abundant barcodes (Figure 4B). Instead, the variance is better fit by a non-linear convex function of the mean. We capture this nonlinearity with the model

**Figure 4.**
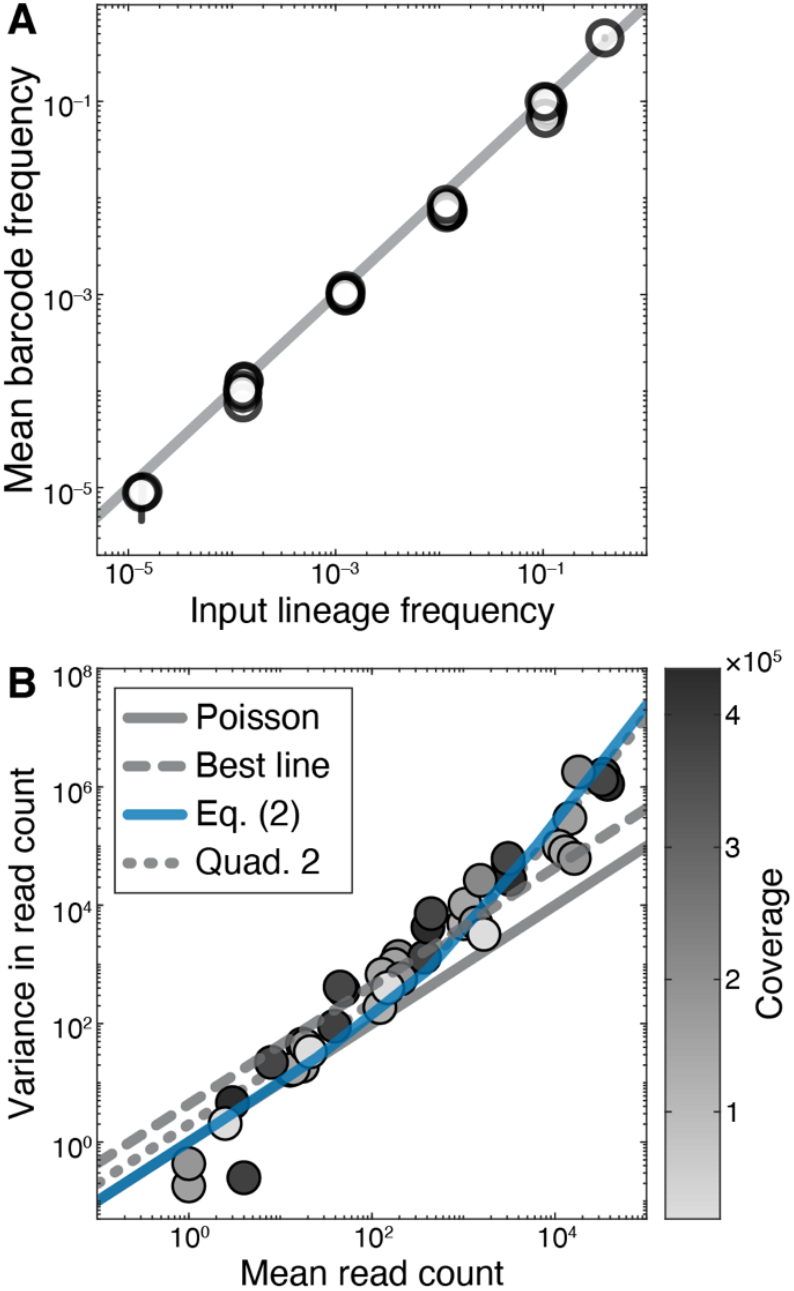
Properties of the measurement process. **A.** Barcode frequency estimated from barcode sequencing data (abscissa) versus the lineage frequency set experimentally (ordinate). Each point is a unique barcode. Error bars show one standard deviation of the mean. The 1-to-1 line is shown in gray. **B**. Variance of the read count versus the mean read count. Each point represents the data from multiple barcodes of the same lineage frequency. Shade represents coverage. Solid blue line is the parabola given by equation (2). Other curves are described in the Materials and Methods and their fitted parameter values are provided in Table S2.

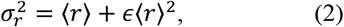

where *ϵ* is a measurement noise parameter that controls the degree of overdispersion (Figure 4B, blue line). We estimate that *ϵ* = 2.4×10^−3^ in the calibration experiment, which is significantly different from zero (Table S2). Since the specific value of *ϵ* might depend on experimental details, we consider equation (2) as a constraint on our model of noise, and allow *ϵ* to be a free parameter at each time interval. Since the functional form (2) requires a more flexible distribution than afforded by either the Poisson or the Levy et al distributions, we model the measurement process with a negative binomial (NB) distribution (equation (5) in Materials and Methods) with mean ⟨*r*⟩ = *nR*/*N* and the variance given by equation (2).

### Bayesian filtering

For each lineage *I*, we seek to estimate the joint belief distributions 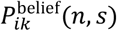 of lineage size *n* and fitness effect *s* at each sampling time *t*_*k*_. First, assuming that the past belief distribution 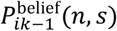 at *t*_*k*−1_ and the mean fitness 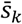 in the time interval (*t*_*k*−1_, *t*_*k*_) are known, we obtain the prior distribution 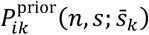 at the current time *t*_*k*_ by project-ing the past belief distribution 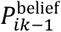 using our model of evolution. The prior distribution 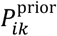 represents our best guess for the abundance of lineage *I* and its fitness at the current time point *t*_*k*_, based on its past trajectory and the population’s mean fitness. The observed read count *r*_*ik*_ for this lineage may or may not be consistent with this guess, and we use this new data to update our belief distribution using the Bayes’ theorem (Figure 3B),

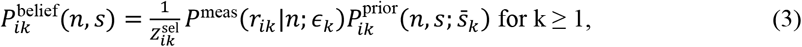

where *P*^meas^ is the NB distribution defined in the “Model of measurement” section and 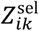 is the marginal probability that *r*_*ik*_ reads are observed at *t*_*k*_, given all of our prior knowledge (subscript *S* stands for “selection” and refers to the fact that this model allows for the fitness of the lineage to be non-zero). Note that 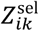 depends on the noise parameter *ϵ*_*k*_ and the mean fitness 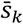 in the current time interval (*t*_*k*−1_, *t*_*k*_), which are assumed to be known.

### Initialization

We initialize the belief distribution for any lineage *i* as 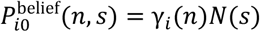, where γ_*i*_(*n*) is a Gamma distribution with parameters based on the lineage’s initial read count and *N*(*s*) is the normal distribution with zero mean and standard deviation 0.1. In other words, we assume that all lineages are initially most likely neutral, but with large uncertainty.

### Estimating population’s mean fitness

As mentioned above, our procedure for updating the belief distribution at time *t*_*k*_ depends on the mean population fitness 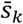 and the noise parameter *ϵ*_*k*_ at the time interval (*t*_*k*−1_, *t*_*k*_). To estimate 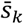 and *ϵ*_*k*_, we randomly choose 3,000 lineages and classify them as either putatively adapted (set *A*_*k*_) or putatively neutral (set *N*_*k*_). A lineage is classified as putatively adapted if the mean of its marginal belief distribution 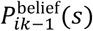 is positive and at least three times larger than its standard deviation; and it is classified as putatively neutral otherwise. We then estimate 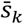 and *ϵ*_*k*_ by maximizing the log-likelihood function

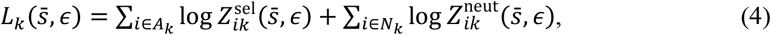

where 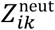 is a quantity analogous to 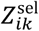 but for the neutral model where the selection coefficient of the lineage is zero.

### Identification of adapted lineages

As a result of the procedure described above, for each lineage *i*, we obtain a time-varying belief distribution 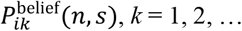 We calculate the final marginal belief distribution for the selection coefficient of lineage *i* as 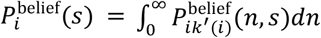 choosing the time point 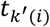 where the *s*-variance of the belief distribution 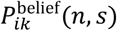 is minimal. The choice to minimize *s*-variance (rather than simply using the last available time point) is motivated by the fact that secondary adaptive mutations arising later in the experiment may cause widening of the belief distribution. While such widening may be biologically informative, the main purpose of BLT experiments is to estimate the effects of single nascent adaptive mutations, suggesting that one should select a time point before secondary mutations become sufficiently abundant. In practice, this choice appears unimportant, as the *s*-variance is minimized at the last time point for the overwhelming majority of lineages across all datasets that we analyzed (see Data S1, Tab 3).

Once we obtain the lineage’s final belief distribution 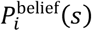, we call lineage *i* adapted if the mean *ŝ*_*i*_ of this distribution is sufficiently separated from 0, that is, if *ŝ*_*i*_ *≥ β σ*_*i*_, where *σ*_*i*_ is the standard deviation of 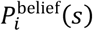. We refer to β as the “confidence factor”, a hyper-parameter that must be determined empirically. Intuitively, larger β would require a lineage to have a higher fitness to be recognized as adapted, which would increase precision but reduce recall, or, equivalently, reduce the rate of false positives at the expense of increasing the rate of false negatives. In other words, a higher confidence factor would allow us to be more confident that the lineages identified as adapted are in fact adapted, albeit at the expense of missing more adapted lineages for which evidence of adaptation is weaker. On the other hand, decreasing β would increase recall but reduce precision, or, equivalently, it would allow us to identify more adapted lineages, albeit at a cost of also misidentifying more neutral lineages as adapted. Recall and precision usually exhibit a trade-off, and there is no general principle for choosing β. In the next section, we test our method on simulated data, which provides us with a guideline for the value of the confidence factor.

### Comparison of methods on simulated data

For an initial assessment of BASIL performance and to determine the value of the confidence factor β, we applied BASIL to simulated BLT data described above (see Section “Estimation of mean fitness from the decline of neutral lineages can produce biased estimates of fitness effects” and Section 4 of the Supplementary Information). We found that BASIL accurately captures the dynamics of mean fitness in both regimes (compare Figures 1 and 5), even at relatively late time points where the other approaches fail. To determine the hyper-parameter β, we varied it between 0.0 and 6.4 with the step size of 0.1 and calculated the F1 score for adapted lineage calls, which is the harmonic mean of recall and precision (Figure 5E,G). We found that the F1 score is maximized at β = 3.4 and β = 3.2 for the weak and strong selection regimes, respectively. Thus, we chose the value of β = 3.3 for all our subsequent analyses (see Section 4.1 in the Supplementary Information and Figure S5). Using this confidence factor, we identified 2,570 adapted lineages (430 false negatives) in the weak selection regime, with 31 false positives, resulting in 98.8% precision and 85.7% recall. In the strong selection regime, we identified 2,481 adapted lineages (519 false negatives) with 7 false positives, resulting in 99.7% precision and 82.7% recall. For lineages identified as adapted, our fitness estimates 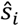 are both accurate and precise in both regimes. The full comparison between methods is provided in Table 1).

**Table 1.**
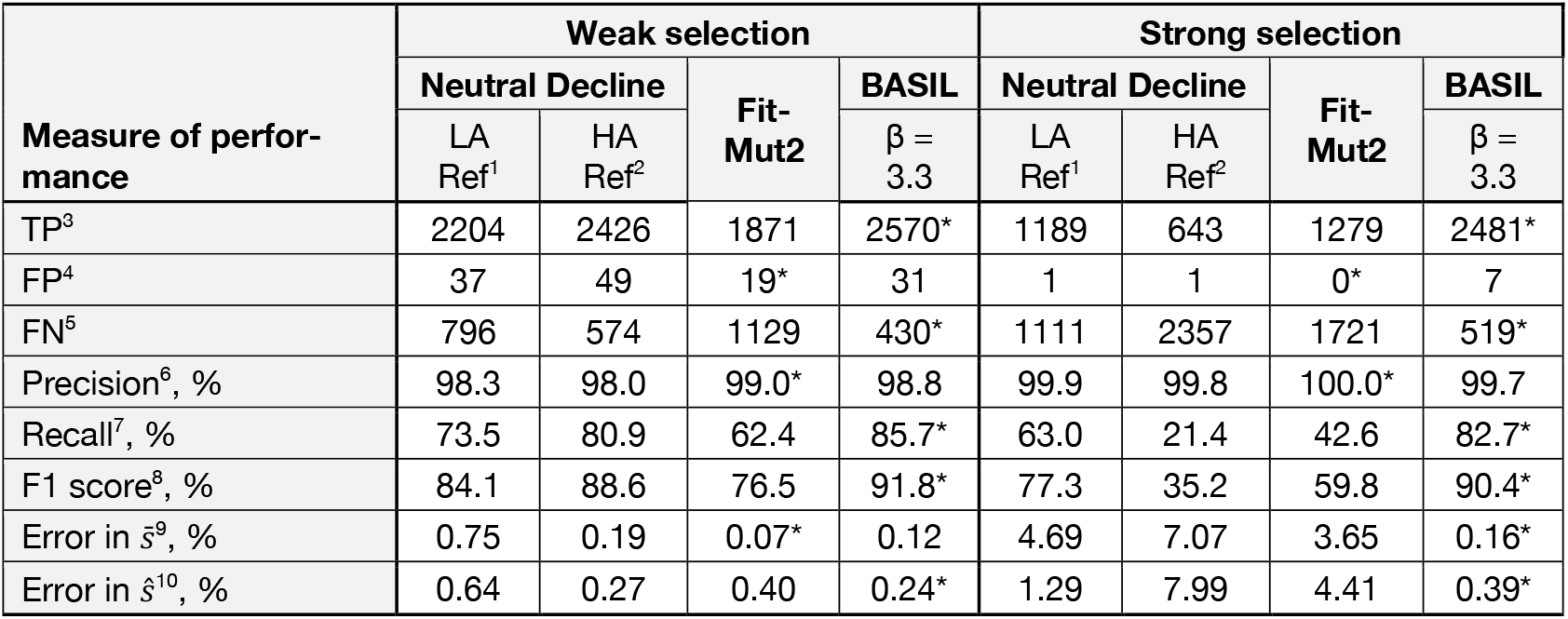
Comparison of BLT analysis methods on simulated data. ^1^Low-abundance (LA) lineages used as reference; ^2^High-abundance (HA) lineages used as reference; ^3^Number of true positives; ^4^Number of false positives; ^5^Number of true positives; ^6^Precision = TP/(TP+FP); ^7^Recall = TP/(TP+FN); ^8^F1 score = 2TP/(2TP+FP+FN); ^9^Absolute difference between true and inferred, averaged across time; ^10^Absolute difference between true and inferred fitness averaged over true positive lineages. *indicates the best performance.

**Figure 5.**
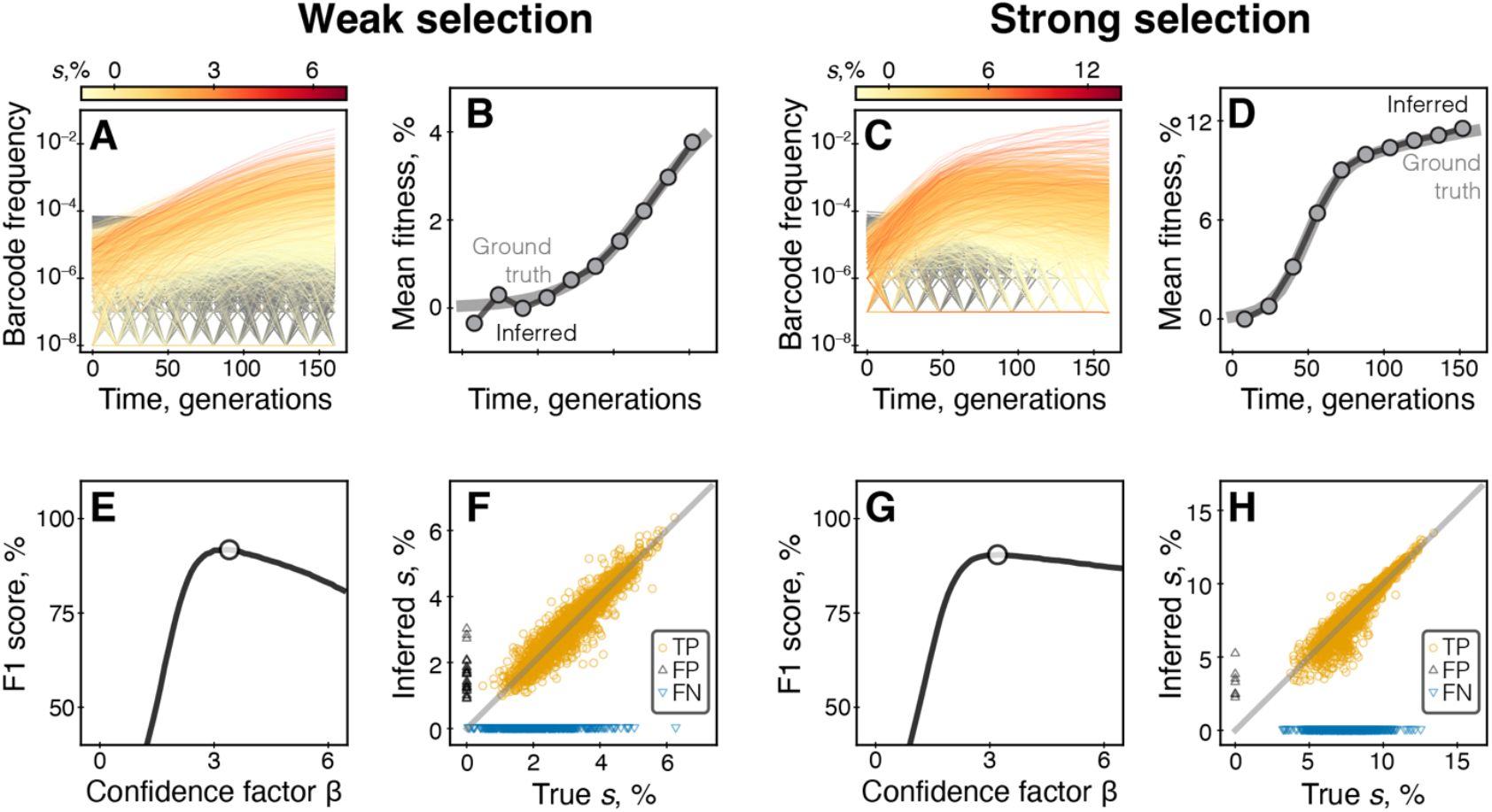
Performance of BASIL on simulated data. Left panels A, B, E, F show simulations in the weak selection regime; right panels C, D, G, H show simulations in the strong selection regime. **A, C.** Observed barcode frequency trajectories colored by fitness. **B, D**. Mean fitness trajectories. Grey lines show the ground truth, circles show values inferred by BASIL. **E, G**. F1 score as a function of the confidence factor β. White circle denotes the maximal value. **F, H**. Inferred versus true selection coefficients of individual lineages. Lineages are called adapted with β = 3.3 (see main text). Each symbol corresponds to a lineage; orange circles, black upward triangles and blue downward triangles represent true positives, false positives and false negatives, respectively; true negatives are not shown.

While the performance characteristics of our method are comparable or superior to those of the existing approaches, BASIL nevertheless fails to identify a significant proportion (between 15 and 20% in our simulations) of truly adapted lineages (Table 1). This is probably a biological inevitability rather than a flaw of the method itself. Some lineages, particularly those with weak adaptive mutations, may simply not reach large enough population sizes before they begin to decline due to clonal interference, which would prevent them from being identified as adapted by any method. Indeed, we find that false negative lineages in our simulations are typically only slightly more fit than the ancestor and their trajectories are essentially indistinguishable from those of truly neutral lineages, which supports our conjecture (see Section 4.2 in the Supplementary Information and Figures S6–S8 for more details).

### Comparison of methods on published BLT data

We next sought to compare the performance of BASIL against FitMut2 on data from published BLT experiments. In the absence of ground truth, we took two approaches to carry out this comparison. First, Venkataram et al sampled 410 yeast clones from their BLT experiments and estimated their fitness in two separate competition assays, carried out in the presence and absence of the alga *Chlamydomonas reinhardtii* [33]. Although the fitness of these clones may differ from the fitness of the corresponding adapted lineages in the original BLT experiments, these estimates represent the best available approximation of the ground truth. Thus, we asked how accurately BASIL and FitMut2 predict the fitness of these clones based on the frequency trajectories of the corresponding lineages in the BLT experiments. Second, we evaluated how the performance of FitMut2 and BASIL degrades as we subsample the Levy et al data [3], which is the largest and the “cleanest” BLT dataset available so far (see Data S1, Tab 3 and Figure 2E).

### Comparison with competition assay measurements in Venkataram 2023 data

We find that the Pearson correlation coefficient between the fitness of yeast clones measured in competition assays and the corresponding BLT fitness estimates produced by BASIL is 0.751 and 0.636 (Figure S9) for the experiments with and without the alga, respectively; and the average difference between the estimates is 2.56% and 3.52%. For FitMut2, the Pearson correlation coefficients in the two environments are 0.257 and 0.47 and the average difference between the estimates is almost three times larger than for BASIL, 6.02% and 10.34%, respectively.

### Down-sampling analysis on Levy 2015 data

To test how the amount of data affects BASIL and FitMut2 performance, we down-sampled the Levy 2015 dataset [3] in two ways (see Materials and Methods). First, we reduced average coverage from about 160× to about 10× per lineage. Second, we reduced the sampling frequency from 12 time points to 8. We then compared how various performance metrics for each method degrade relative to the full dataset.

The results of this analysis are shown in Figure 6. We find that when coverage is reduced, the mean fitness trajectories inferred by either method remain virtually unchanged (Figure 6A,D) but the number of lineages identified as adapted decreases. Specifically, BASIL identifies 49,330 lineages as adapted in the reduced data compared to 60,920 in the full data, a 19% decrease. In contrast, FitMut2 identifies only 1,131 lineages as adapted in the reduced data compared to 15,386 in the full data, a 92.6% decrease. For both methods, the overwhelming majority (≥ 98.8%) of lineages identified as adapted in the reduced data are also identified as such in the full data, and the estimates of selection coefficients remain largely unchanged (Pearson correlation coefficient of 95.8% and 82% for BASIL and FitMut2, respectively; and an average change of 0.20% and 0.83% in the estimated value of *s*, respectively; Figure 6B,E and Table S7). Notably, for BASIL, lineages that are no longer identified as adapted after down-sampling are those with small selection coefficients whereas FitMut2 loses adapted lineages even with large selection coefficients (Figures 6C,F and Table S7).

**Figure 6.**
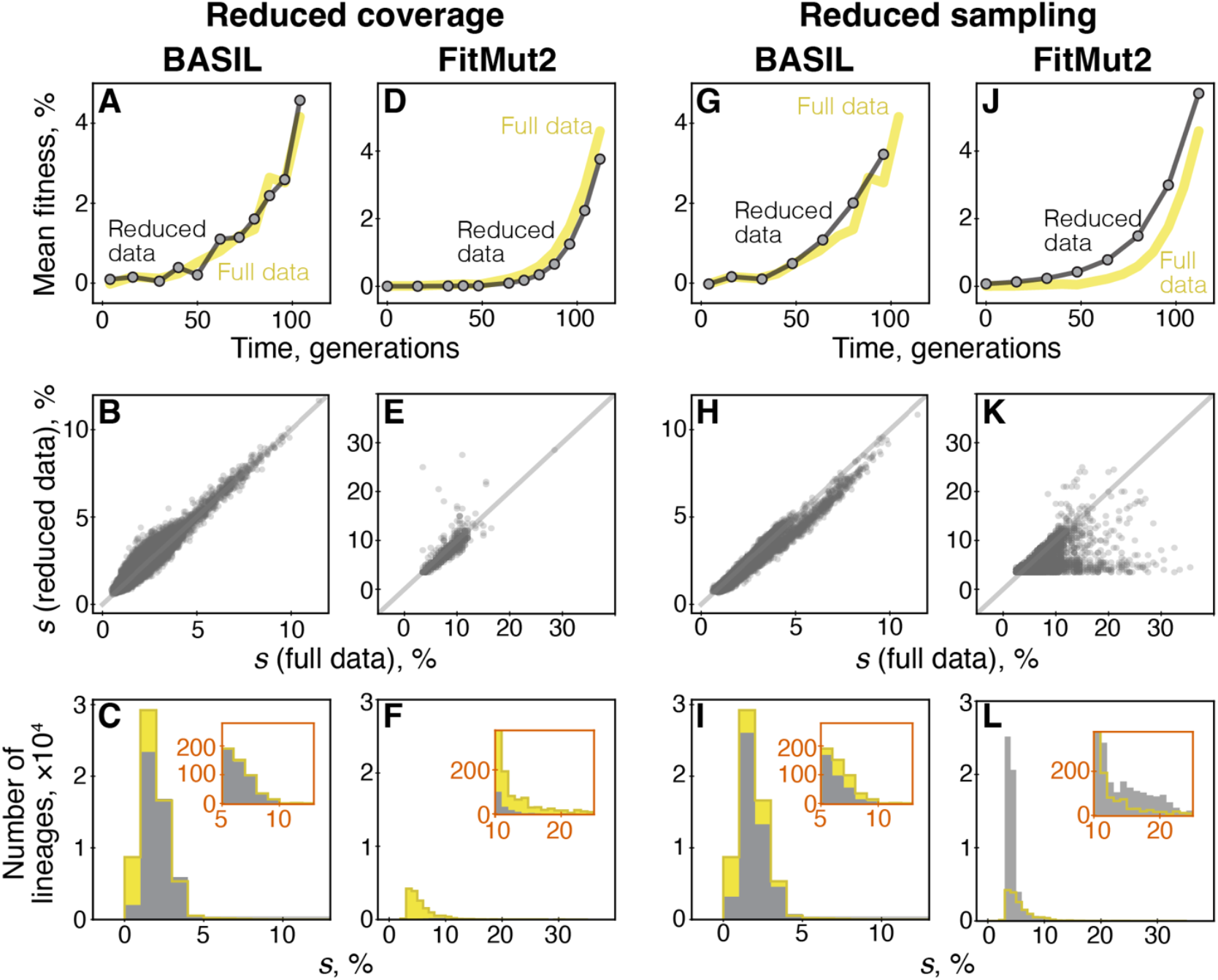
The effects of down-sampling on BASIL and FitMut2 performance. We reanalyzed the Levy 2015 dataset (Replicate 1) using BASIL and FitMut2 after reducing sequencing coverage or sampling frequency. See Materials and Methods for details. **A.** Mean fitness trajectories inferred by BASIL on full and reduced data. **B**. Inferred selection coefficients for lineages identified as adapted by BASIL in both full and reduced datasets. The 1-to-1 line is shown in light gray. **C**. Distribution of fitness effects of lineages identified as adapted by BASIL in full and reduced data. Inset shows the tail of the distribution. **D–F**. Same as A–C but for FitMut2. **G–L**. Same as A–F but for reduced sampling.

When we reduce sampling frequency, BASIL exhibits a similarly modest degradation in performance. The number of adapted lineages decreases by 21% (from 60,920 to 48,206), with 99.9% calls being the same as in the full dataset. Their inferred fitness effects also remain consistent (Pearson correlation coefficient of 97.9% and 0.18% change in *s*, Figure 6H and Table S7). One noteworthy difference compared to reduction of coverage is that, when sampling is reduced, BASIL loses power to identify lineages with both weak and strong adaptive mutations (compare panels C and I in Figures 6).

In contrast to BASIL, FitMut2 exhibits a dramatic degradation of performance under sampling frequency reduction. Pathologically, in addition to 15,386 lineages identified as adapted in the full data, FitMut2 finds 39,862 additional likely spurious adapted lineages in the reduced data, most of which have small selection coefficients (Figure 6L and Table S7). Moreover, for those lineages that are identified as adapted in both full and reduced data, FitMut2 estimates different selection coefficients (Pearson correlation coefficient of 62.0% and 1.29% change in *s*, Figure 6K and Table S7).

In summary, reduction in sampling frequency is more detrimental for inference than a uniform reduction in coverage across all time points for both methods. Yet, while both types of data reduction degrade BASIL’s performance only to moderate degree, FitMut2 exhibits a dramatic loss of power under coverage reduction and a troubling increase in the rate of false positives under sampling frequency reduction. Thus, taken together, these analyses demonstrate that BASIL is a more statistically robust, accurate and precise approach for the analysis of BLT data than FitMut2.

### Application of BASIL to data from BLT experiments

Finally, we applied BASIL to five published BLT experiments (including Levy et al [3] and Venkataram et al [33]) carried out with various strains of yeast *S. cerevisiae* propagated in different environmental conditions (see Data S1, Tab 3). The main goal of this analysis was to expose BASIL to a variety of evolutionary scenarios, examine its outputs and look for potential failure modes that we have not observed so far. To this end, we analyzed BLT experiments described in Refs. [3,4,29,32,33].

We find that the mean fitness inferred by BASIL is generally consistent across replicates and with a posteriori estimates calculated based on the inferred selection coefficients and frequencies of identified adapted lineages (Figure S10, Data S1, Tab 3; see Methods). However, in the Venkataram 2023 data, we find that the mean fitness inferred by BASIL begins to decline at later time points. Since there is a strong theoretical expectation that the mean fitness in large populations should not decline, this observation indicates that BASIL fails to accurately infer mean fitness at later time points in this dataset. We suspect that this failure is caused by the prevalence of secondary adaptive mutations, i.e., adaptive mutations arising in already adapted lineages [4,5]. Given that adaptation in this experiment appears to be extremely fast (0.28% per generation on average; Data S1, Tab 3), secondary adaptive mutations likely arise sooner than in other experiments. How secondary mutations can lead to declines in the inferred mean fitness can be rationalized as follows. As lineages acquire secondary adaptive mutations, their frequencies begin to rise faster than expected from previous observations. However, at later time points, BASIL has high confidence in the fitness estimates of most lineages. Thus, as it estimates the mean fitness at the next time interval, it attributes the unexpected frequency increases of multiple lineages to a decline in the mean fitness. We note that, while BASIL’s apparent inability to accommodate changes in belief distributions at later stages of the BLT time course is sub-optimal, it does not represent a major problem. Indeed, the main purpose of BLT experiments is to measure the fitness effects single-mutant neighbors of a given wildtype genotype and to isolate some of them for further investigation [3,29,32,33,37,38]. Thus, the focus on typical BLT experiments is on the early stages of adaptation before secondary mutations become prevalent (but see Refs. [4,5] that examine longer-term dynamics).

Next, we examine the fitness effects of individual adaptive mutations. We find that the number of identified adapted lineages can vary across replicate BLT experiments by up to threefold (Data S1, Tab 3). We also find that, while the measured distributions of fitness effects (mDFEs) look similar across replicate experiments (Figures 7, S11), they are statistically distinguishable by several measures (Table S6). These differences could arise from real biological variation, measurement noise, or both. As discussed above, variation in coverage and sampling frequency leads to variation in statistical power to detect beneficial mutations (see Section “Comparison of methods on published BLT data”). Our analysis showed that this variation in power is modest in the Levy 2015 data and would not be sufficient to fully explain the differences between the numbers of detected adaptive mutations across replicates. Biologically, strong rare beneficial mutations could arise in different replicates at different times, causing real biological variation in the mean fitness trajectories (Figure S10), which, even if small, could in turn lead to variation in the probability of establishment of weaker beneficial mutations. These observations highlight the difficulties of inferring the shape of the true underlying DFE from the fitness identified adapted lineages.

**Figure 7.**
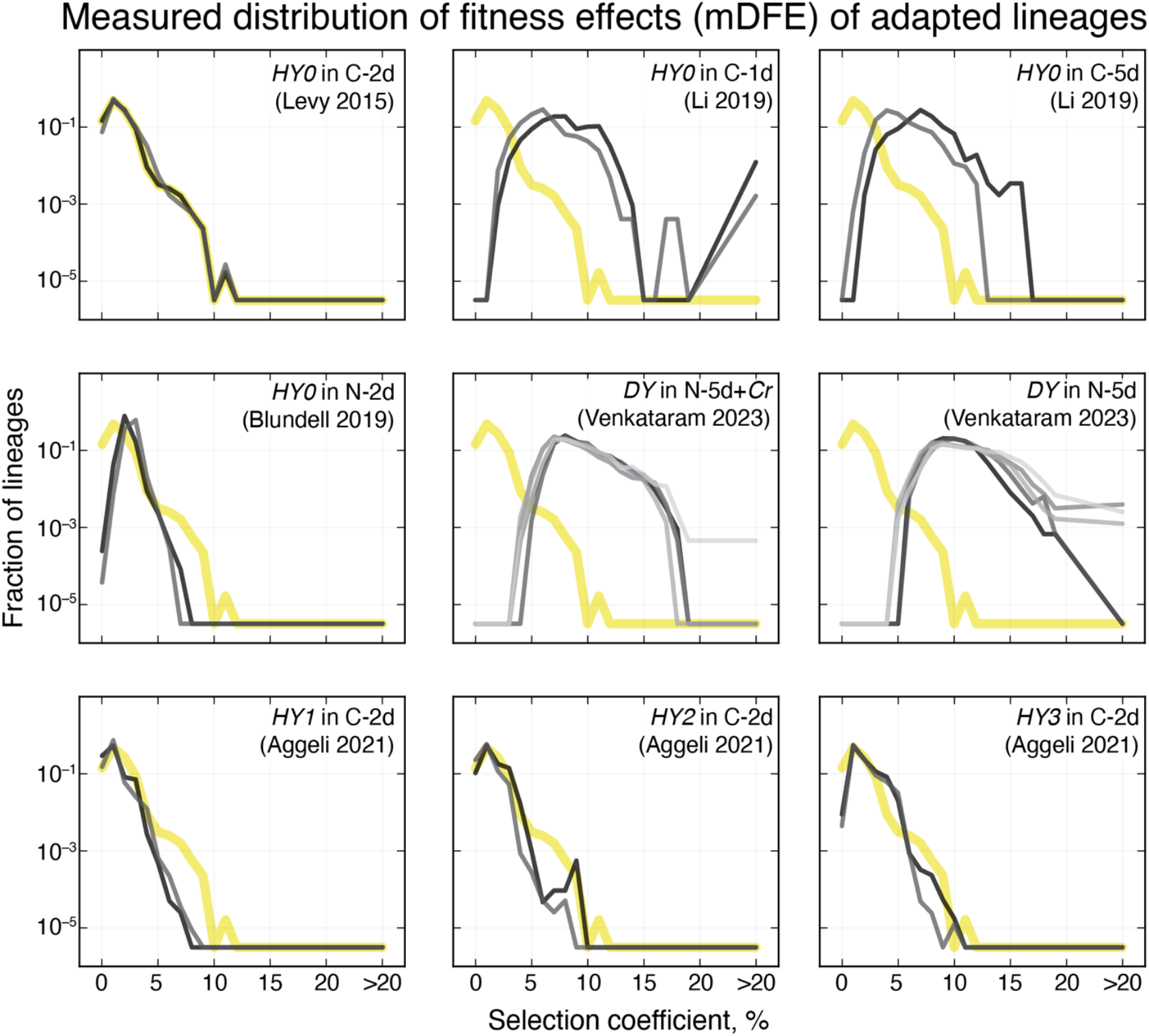
Histograms of the measured distributions of fitness effects (mDFEs) of adapted lineages inferred by BASIL in nine BLT experiments. Each panel shows a BLT experiment, as indicated. Gray lines are different replicates. In the top right corner, we provide a short-hand description of the strain (haploids are denoted HY*i*, diploids are denoted by DY) and environment used in each experiment. “C” and “N” refer to the likely limiting nutrient in the medium, -*x*d indicates the number days in the growth and dilution cycle, and “+*Cr*” indicates the presence of the alga *Chlamydomonas reinhardtii* in the evolution environment (see Data S1, Tab 3 for additional details). Thick yellow line shows the mDFE of *HY0* in C-2d (Levy 2015, Replicate 1) as reference.

## DISCUSSION

In this work, we showed that the existing approaches to the analysis of BLT data have significant shortcomings. Using simulated data with pre-existing mutations, we found that both the original Levy-Blundell (LB) method and its improved version implemented in the FitMut2 package, can produce biased estimates of the fitness effects of new mutations, particularly under strong selection. We provided the likely mechanistic explanation for the suboptimal performance of the LB method. Using data from the Levy et al BLT experiment [3], we showed that FitMut2 can both exhibit significant loss of statistical power and elevated rates of false positives when the amounts of available data are reduced.

To overcome these shortcomings of existing approaches, we developed BASIL, a Bayesian filtering method for identifying barcoded lineages that carry nascent adaptive mutations and for inferring their selection coefficients. BASIL estimates belief distributions for these selection coefficients and the corresponding lineage sizes, updating them over time based on the observed barcode read counts. Measurement noise is a key feature of our model, and our new experimental data suggests that the measurement variance in the read count scales quadratically with the expected read count, as opposed to a linear scaling that has been postulated in previous studies. This scaling (with *ϵ* being a free parameter) is implemented in BASIL, and we suggest to use this scaling in future barcode-analysis models. One notable advantage of our framework is that it estimates mean fitness based on reference lineages, without the need to be designated neutral reference lineages a priori. This makes our approach applicable to a wide range of evolutionary scenarios, including those with strong selection.

When we applied BASIL to real BLT data, we found that, in contrast to FitMut2, BASIL exhibits only a moderate loss of power when the sequencing coverage or sampling frequency are reduced, and its estimates of selection coefficients of adaptive mutations remain consistent.

BASIL computes a time-dependent joint belief distribution for the selection coefficient and size of every lineage. So far, we did not utilize all the information contained in this distribution, but instead chose to classify lineages into adapted and neutral with a simple linear classifier that is based on a snap-shot of this distribution. In our simulations, this simple classifier with the confidence factor of 3.3 (which controls the slope of the classification line) was very effective. How effective it is on real data, which is arguably more complex than our simulations, is difficult to ascertain. We visualize our classification on two-dimensional diagnostic plots where the mean of the fitness belief distribution is plotted against its standard deviation (Figures S5, S12). In some datasets, these plots reveal clustering of lineages, with some clusters clearly containing highly adapted lineages (Figure S12), similar to our simulations (Figure S5). It is possible to isolate these highly adapted clusters by adjusting the confidence factor, which can be easily done by the user in the current implementation of BASIL. However, while such an adjustment would likely improve precision, it may significantly worsen recall, since not all adapted lineages belong to the highly adapted clusters, as we saw in our simulations (Figure S5). Instead, developing more sophisticated classifiers and testing them on more realistic simulations could be a valuable direction for future research.

Our approach has certain limitations. On the technical side, BASIL is quite computationally expensive since it requires extensive Markov Chain Monte Carlo (MCMC) sampling to estimate posterior distributions of lineage fitness and sizes. For instance, the analysis of a BLT experiment with 5×10^5^ lineages sampled at 10 time points took us ∼50 hours of running time on AMD Ryzen 5 7600X 6-Core Processor, and the running time increases linearly in the number of lineages and sampling time points. However, since MCMC computations for different lineages are independent, the algorithm is easily parallelizable. In our analyses, we found it convenient to use 12–32 processor cores.

One important conceptual limitation of our model is that it assigns a single constant selection coefficient to each barcoded lineage, but many barcoded sub-populations are in fact polymorphic. Indeed, each new adaptive mutation arises in a single cell, and the mutant lineage initially constitutes only a small fraction of the parental barcoded sub-population [3]. The mutant lineage outcompetes the wildtype within this sub-population gradually, such that the barcode frequency increases slower than would be expected from the mutant’s selection coefficient alone. Since BASIL ignores these intra-lineages dynamics, it probably underestimates the selection coefficient of lineages that arise later in the BLT experiment and also undercounts them. However, the magnitude of this bias is hard to predict a priori because it depends on the rate of adaptive mutations and their fitness effects, the very quantities that we are trying to estimate.

A barcoded sub-population also becomes polymorphic when secondary adaptive mutations arise and begin to establish [5]. In principle, BASIL is capable of updating the belief distribution for any given lineage if its fitness changes, but in practice, convergence to a stable new distribution is probably slow at later time points when the confidence in the current belief is already high. As a result, when secondary mutations become dominant, BASIL can fail. While this represents a limitation, it is not a major one since the purpose of typical BLT experiments is to estimate the selection coefficients of single adaptive mutations rather than their combinations. Thus, when secondary beneficial mutations become common, BLT experiments become much less informative and are typically aborted. But given this limitation, BASIL is currently not expected to be reliable in BLT experiments under fluctuating or frequency-dependent selection or in spatially structured populations where expansion of adapted lineages is sub-exponential [55]. However, it should be possible to extend our approach to these more general cases.

It is important to keep in mind that BASIL only identifies adapted lineages and infers their selection coefficients but does not infer the rate at which beneficial mutations arise. To infer the latter, we would need to know not only how many beneficial mutations were detected but also how many arose but were lost by drift while rare and how many were established but not detected. In other words, we would need a model of mutant’s evolutionary dynamics as well as a model of sampling and detection. Previously, Levy et al developed the former model and inferred the rates of adaptive mutations with different fitness effects [3], but their model did not account for biases introduced by sampling and detection. Our results suggest that these biases may not be negligible (Figures 6 and 7 and Table S6). Therefore, incorporating them into a model of beneficial mutation rate inference is an important problem for future research.

## MATERIALS AND METHODS

### Code and data availability

#### Code

All analysis code is written in Python and is available at https://github.com/Huanyu-Kuo/BASIL-public. Computationally intensive analyses were run on the Triton Supercomputing Cluster (TSCC).

#### Data

Barcode sequencing data generated in this work are available at https://data-view.ncbi.nlm.nih.gov/object/PRJNA1227420.

### Estimation of noise in the barcode frequency measurement process

#### Strains

26 clones were isolated from the barcoded library of the diploid strain GSY6699 of yeast *Saccharomyces cerevisiae* used in our previous work [33]. Details of the strain and library construction are provided in Refs. [3,31]. Briefly, each clone carries a unique 30-bp DNA barcode that replaces one copy of the YBR209W locus; all clones are otherwise identical in the rest of the genome. Barcode sequences of all clones have been determined previously [33] and are reported in Data S1, Tab2.

#### Culture preparation

To probe the measurement noise process, we aimed to construct a barcoded clone mixture consisting of 5 clones at each of five frequencies 0.1, 0.01, 10^−3^, 10^−4^, 10^−5^, and one clone at the remaining frequency of approximately 0.45 (26 clones total). We started a liquid culture of each clone from frozen stocks in 3 ml of YPD (10 g yeast extract, 20 g peptone, 20 g dextrose, 1 L Milli-Q water) in 16-mm test tubes, and incubated these cultures overnight, in a shaking incubator set at 200 rpm at 30°C. After overnight growth, culture densities were measured with a Coulter Counter and were found to be around 1.5×10^8^ cell/ml. The measured values for all cultures are provided in Data S1, Tab2. For clones with the four lowest target frequencies (those below and including 0.01), we diluted the overnight cultures 1:10, 1:100, 1:10^3^ and 1:10^4^ in PBS, respectively. To make the final clone mixture, we pooled 200 μl of each diluted culture, then added 200 μl of the five cultures with target frequency 0.1, and then added 600 μl of the culture with the largest frequency. This resulted in final mixture volume of 5.6 ml at density approximately 4.6×10^7^ cell/ml with barcodes clustered around six distinct frequencies: 0.395, 0.108, 0.019, 1.24×10^−3^, 1.28×10^−4^ and 1.41×10^−5^. Our estimates of clone frequencies in the mixture are provided in Data S1, Tab2. We spun down the clone mixture, re-suspended it in 20 ml of PBS (final density 1.57×10^7^ cell/ml, split it into 20 aliquots (1 ml each in a 2 ml cryogenic tube), supplemented with 500 μl of 80% glycerol, and stored at –70°C.

#### Barcode sequencing

We followed the same protocols for DNA extraction and sequencing library preparation as in our previous work [33]. We repeated this procedure 10 times, each time using one aliquot of the barcoded clone mixture stocks described above. All ten replicate libraries were sequenced on the Illumina HiSeq platform, resulting in approximately 2.3×10^−6^ total reads.

#### Data analysis

We used BarcodeCounter2 to identify barcodes in the sequencing data and count them [56]. The data are shown Data S1, Tab2. One replicate was excluded from analysis due to low coverage. In addition, we could not identify barcodes for two out of 26 clones, and two other clones (in the lowest frequency class) were not detected in the sequencing data. These four barcodes were removed from further analysis. Thus, we retained 22 barcodes present in all *n*_rep_ = 9 replicates.

Denote by *X*_*p*_ the set of barcodes *b* with the same target frequency *p* and denote the read count for barcode *b* in replicate *i* by *r*_*bi*_. For each frequency class *X*_*p*_ and replicate *i*, we estimate the variance in the read count due to measurement noise as 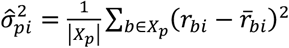, where 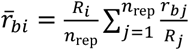 is the estimate of the expected read count for barcode *b* in replicate *i, R*_*i*_ is the total coverage in replicate *i* and |*X*_*p*_| denotes the total number of barcodes in the frequency class *p*. More details are provided in Section 1 in the Supplementary Information.

We model the relationship between read count mean 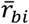 and variance 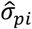 shown in Figure 4B as 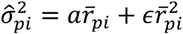 and test four special cases of this equation. To recapitulate the Poisson distribution, we set *a* = 1 and *ϵ* = 0. We refer to the noise model proposed by Levy et al (see equation S8) as “Linear general”. In this model, *ϵ* = 0 and *a* is a free parameter. To capture the possibility that the degree of overdispersion varies across read depths, we test a quadratic relationship with *a* = 1 and a fitting parameter *ϵ* > 0, which matches the Poisson model for small 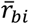. This is the model in equation (2), and we refer to it as “Quadratic with 1-parameter”. Finally, we also fit the general quadratic model with two fitting parameters *a* and *ϵ*.

We find the best-fit parameters using the non-linear least squares method. Since the estimated measurement noise variances span six orders of magnitude (see Figure 4B), we carry out the fitting in the log space, i.e., we minimize the squared error between 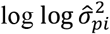 and 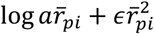. In Table S1 we report the best-fit parameters, residual sums of squares (RSS), as well as the coefficient of determination *R*^2^, computed as 1–RSS/TSS, where TSS denotes the total sum of squares. We compare the goodness-of-fit for all pairs of nested models using the *F*-test. Specifically, assuming that the null model has *p*_null_ free parameters and the alternative model has *p*_alt_ parameters *p*_null_ < *p*_alt_, we calculate the *F*-statistic as 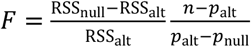, where *n* denotes the number of data points, and RSS_null_ and RSS_alt_ are the residual sums of squares for the null and alternative model, respectively. If the data is generated by the null model, then the *F*-statistic is distributed according to the *F*-distribution with (*p*_alt_ – *p*_null_, *n* – *p*_alt_) degrees of freedom.

The results of this analysis are shown in Figure 4B and Tables S1 and S2. We find that the quadratic models explain at least 93% of variance in our data compared to linear models, which explain 89% or less. We also find that the general quadratic model provides only a marginally better fit than the 1-parameter quadratic model, with a *P*-value of only 0.0235. Therefore, we use the 1-parameter quadratic model as our model for the relationship between read count mean and variance.

### BLT simulations

We carry our simulations in the weak and strong selection regimes (Figure S1). We chose the parameters for the weak selection regime to make it similar to the experimental conditions described by Levy et al [3]. In particular, selection and genetic drift are relatively weak, and the variance in the read count is a linear function of the mean. For the strong selection regime, we chose parameters that should make inference harder. In particular, selection and drift are stronger, and the variance in the read count is a quadratic function of the mean (see equation (2)). Simulation parameters for both regimes are provided in Table S3.

All simulations start with the same number *N*_*L*_ = 10^5^ of barcoded lineages. The initial size of each lineage is drawn from an exponential distribution with mean ⟨*N*⟩/*N*_*L*_, where ⟨*N*⟩ is the expected total population size (see Table S3). In each simulation, 3,000 randomly chosen lineages are adapted, such that all members of such lineages have the same beneficial mutation. We draw these lineages uniformly with the constraint that their initial size must exceed 1.5*D* = 384 individuals, where *D* = 256 is the dilution factor. The selection coefficient (fitness) *s*_*i*_ of each adapted lineage is drawn from the normal distribution with mean ⟨*s*⟩ and standard deviation σ_*s*_, and negative draws are discarded. The fitness of all other lineages is set to *s* = 0. No new mutations arise during the simulation.

After initializing our populations, we simulate their growth and dilution, with each growth-dilution cycle consisting of the following steps:

1. Each cycle begins with the calculation of the population’s mean fitness 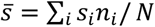, where *n*_*i*_ is the current size of lineage *i* and *N* = ∑_*i*_ *n*_*i*_ is the current population size.
2. We then simulate the dilution process. For each lineage *i*, we draw its new size 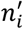 as a random number from the Poisson distribution with mean *n*_*i*_/*D*. Thus, after dilution, the population’s bottleneck size equals 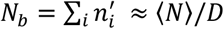.
3. We then simulate population expansion. Each lineage *i* expands by the factor 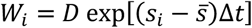, where Δ*t* = 8 generations is the duration of the growth phase. Thus, at the end of the growth phase, the new lineage size is 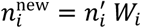 (rounded to the nearest integer) and the total population size is again 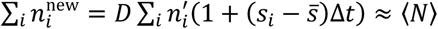.

Each simulation lasts for a total 20 growth-dilution cycles (160 generations). We simulate sampling and barcode sequencing by selecting *R* random individuals from the population at the end of every other cycle (i.e., before dilution). For each sampled barcode *i*, the number of sequencing reads *r*_*i*_ is drawn from the negative binomial distribution with mean ⟨ *r*_*i*_ ⟩ = *n*_*i*_ *R*/*N* and variance 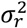, where *R* is the sequencing depth (see Table S3). The output of each simulation is a table of “sequenced” barcode counts at all sampling time points.

### The neutral decline method

The neutral decline method retains the essential features of the original LB method but simplifies it in two major ways. First, in the LB model, adapted lineages are characterized by two parameters, the selection coefficient *s* and the establishment time τ, whereas the neutral decline method has only one parameter *s* per lineage. However, in our simulations, all adapted mutations are pre-existing, implying that our simpler one-parameter model is sufficient. The second important difference is that, in addition to the mean fitness 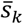, the LB method also estimates the noise parameter κ (see Section 2 in the Supplementary Information). To infer both 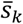 and κ, the LB method uses a model that describes how the entire distribution of read counts changes over time (equations (S8), (S9)). The mean of this distribution is described by equation (1), which does not depend on κ. Thus, equation (1) is sufficient if we are interested only in 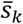 (see Figure S2). Importantly, since equation (1) directly follows from the LB model, a failure of this equation would imply a failure of the full LB model. Therefore, the focus our analysis on the neutral decline rather than the full LB model is justified.

The neutral decline method proceeds as follows.

1. For each consecutive time interval (*t*_*k*–1_, *t*_*k*_), we choose the neutral reference lineages as those having certain read counts *r*_*k*–1_ at the beginning of the interval with either 20 ≤ *r*_*k*−1_ ≤ 40 (lowabundance reference) or 80 ≤ *r*_*k*−1_ ≤ 100 (high-abundance reference).
2. For each initial read count *r*_*k*–1_, we calculate the mean fitness 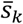 using equation (1) and over all initial read counts (for reference lineages only). We repeat the mean fitness calculation for all sampling intervals and thereby obtain the full mean-fitness trajectory 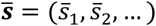.
3. To infer the fitness effect *s*_*i*_ of lineage *i*, we maximize the log-likelihood function of the lineage trajectory 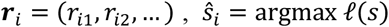, where 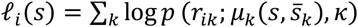 with *p*(*r*: *μ, κ*) given by equation (S8), and 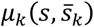 given by equation (S9) and *κ* = 3.
4. To determine whether lineage *i* is adapted, we apply the likelihood ratio test to compare the neutral model (*s*_*i*_ = 0, null hypothesis) and the adaptive model (*s*_*i*_ = ŝ_*i*_), alternative hypothesis). The likelihood-ratio test statistic is given by *R*_*i*_ = ™2(*ℓ*_*i*_ (0) − ℓ_*i*_ (*ŝ*,)). We call the lineage adapted if *ŝ*, > 0 and *R*_*i*_ ≥ 3.84, which is the 95th percentile of the χ^2^-distribution with 1 degree of freedom.

### Empirical test of the neutral decline equation

In Figure 2, to test the neutral decline equation (1), we use the barcode read count data from two BLT simulations and the three BLT experiments Levy 2015 (replicate 1 in Ref. [3]), Li 2019 (replicate 2 from the 1-day transfer cycle experiment from Ref. [29]), and Venkataram 2023 (replicate 5 of the experiment with the alga from Ref. [33]). For each real or simulated BLT experiment, we selected one “early” and one “late” sampling time interval (*t*_*k*–1_, *t*_*k*_), as described in the caption to Figure 2. For each interval, we calculate the average read count 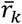 and the rescaled average read count 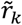 for lineages that had the same read count *r*_*k*−1_ ∈ [0,100]. If the number of lineages with a given *r*_*k*–1_ was less than 5, we did not estimate the corresponding 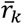. To investigate whether 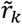 is a linear function of *r*_*k*–1_ as predicted by equation (1) for neutral lineages, we plot 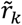 against *r*_*k*−1_. For some *r*_*k*–1_ in some datasets, this dependence becomes very noisy or clearly deviates from linearity. Thus, we first visually identify the intervals of *r*_*k*–1_ where the data behaves approximately linearly. We choose the interval 5 ≤ *r*_*k*−1_ ≤ 15 for the Li 2019, Venkataram 2023 and the strong-selection simulated datasets at the late time point; we choose the interval 5 ≤ *r*_*k*−1_ ≤ 40 for the Venkataram 2023 dataset at the early time point; and we choose 5 ≤ *r*_*k*−1_ ≤ 99 for all the other datasets. Then, using the data only within these intervals, we fit a linear model 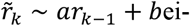- ther with *b* = 0 (restricted model) or with *b* being a free parameter (full model) and ask whether the full model provides a statistically better fit to our data using the *F*-test (see Section “Estimation of noise in the barcode frequency measurement process”).

### BASIL

The detailed description of BASIL is given in Section 3 of the Supplementary Information. Here, we provide the key expressions and brief descriptions of the algorithm.

#### Model of measurement

We model the probability of observing *r* reads for a lineage with *n* cells in a total population of size *N* which was sequenced to depth *R* as

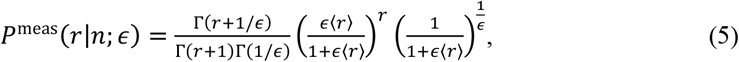

where *r*⟩ = *nR*/*N* is the expected number of reads.

#### The belief distribution for lineage size and fitness

Our main goal is to estimate the belief probability distributions 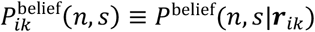, where ***r***_*ik*_ = (*r*_*i*0 =_, …, *r*_*ik*_) is the read count vector for lineage *i* up to and including sampling time point *t*_*k*_. We express these belief distributions as *P*^belief^(*n, s*|***r***) = *P*^belief^(*n*|*s*, ***r***) *P*^belief^(*s*|***r***) with parametric forms *P*^belief^(*n*|*s*, ***r***) = *γ*(*n*: *ϰ*(***r***), *θ*(*s*, ***r***)) and *P*^belief^(*s*|***r***) = *N*(*s*: *μ*(***r***), *σ*^2^ (***r***)), where *N*(*s*: *μ, σ*^2^) is the normal distribution with mean µ and variance σ^2^ and *γ*(*n*: *ϰ, θ*) is the gamma distribution with the shape parameter *κ* and scale parameter θ and we use the family of functions

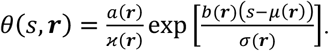

Thus, our belief distributions come from a family with five parameters µ, σ^2^, *x*, *a* and *b* all of which are fit based on the data vector ***r***.

At *t*_0_, we set µ = 0, σ^2^ = 0.1 and *b* = 0 for all lineages; and for lineage *i*, we set *κ*(*r*_*i*0_) = *r*_*i*0_/(1+0.01*r*_*i*0_), *a = r*_*i*0_*N*/*R*_0_ where *R*_0_ is the coverage at the initial time point. We believe that the prior distribution should be fairly broad (i.e., uninformative), but this particular choice is likely unimportant.

#### Projecting the belief distribution

To obtain the prior distribution 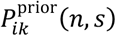 for the sampling time *t*_*k*_, we project the belief distribution 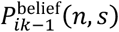 as follows. First, we obtain the 1-cycle pro-jected probability distribution

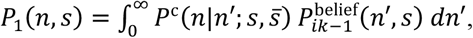

where 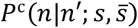 is the one-cycle transition probability given by equation (S34). The marginal distribution for *s* remains unchanged during projection. The probability that the lineage goes extinct during one cycle is 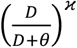 where *D* is the dilution factor and *κ* and θ are the parameters of the belief distribution 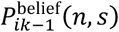. Conditional on lineage survival, we approximate the distribution *P*_*_ (*n*|*s*, ***r***) by the gamma distribution with shape parameter *κ*_1_ and scale parameter θ_1_, which can be calculated as functions of *s*, *κ* and θ (equations (S42) and (S43)). If multiple cycles elapse between successive sampling time points, the prior probability 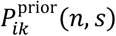 is obtained by applying this procedure recursively.

#### Updating the belief distribution

The belief distribution is updated using Bayes’ theorem (equation (3)). We would like to use the same analytical parametric as above for the updated belief distribution, but since the numerator of this equation is a complex function of both *n* and *s*, the normalization constant 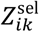 and the moments of the right-hand side of equation (3) cannot be computed analytically. Therefore, to estimate the parameters of the updated belief distribution, we use the Markov Chain Monte Carlo (MCMC) approach described in Section 3.2.2 in the Supplementary Information.

#### Calculation of 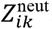

Equation (4) for the estimation of mean fitness 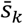 and the noise parameter *ϵ*_*k*_ in the time interval (*t*_*k*−*1*_, *t*_*k*_) depends on the marginal likelihood 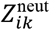 that lineage *i* which was classified as putatively neutral in this interval has *r*_*ik*_ reads at time *t*_*k*_. We obtain 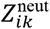 as follows. First, we approximate the marginal belief distribution 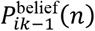 for lineage size *n* at *t*_*k*−1_ by a gamma distribution with shape parameter 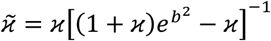 and scale parameter 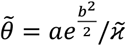, where *κ*, *a* and *b* are the parameters of the original distribution 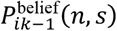. We then obtain the prior distribution 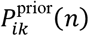 under the neutral model analogously to how we obtain 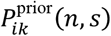 in the model with selection. Finally, we use MCMC to estimate 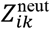 (see Section 3.3 in the Supplementary Information for details).

### Performance of BLT analysis methods on experimental data

#### Down-sampling

To test how sequencing coverage and sampling effort affect our mDFE inference, we down-sampled the data from Replicate 1 of the original BLT experiment with strain HY0 in the C-2d environment [3]. The original data has the average coverage of ∼160× per lineage, and sampling occurred at 12 time points, at generations 0, 16, 32, 40, 48, 64, 72, 80, 88, 96, 104 and 112. To reduce coverage, for each barcode whose read count is *r*_*ik*_ in the original data, we draw a random number 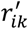 from a Poisson distribution with mean *r*_*ik*_/16, such that the average coverage per lineage is ∼10. To reduce sampling frequency, we remove sampling time points at generations 40, 72, 88 and 104, such that the total number of samples is reduced from 12 to 8.

#### Mean fitness based on inferred adapted lineages

Since the true mean fitness in the BLT experiments is unknown, we follow Levy et al [3] and validate the mean-fitness estimates obtained by BASIL by comparing them to a posteriori mean-fitness estimates based on the inferred selection coefficients and frequencies of identified adapted lineages 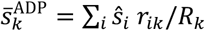 where *ŝ*, is the inferred selection coefficient of adapted lineage *i*.

#### Comparison of replicate mDFEs

BASIL estimates of the selection coefficients of adapted lineages for all datasets shown in Figure 7 are provided in Data S1, Tabs 4–12. To construct the measured distribution of fitness effects of new mutations (mDFE) shown in Figure 7, we group lineages based on their inferred selection coefficient *ŝ*, into 1-percent bins as well as a bin of all lineages with *ŝ*, > 0.2. The statistics of these mDFEs are reported in Data S1, Tab 3.

We test the consistency of mDFEs across replicates of the same BLT experiment using quantile-quantile (Q-Q) plots (Figure S11). We find that, by this metric, most mDFEs inferred from replicate experiments are consistent with each other, although experiments HY0-C1d [29], HY2-C2d, HY3-C2d [32] and DY-N5d [33] show some discrepancies. In addition, we compare the mean, the median, the interquartile range (IQR), and the interdecile range (IDR) across replicate mDFEs using a permutation test. To this end, we create a list of selection coefficients of all lineages identified as adapted in any of the replicates and then randomly reassign each selection coefficient to each replicate *i* while preserving the original number of selection coefficients in each replicate. We estimate each mDFE statistic of interest for each permuted replicate. Then, for each pair of replicates it in the permuted data, we calculate the absolute value of the difference between the statistic values; and we average them over all pairs to obtain our test statistic. We repeat this permutation procedure 5,000 times to obtain the null distribution for each mDFE statistic and calculate the empirical *P*-value based on this distribution. The results of this more sensitive test are reported in Table S6.

## ACKNOWLEDGEMENTS

We thank members of the Kryazhimskiy lab for discussions and feedback and anonymous reviewers for constructive feedback on the initial version of the manuscript. The UC San Diego Triton Computing Cluster assisted with computational work. This work was supported by the Career Award at Scientific Interface (1010719.01) from the Burroughs Wellcome Fund to SK, the Alfred P. Sloan Research Fellowship (FG-2017-9227) to SK, the Hellman Fellowship to SK, and the NIH grants 1R01GM137112 and R35GM153242 to SK.

## AUTHOR CONTRIBUTIONS

Conceptualization: HK, SK; Experimental design: HK, SK; Experiments: HK; Analysis: HK, SK; Writing: HK, SK.

## SUPPLEMENTARY MATERIALS

### Supplementary Data

Data S1 [Excel]. Aggregated Data Tables.

Tab 1. Contents.

Tab 2. Measurement noise experiment.

Tab 3. Characteristics of published BLT studies and some summary statistics of the BASIL analysis.

Tabs 4–12. BASIL results for published BLT experiments.

## Raw Data Repository

https://dataview.ncbi.nlm.nih.gov/object/PRJNA1227420

## Code

https://github.com/HuanyuKuo/BASIL-public

## Supplementary Materials

### 1 Estimation of measurement noise variance in the calibration experiment

Denote by *X*_*p*_ the set of barcodes with the same target frequency *p*. Variation in the number of reads *r*_*bi*_ across barcodes *b* ∈ *X*_*p*_ and replicates *i* arises from three sources. First, the same barcode is represented by different numbers of reads in different replicates since replicates receive different total coverage. Specifically, if the barcode’s true frequency in the pool is *p*_*b*_, we expect to obtain *p*_*b*_*R*_*i*_ reads for this barcode in replicate *i* with total coverage *R*_*i*_. Second, the actual number of reads obtained for barcode *b* will deviate from *p*_*b*_*R*_*i*_ by some random amount *δr*_*bi*_ due to noise during library preparation and sequencing [2]. We refer to *δr*_*bi*_ as “measurement noise”. Finally, while we aimed to seed the barcoded clones within the same frequency class *X*_*p*_ with exactly the same target frequency *p*, in reality, the frequency *p*_*b*_ of barcode in the pool actually deviates from *p* by some amount *δp*_*b*_, i.e., *p*_*b*_ = *p* + *δp*_*b*_. We refer to *δp*_*b*_ as the “experimental noise”. Putting all of this together, we can model the read count *r*_*bi*_ for barcode *b* ∈ *X*_*p*_ as

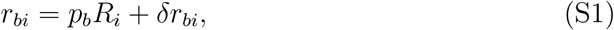

where *p*_*b*_ is unknown and we assume that all *δr*_*bi*_ are independent random variables with zero mean, and all *δr*_*bi*_ with the same *b* are identically distributed with variance 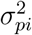(the noise variance) where index *p* indicates that noise variance can vary between frequency classes *X*_*p*_, and index *i* indicates that it can also vary between replicates *i*. Our goal is to estimate 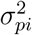.

The challenge with estimating 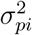 is that each barcode has an unknown expected number of reads *p*_*b*_*R*_*i*_ in each replicate *i*. If all barcodes had exactly the same frequency *p*, then we could use *r*_*bi*_ from different barcodes in the same replicate to estimate *p*_*b*_*R*_*i*_ as

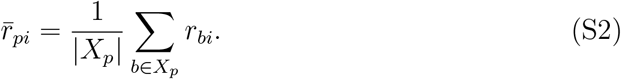

However, different barcodes *b* ∈ *X*_*p*_ in fact have different frequencies *p*_*b*_, and the estimate (S2), which ignores this variation, would provide a poor estimate for the expected barcode count *p*_*b*_*R*_*i*_ and risks inflating our estimate of the measurement noise. To overcome this problem, we first estimate *p*_*b*_ as

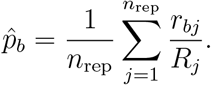

Then, we estimate the expected number of reads 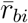 for barcode *b* in replicate *i* as

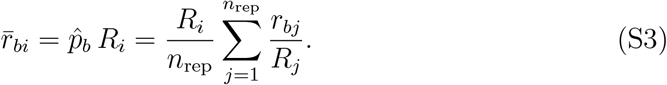

We can now use expectation (S3) to estimate the variance in the read count due to measurement noise in frequency class *X*_*p*_ and replicate *i* as

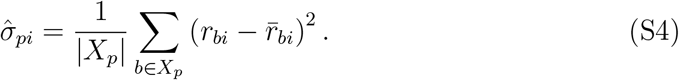

In Appendix A, we derive the expectation for this estimator. We find that this estimator is biased, approximately by a factor 1 − 1*/n*_rep_ and derive a bias-corrected estimate (see equation (S73)). However, this corrected estimate is not guaranteed to be non-negative, and indeed we found that the variance estimate for one of the replicates in the lowest frequency class becomes negative upon applying this correction. We also found that for 36 out of 41 (88%) values of 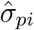 that we could estimate, the correction changes the estimated value by less than 25%. For these reasons, we decided to use the uncorrected estimate given by equation (S4) for the subsequent analysis.

### 2 Existing methods for the analysis of BLT data

Levy et al developed the first computational method for the analysis of barcode lineage tracking (BLT) data [3], which we refer to as the “Levy-Blundell” method, or the “LB method” for short. More recently, Li et al developed its extension, called FitMut2, and showed that it is superior to the LB method [4]. In this section, we briefly review both the original LB method and FitMut2.

#### 2.1 Review of the Levy-Blundell method

The LB method is based on a stochastic birth-death process that describes a mutant lineage starting from *n*_0_ founding cells that randomly divide and die with certain per capita birth rate *b* and death rate *d*. Levy et al derived the full stochastic expression for the probability *P*_*t*_(*n*) that the number of cells in the mutant lineage after time *t* is *n* (see equation (126) in the Supplementary Information to Ref. [3]). They then approximate this discrete distribution for large *n* by a continuous probability distribution density

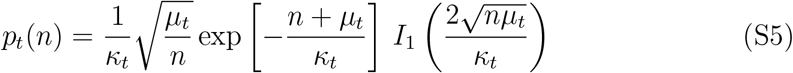

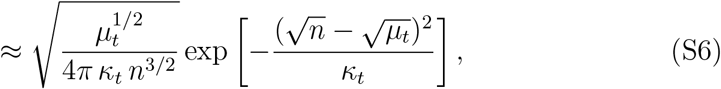

where *I*_1_(*x*) denotes the modified Bessel function of the first kind. They then use the approximated expression for the analysis of their data (see equation (127) in the Supplementary Information to Ref. [3])). equation (S6) depends on two parameters: the mean of lineage size *µ*_*t*_ = *n*_0_*e*^*λt*^ and the variance to mean ratio 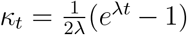, where *λ* = *b − d* is the mutant growth rate. One important difference between the full solution of the birth-death process and the approximation (S6) is that the full probability distribution has a non-zero weight at *n* = 0 (i.e., extinction is possible) whereas in the approximation (S6) the probability of extinction is zero.

The LB method then uses the same general form of equation (S6) to describe the dynamics of barcode read counts, that is, if a lineage with selection coefficient *s* is observed to have *r*_*k*−1_ read counts at the previous cycle *k* − 1, then the probability density *p*(*r*) that *r* counts will be observed at the next cycle is assumed to be

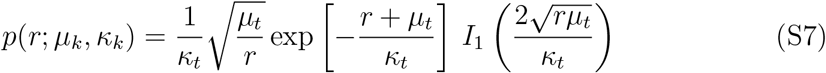

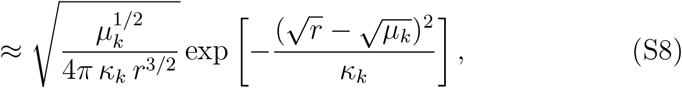

where

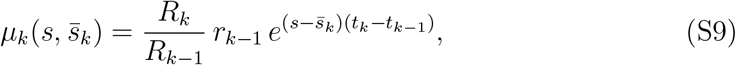

is the expected read count. Here *R*_*k*−1_ and *R*_*k*_ are the total read depths at sampling times *t*_*k*−1_ and *t*_*k*_ respectively and 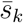 is the mean fitness of the population in the interval (*t*_*k*−1_, *t*_*k*_). Unlike in the birth-death process, the variance parameter *κ*_*k*_, which now characterizes all the noise in the BLT experiments including growth, dilution, DNA extraction, DNA amplification, etc., is unknown a priori. To determine *κ*_*k*_, Levy et al performed additional experiments and estimated *κ*_*k*_ ≈ 3 (see equations (40, 45) in the Supplementary Information to Ref. [3]). However, *κ*_*k*_ is treated as a fitting parameter when BLT data are analyzed, as described below.

Levy et al apply the following procedure to infer the lineage selection coefficients *s* from the BLT data. At each cycle *k*, they first infer the mean fitness 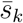 using lowabundance lineages, i.e. those with read counts 20 ≤ *r*_*k*−1_ ≤ 40. The assumption is that these lineages are so small in size that they have not acquired any adaptive mutations, and so the selection coefficient *s* for such lineages is zero. For such lineages, equation (S9) simplifies to equation (1) in the main text, which can be used to infer *µ*_*k*_ and therefore 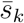. However, instead, Levy et al construct the entire empirical distribution for *r*_*k*_ and fit equations (S8), (S9) to it, using 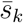 and *κ*_*k*_ as free parameters (see equations (45)–(48) in the Supplementary Information to Ref. [3]). Having inferred the 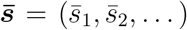 and ***κ*** = (*κ*_1_, *κ*_2_, …) from low-abundance lineages, the LB approach takes the entire barcode read-count trajectory ***r***_*i*_ = (*r*_*i*1_, *r*_*i*2_, …) for each lineage *i* to infer its selection coefficient *s*_*i*_ and the establishment time *τ*_*i*_, using a Bayesian approach. Specifically, they define the likelihood function for the read-count trajectory ***r***_*i*_ as

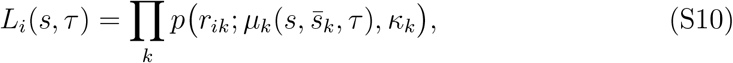

where *p* is given by equation (S8) and an expression for 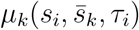 that is adjusted to account for the fact that an adapted lineage contains a subpopulation of adapted cells with *s*_*i*_ *>* 0 growing from established time *τ*_*i*_ and a non-adapted subpopulation (see equations (58)–(62) in the Supplementary Information to Ref. [3]). To determine if lineage *i* is adapted or not, they construct the ratio of posterior probabilities as

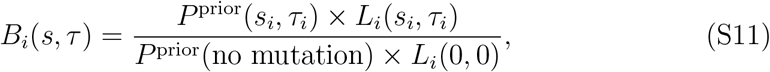

where *P* ^prior^ denote prior probability for each hypothesis (see equations (64)–(67) in the Supplementary Information to Ref. [3]). Finally, they find values *ŝ*_*i*_ and *τ*_*i*_ that maximize *B*_*i*_. Lineage *i* is identified as adapted if *B*_*i*_(*ŝ*_*i*_, *τ*_*i*_) *>* 1.

#### 2.2 Review of FitMut2

FitMut2 is an improved algorithm based on the LB method. The key difference between the LB method and FitMut2 is that FitMut2 uses an iterative approach to simultaneously call adapted lineages and infer mean fitness from them in a consistent way, which removes the need for a large number of neutral lineages required by the LB method. For convenience, we provide the summary of similarities and differences between all the BLT analysis methods used in this paper in Table S4.

Specifically, for each lineage *i*, FitMut2 considers two hypotheses, that the lineage acquired an adaptive mutation (Θ = 1) or not (Θ = 0) during the BLT experiment. FitMut2 calculates the probability for the observed read-count trajectory ***r***_*i*_ for lineage *i* as

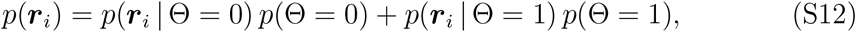

where *p* (Θ = 1) and *p* (Θ = 0) are the prior probabilities of the adaptive and neutral hypotheses, respectively.

For the neutral model (Θ = 0), they set *p*(Θ = 0) = 1 and expand the probability of the trajectory as

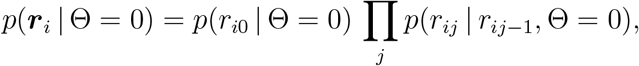

where *p*(*r*_*i*0_ |Θ = 0) = *p*(*r*_*i*0_) is the initial prior (see equations (14)–(15) in the Supplementary Information to Ref. [4]) and the transition probability *p*(*r*_*ij*_ | *r*_*ij*−1_, Θ = 0) is given by equation (S7)) with the expected read count *µ*_*k*_ given by equation (S9) with *s* = 0 (see equation (11) in the Supplementary Information to Ref. [4]).

For the adaptive model (Θ = 1), they express the probability *p* (***r***_*i*_ | Θ = 1) *p* (Θ = 1) as an integral over all possible ways to be an adapted lineage with fitness effects *s* and establishment time *τ*

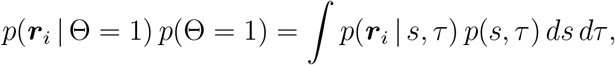

where *p*(*s, τ*) is the prior probability that a lineage acquires a beneficial mutation (defined in the Discussion of Ref. [4]). Then the probability of the adaptive trajectory is factorized as

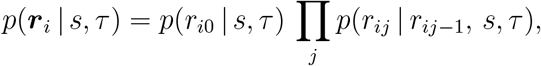

where *p*(*r*_*i*0_ | *s, τ*) = *p*(*r*_*i*0_) is the same initial prior as in the neutral model and *p*(*r*_*ij*_ | *r*_*ij*−1_, *s, τ*) is given by equation (S7) with the same expected read count *µ*_*k*_ as in the LB method (essentially equation (S9) adjusted for the fact that the adaptive lineage is segregating within the barcoded subpopulation), defined in equation (13) in the Supplementary Information to Ref. [4].

To choose the best model for lineage *i*, FitMut2 applies Bayes’ theorem to calculate the posterior probability of Θ = 1, given the data ***r***_*i*_,

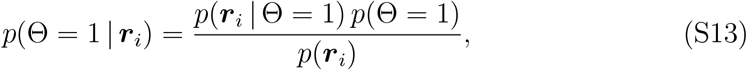

where *p* (*{r*_*ik*_*}*) is given by equation (S12). If *p* (Θ = 1 | ***r***_*i*_) *>* 0.5, lineage *i* is called putatively adaptive, with putative estimates of *s* and *τ* obtained by maximizing the posterior log-likelihood *L*(*s, τ*) = ln (*p*(***r***_*i*_ | *s, τ*) *p*(*s, τ*)).

Then, they calculate the mean fitness trajectory 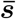 at time *k* from the frequencies *f*_*ik*_ of all putatively adapted lineages *i*

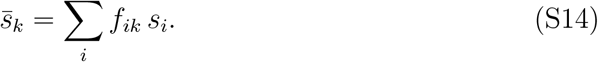

With the newly estimated mean fitness trajectory 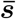, FitMut2 then re-identifies putatively adaptive lineages and re-estimates their selection coefficients and establishment times. FitMut2 then continues this process iteratively until it converges.

#### 2.3 Key assumptions of the Levy-Blundell method

The LB method—and the neutral decline method as its simplified version—is based on two key assumptions. In this section, we discuss their validity and possible consequences for inference if they are violated.

##### 2.3.1 Assumption 1. Low-abundance lineages are neutral

Low-abundance lineages can be used to infer the mean-fitness trajectory as long as these lineages are neutral, i.e., as long as they have not yet acquired adaptive mutations. If the lineages chosen as reference are in fact not neutral, the inferred mean fitness will be biased, as we demonstrate in the main text. Thus, it is critical for the accuracy of the LB method that this assumption holds.

Of course, as adaptation proceeds, all neutral lineages eventually go extinct and this assumption will eventually be violated. The time scale of persistence of neutral lineages depends on the speed of adaptation, which in turn is determined by the availability of adaptive mutations and their fitness effects, and is therefore a priori unknown.

This assumption can be relaxed if we allow for adapted lineages with known fitness to be used as reference. This can be implemented in practice as follows. By definition, in an initially clonal population, most cells are “wildtype” and any lineage can be used as reference at the initial phase of adaptation. Some lineages will acquire adaptive mutations earlier than others and their fitness can be estimated reliably while neutral lineages are still present in the population. These early adapted lineages can be then used as reference themselves until they acquire secondary adaptive mutations.

##### 2.3.2 Assumption 2. Read frequency dynamics faithfully represent lineage frequency dynamics

According to the population genetics theory, the expected rate of change *dx/dt* in the frequency of a mutant is determined by its current frequency *x*, its selection coefficient *s* and population’s mean fitness 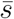,

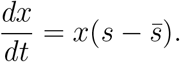

Taking the mean fitness as constant 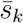 over a short period of time Δ*t*_*k*_ = *t*_*k*_ − *t*_*k*−1_ (e.g., between two consecutive sampling time points *t*_*k*−1_ and *t*_*k*_ in the BLT experiment), we integrate this equation and obtain the expected frequency *x*_*k*_ at time *t*_*k*_,

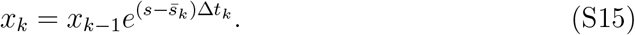

Thus, if the lineage frequencies are known, we can solve equation (S15) for 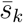 (if *s* is known) or for *s* (if 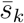 is known).

However, *x*_*k*_ are never known with certainty but are estimated. In particular, in BLT experiments, *x*_*k*_ must be estimated from the corresponding barcode readcount data. The LB method (and the neutral decline method as a consequence) makes an implicit but crucial assumption that the dynamics of measured barcode *read* frequencies are an unbiased (albeit noisy) representation of the dynamics of *cell* frequencies. While this assumption is reasonable, it is in general not true.

We will first show that this assumption is indeed implied by equation (S9). Since *µ*_*k*_ is the expected number of reads at time *t*_*k*_, equation (S9) can also be re-written as

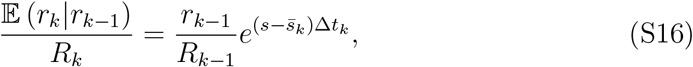

which implies that the expected read frequency *E* (*r*_*k*_|*r*_*k*−1_) */R*_*k*_ depends on the previous read frequency *r*_*k*−1_*/R*_*k*−1_ in exactly the same way as the cell frequency *x*_*k*_ in equation (S15).

We will now show that equation (S16) is in fact not generally true. Suppose that the measurement process is described by the probability distribution *P* ^meas^(*r* | *x*; *R*), which determines the number of reads *r* that are obtained from a lineage whose actual population frequency is *x* if the total read depth is *R*. Let us assume that measurement is unbiased, i.e.,

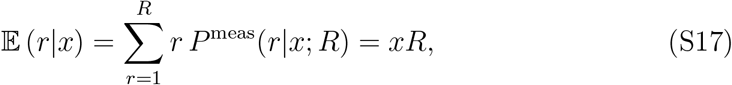

which is supported by our data (see Figure 4A in the main text). We rewrite *E* (*r*_*k*_ | *r*_*k*−1_) in terms of the measurement process *P* ^meas^ and the evolutionary process *P* ^evol^.

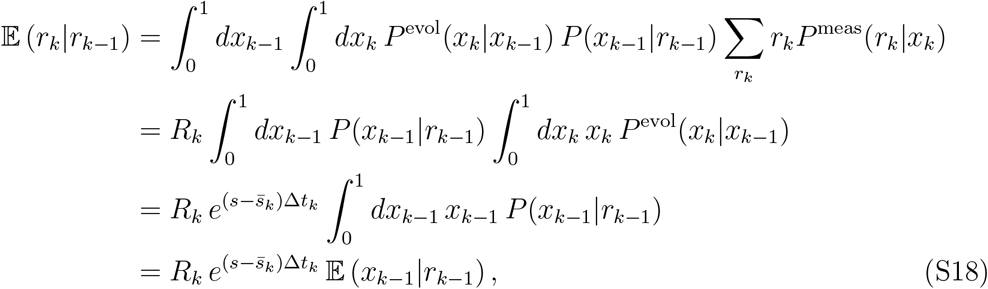

where to simplify the notations we dropped the parameters from the expressions for various probability distributions. equation (S18) shows that equation (S16) would be true if the expected frequency of lineages represented by *r* out of *R* reads is in fact *r/R*,

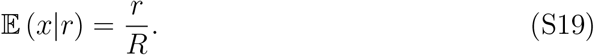

Counterintuitively, equation (S19) is in general not true. To demonstrate this, we first use the Bayes’ theorem to re-write it as

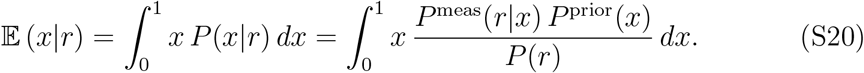

Here, *P* (*r*) is the normalization constant and *P* ^prior^(*x*) is the a priori probability that the frequency of the focal lineage in the population is *x*. Note that in general *P* ^prior^(*x*) is not the distribution of lineage frequencies in the population. Indeed, at later time points in the BLT experiment, *P* ^prior^(*x*) for different lineages are different, because they are informed by previous observations. However, at the beginning of the BLT experiment, all lineages are a priori identical, and *P* ^prior^(*x*) can be approximated by the distribution of lineage frequencies in the population. Previous BLT studies have shown that, despite efforts to introduce all barcodes at the same frequency, the initial distributions of barcode frequencies are often be quite broad and may be close to exponential [2]. We now use four specific examples to show that equation (S19) does not hold.

**Example 1. Poisson measurement noise and a uniform frequency distribution.** First, we consider the simplest case, when the prior frequency distribution is uniform and the measurement noise is Poisson, i.e.,

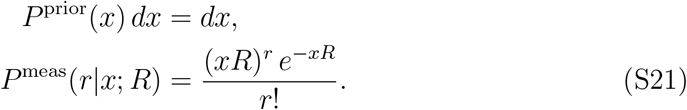

We show in Appendix B, that the conditional distribution for frequency *x* given the read count *r* is approximately a gamma distribution with shape parameter *r* + 1 and scale parameter 1*/R*, such that

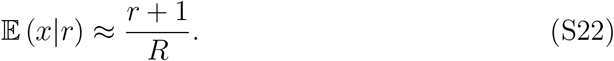

Reassuringly, when *r* ≫ 1, expression (S22) converges to *r/R*, suggesting that Assumption 2 is reasonably accurate as long as all lineages are represented by many reads. However, at small numbers of reads, expression (S22) can deviate substantially from equation (S19).

**Example 2. Poisson measurement noise and an exponential frequency distribution.** We now consider an exponential frequency distribution, which may be adequate for initial time points [2],

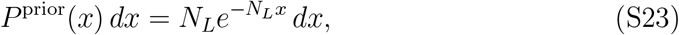

where *N*_*L*_ is the number of barcoded lineages, such that *E* (*x*) = 1*/N*_*L*_. We show in Appendix B, that the conditional distribution for frequency *x* given the read count *r* is approximately a gamma distribution with shape parameter *r* + 1 and scale parameter 1*/*(*R* + *N*_*L*_), such that

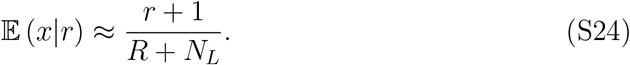

Since *R/N*_*L*_ ≳ 100 in typical BLT experiments, expression (S24) is still close to *r/R* for large *r*, but it again can deviate substantially from (S19) when the number of reads is small.

**Example 3. Measurement noise with an increasing variance to mean ratio and a Gamma frequency distribution.** We now consider a more realistic noise model, where the variance to the mean ratio increases with the mean according to equation (2) in the main text. We model the read count as a negative binomial random variable with mean *xR* and the variance to the mean ratio 1 + *ϵxR*.

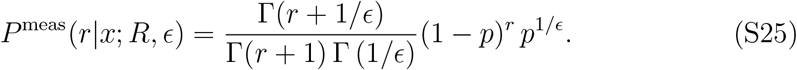

with *p* = (1 + *ϵxR*)^−1^. We also consider a Gamma frequency distribution with shape parameter *α* and scale parameter 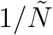,

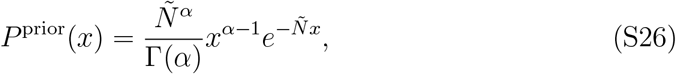

which may be a more accurate description of our prior knowledge at later time points with 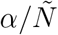 representing the a priori expected frequency. If *≪* 1 and *r* ≪ 1*/ϵ*, the posterior distribution is approximately Gamma with shape parameter *r* + *α* and scale parameter 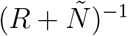 (see Appendix B), such that

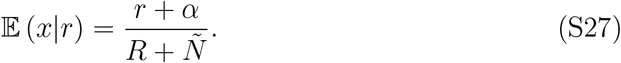

**Example 4. Measurement noise with a constant variance to mean ratio and an exponential frequency distribution.** Finally, we consider a noise model where the variance to the mean ratio is constant and equal to 2*κ* as in Ref. [3]. We again model the read count as a negative binomial random variable,

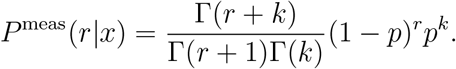

where *p* = 1*/*2*κ* and *k* = *xR/*(2*κ* − 1). In this case, as we show in Appendix B, we have

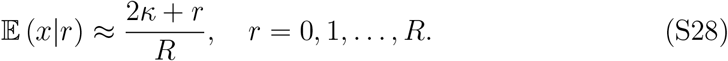

Substituting any of the equations (S22), (S24), (S27) or (S28) into equation (S18), we see that the relationship between the expected number of reads at time *t*_*k*_ and the observed number of reads at *t*_*k*−1_ is not the same as the relationship (S15) between the expected lineage frequency at time *t*_*k*_ and the actual frequency at *t*_*k*−1_.

Specifically, unlike the relationship between *E* (*x*_*k*_|*x*_*k*−1_) and *x*_*k*−1_, the relationship between *E* (*r*_*k*_ | *r*_*k*−1_) and *r*_*k*−1_ appears to generally have a non-zero *y*-intercept, whose value depends on the parameters of the prior distribution and measurement noise.

However, these two relationships become identical when *r*_*k*−1_ is sufficiently large, although how large it needs to be depends on the noise model.

### 3 BASIL

#### 3.1 Model of measurement

As discussed in the main text, we cannot directly observe the size *n* of a barcoded lineage. Instead, a sample from the population is sequenced, and we observe the number of reads that contain the focal barcode.

We model the measurement process with the negative binomial distribution with mean ⟨*r*⟩ = *nR/N* and variance 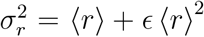 (see Section 1), where *N* is total number of cells in the population during sampling, *R* is the total coverage, and *r* is the number of reads that contain the focal barcode, and *ϵ* is a free parameter controlling overdispersion. The negative binomial distribution is typically parameterized by the number of “successes” *k* and the success probability *p*, with mean and variance expressed in terms of these parameters as

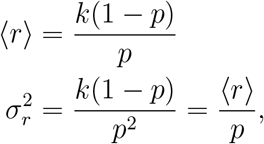

which yield

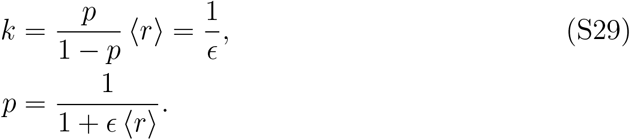

Thus, the probability of observing *r* reads for lineage with *n* cells is

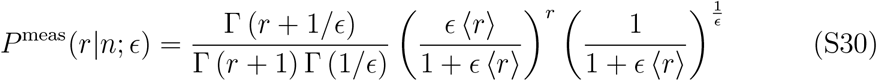

with ⟨*r* ⟩ = *nR/N*. The measurement distribution (S30) depends on the noise parameter *ϵ*, which we fit, as well as on *N* and *R* which are assumed to be known. Furthermore, this distribution becomes a point measure at *r* = 0 if *n* = 0 and it converges to the Poisson distribution with mean ⟨*r*⟩ = *nR/N* as *ϵ* → 0.

#### 3.2 The belief distribution for lineage size and fitness

Our main object is the belief probability distribution *P* ^belief^ (*n, s* | ***r***) that a lineage with the observation vector ***r*** = (*r*_0_, …, *r*_*k*_) has size *n* and selection coefficient *s*. For a given lineage *i* whose observation vector up to and including time point *t*_*k*_ is ***r***_*ik*_ = (*r*_*i*0_, …, *r*_*ik*_), we will sometimes use a simplified notation 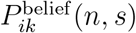 instead of *P* ^belief^ (*n, s* | ***r***_*ik*_). For mathematical tractability, we express the belief distribution as

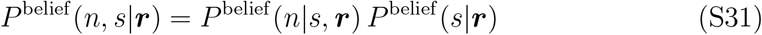

and assume the following parametric forms

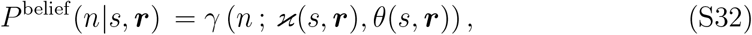

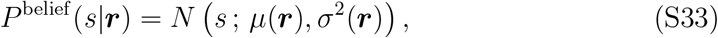

where *N* (*s*; *µ, σ*^2^) is the normal distribution with mean *µ* and variance *σ*^2^ and *γ* (*n*; *x*, *θ*) is the gamma distribution with shape parameter *x* and scale parameter *θ*.

Thus, *P* ^belief^ (*n, s* | ***r***) belongs to a family of distributions with four parameters *µ, σ*^2^, *x*, *θ* where the latter two parameters can be functions of *s*. In this section, we describe how we update these parameters as the observation vector is augmented from one time point to the next. We do so in two steps, which we refer to as “projection” and “update”. During the projection step, which is described in Section 3.2.1, we project the past belief distribution 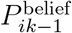 to obtain the prior distribution 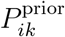 for the time point *t*_*k*_. During the update step, which is described in Section 3.2.2, we use the Bayes’ theorem to obtain the new belief distribution 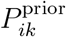 based on the prior distribution 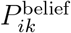 and the read count *r*_*ik*_ observed at time *t*_*k*_ (see equation (3) in the main text). Importantly, the parametric distribution given by equation (S32) has no weight at *n* = 0. In other words, it is valid only for lineages that are present in the population and have not yet gone extinct. We describe how we treat lineage extinctions in Section 3.2.2.

##### 3.2.1 Projecting the belief distribution

To obtain the prior distribution *P* ^prior^(*n, s*|***r***^′^) for the time point *t*_*k*_, where ***r***^′^ = (*r*_0_, …, *r*_*k*−1_) is the observation vector up to and including the previous time point *t*_*k*−1_, we assume that the belief distribution *P* ^belief^ (*n, s* | ***r***^′^) at the previous time point is known and has the parametric form given by equations (S32), (S33) with parameters *µ, σ, θ*, *x*. To simplify notations, in this section, we will omit the explicit dependence of all probabilities on the observation vector ***r***^′^.

The sampling time points *t*_0_, *t*_1_, … in the BLT experiment may be separated by one or multiple growth and dilution cycles. We will first consider one such cycle and obtain the “1-cycle projected distribution” *P*_1_ (*n, s*) whose parameters we denote as *µ*_1_, *σ*_1_, *ϰ*_1_, and *θ*_1_. We then use *P*_1_ (*n, s*) to obtain the prior probability for *n* and *s* after multiple cycles.

To derive *P*_1_ (*n, s*), we consider a barcoded lineage that has fitness *s* relative to the ancestor and that is represented by *n* cells immediately prior to dilution. At dilution, a fraction 1*/D* of the population is transferred into fresh medium, and the rest is discarded. Immediately after dilution, the lineage’s size becomes *n*^′^, which we model as a Poisson random variable with mean *n/D*, such that

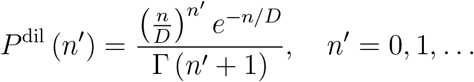

We assume that after dilution, the lineage grows deterministically based on the difference between its fitness *s* and the population’s mean fitness 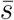, such that by the end of the cycle, its size is *n*_1_ = *An*^′^, where

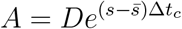

and Δ*t*_*c*_ is the cycle duration in generations. Then, the one-cycle transition probability is

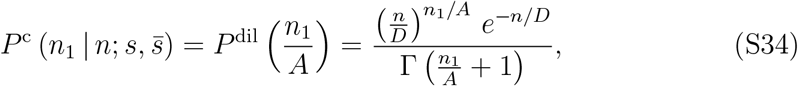

s -where formally *n*_1_*/A* = 0, 1, 2, …. Then, for the 1-cycle projected distribution we have

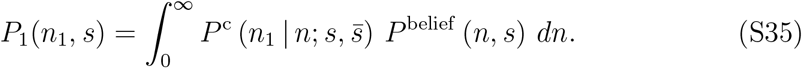

Integrating equation (S35) with respect to *n*_1_, we find, as expected, that the projection does not change the marginal distribution for *s*,

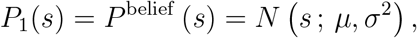

which implies that *µ*_1_ = *µ* and *σ*_1_ = *σ*. Since we can express the joint distribution *P*_1_ (*n*_1_, *s*) through this marginal distribution as *P*_1_ (*n*_1_, *s*) = *P*_1_ (*n*_1_ | *s*) *P*_1_ (*s*), we now only need to obtain the conditional probability *P*_1_ (*n*_1_ | *s*). To do so, we note that

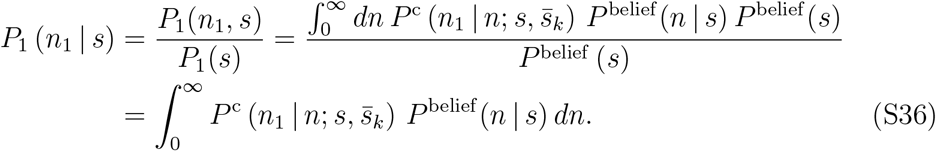

Substituting expressions (S34) and (S32) into equation (S36), we obtain

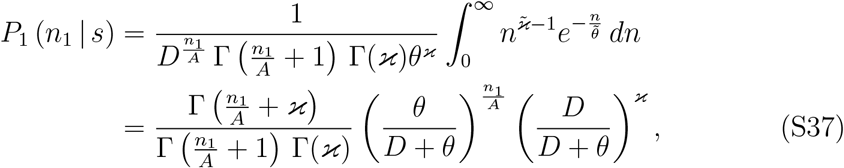

with 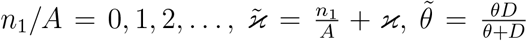. equation (S37) shows that, conditional on *s, n*_1_*/A* is distributed as negative binomial with success probability *D/*(*D* + *θ*) and number of successes *ϰ*. Therefore,

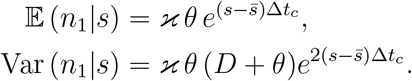

Furthermore, the lineage goes extinct after one growth and dilution cycle with probability

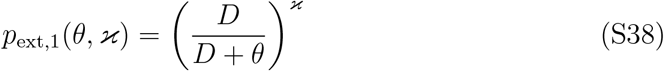

and survives with probability

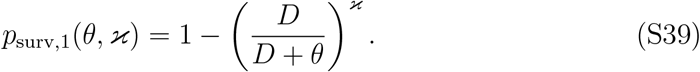

It is then easy to show that the mean and variance conditional on survival are

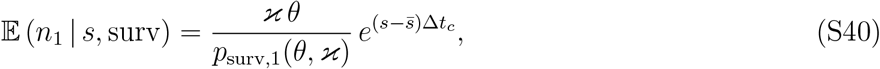

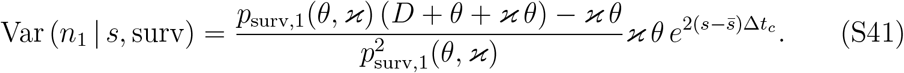

To facilitate further calculations, we approximate the distribution of *n*_1_, conditional on *s* and lineage survival, by a gamma distribution with the shape parameter *ϰ*_1_ and scale parameter *θ*_1_. To determine *ϰ*_1_ and *θ*_1_, we equate the mean *ϰ*_1_*θ*_1_ and variance 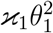 of this gamma distribution to the conditional mean and variances given by equations (S40) and (S41) and find *θ*_1_ = *f*_*θ*_(*s, θ, ϰ*) and *x*_1_ = *f*_*x*_(*θ, ϰ*) where

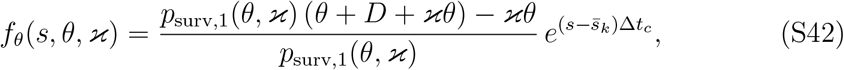

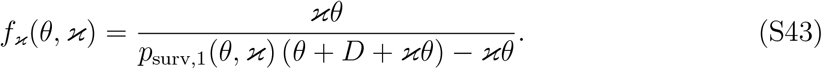

Therefore, the full projected probability for a lineage after one growth and dilution cycle is

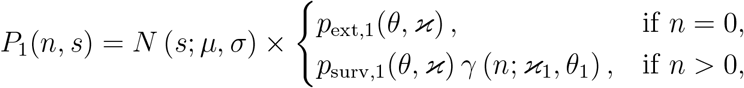

where *p*_ext,1_ and *p*_surv,1_ are given by equations (S38) and (S39) and parameters *θ*_1_ and *x*_1_ are the functions of *θ* and *x* given by equations (S42) and (S43).

If the next sampling occurs at the end of a single growth and dilution cycle after the previous sampling, then we set *P* ^prior^(*n, s*) = *P*_1_(*n, s*). If *L >* 1 growth and dilution cycles elapse between successive samples, then we set the prior probability *P* ^prior^(*n, s*) for the next sampling time point to be equal to

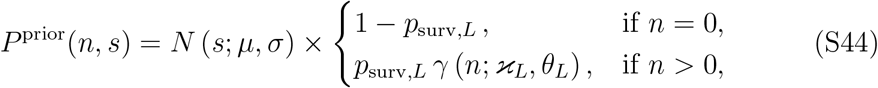

where the *L*-cycle survival probability is given by

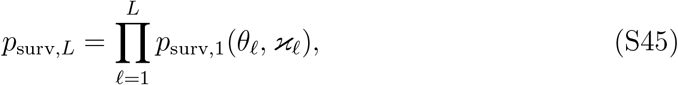

and parameters *θ* = *f*_*θ*_ (*s, θ*_−1_, *x*_−1_), *x* = *f*_*x*_ (*θ*_−1_, *x*_−1_) for *f_* = 1, 2, … *L* are obtained recursively using equations (S42) and (S43) with *x*_0_ ≡ *x*, *θ*_0_ ≡ *θ*. Note that since all *x* and *θ* depend on the parameters *x*, *θ* of the prior belief distribution *P* ^belief^ (*n* | *s*, ***r***^′^), which themselves are functions of *s*, the survival probability *p*_surv,*L*_ is also a function of *s*.

##### 3.2.2 Updating the belief distribution

Next, we apply the Bayes’ theorem to obtain the belief distribution *P* ^belief^ (*n, s* | ***r***) after observing the current read count *r*_*k*_,

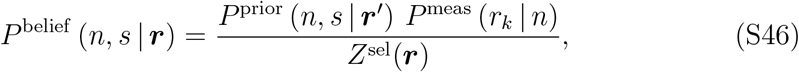

where ***r***^′^ = (*r*_0_, …, *r*_*k*−1_) and ***r*** = (*r*_0_, …, *r*_*k*−1_, *r*_*k*_) are the previous and current observation vectors, *P* ^prior^ (*n, s* | ***r***^′^) is the prior probability that the lineage has selection coefficient *s* and size *n* at the current observation time point before the current measurement is made (see equation (S44)), *P* ^meas^ (*r* | *n*) is the probability of observing *r* reads for a lineage of size *n* (see equation (S30)), and the denominator

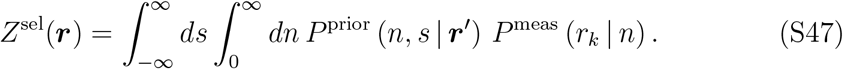

is the marginal probability of observing *r*_*k*_ reads in our model. The superscript “sel” stands for “selection” and denotes the fact that our model allows the selection coefficient of the lineage to vary freely. We will later consider a neutral null model where the selection coefficient is fixed at zero (see Section 3.3). Note that *P* ^prior^ depends on 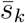, the mean fitness at the current interval (*t*_*k*−1_, *t*_*k*_), and *P* ^meas^ depends on the noise parameters *ϵ*_*k*_ at the current interval (*t*_*k*−1_, *t*_*k*_). Therefore, both *Z*^sel^ and *P* ^belief^ depend on both 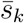 and *ϵ*_*k*_. For now, we can ignore these dependencies treating these parameters as fixed and known. However, these dependencies will becomes important in Section 3.3, where we discuss how 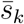 and *ϵ*_*k*_ are estimated.

As discussed above, we would like to approximate the belief distribution with an analytical parametric form given by equations (S32), (S33). However, since the numerator of equation (S46) is a complex function of both *n* and *s*, the normalization constant (S47) and the moments of this distribution cannot be expressed analytically. Therefore, to estimate the parameters of the belief distribution, we draw a random sample (*n*_*j*_, *s*_*j*_), *j* = 1, 2, …, *M* from the belief distribution *P* ^belief^ (*n, s* | ***r***) using the Markov Chain Monte Carlo (MCMC) approach (see Section 3.5 for the details of the algorithm). We consider two cases, *r*_*k*_ *>* 0 and *r*_*k*_ = 0, which differ qualitatively because in the former case, the lineage is guaranteed to have survived until the current sampling time point *t*_*k*_ whereas in the later case it is possible that the lineage has gone extinct.

**Guaranteed lineage survival when *r*_*k*_ *>* 0.** If *r*_*k*_ *>* 0, that is, if the lineage is observed at current time point *t*_*k*_, then *P* ^belief^ (0, *s* | ***r***) = 0 since *P* ^meas^(*r*_*k*_ | 0) = 0 whenever *r*_*k*_ *>* 0. In other words, if we observe the lineage, we know with certainty that it has survived up to the current sampling time point. We then estimate the parameters *µ, σ* of the distribution (S33) from this sample as

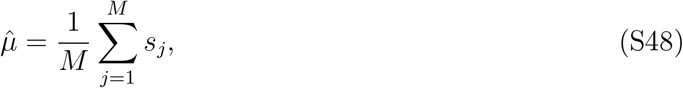

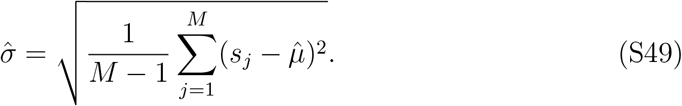

Estimating parameters *x*(*s*) and *θ*(*s*) of the conditional distribution (S32) is more difficult because they can be arbitrary functions of *s*. In principle, one could estimate these functions by independently fitting the shape and scale parameters of the gamma distribution for each *s*-slice of the joint distribution. However, this approach is computationally intensive. To reduce computational burden, we sought to find simple functional forms for the functions *x*(*s*) and *θ*(*s*) and fitting the parameters of these functional forms directly from the joint distribution. To this end, we no-ticed that ln n and s are linearly correlated in our MCMC samples (see Figure S3), implying that

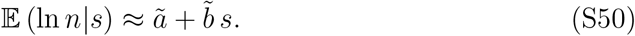

Since

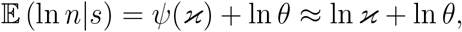

where *ψ* is the di-gamma function (see Appendix C) and the approximation *ψ*(*x*) ≈ ln *x* holds when *x ≫* 1, we obtain

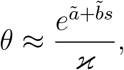

which suggests that an exponential dependence of *θ* on *s* should capture the shape of our belief distribution. To simplify subsequent calculations, we use the ansatz

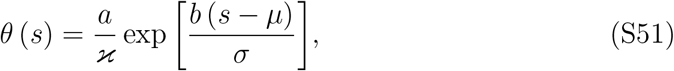

where *x >* 0, *a* and *b* are the new real-valued parameters of distribution (S32).

With this parametrization, we have

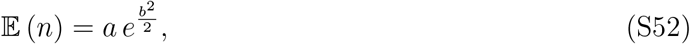

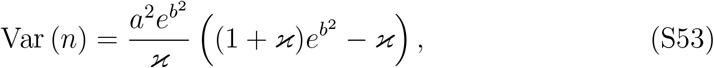

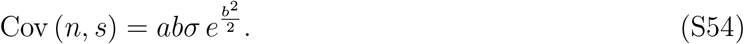

Equations (S52)–(S54) can be solved to yield estimators of the parameters of the gamma distribution (S32) with the functional form (S51),

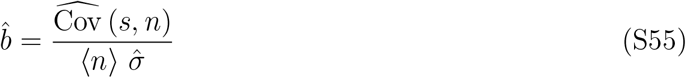

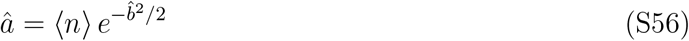

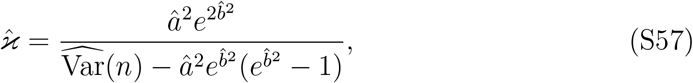

where

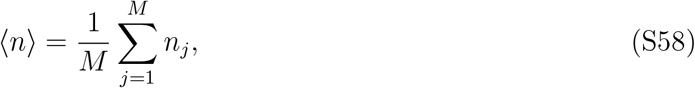

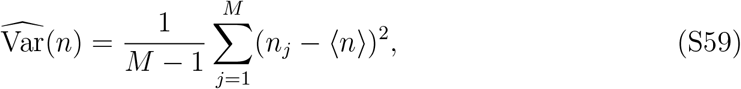

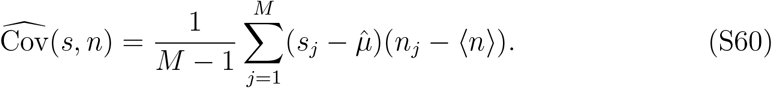

To test how well our parametric form (S32), (S33), (S51) with the parameters fitted using equations (S55)–(S60) fits the MCMC-sampled posterior distribution, we selected four lineages from simulated and real data and then plotted their MCMC-sampled marginal distributions for *s* and *n* as well as their joint distribution (Figure S3). To calculate the marginal distribution for *n*, we generated *M* ^′^ = 10^5^ random samples of *s*_*j*_, *j* = 1, …, *M* ^′^ from the normal distribution with mean 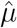 and variance 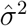. We then approximate the marginal distribution for *n* as

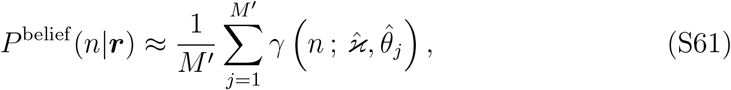

where 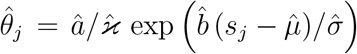 with *s*. Figure S3 shows that our parametric form captures the MCMC-sampled distribution reasonably well.

**Possible lineage extinction when *r*_*k*_ = 0.** If *r*_*k*_ = 0, there are two possibilities. With probability *p*_surv,*L*_ given by equation (S45), the lineage has survived up to the current sampling time point *t*_*k*_ but was not detected due to sampling noise; or, with probability 1 − *p*_surv,*L*_, the lineage has gone extinct between *t*_*k*−1_ and *t*_*k*_. Therefore, the belief distribution *P* ^belief^ (*n, s*|***r***) has a non-zero weight at *n* = 0,

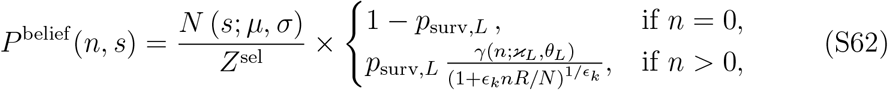

with

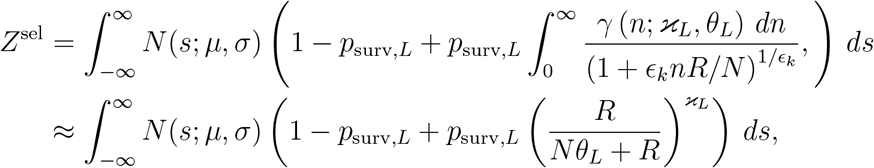

where the approximation holds when *ϵ*_*k*_ *≪* 1, and we have omitted the dependencies on the observation vector ***r*** and mean fitness *s*_*k*_.

equation (S62) shows that the belief distribution can no longer be adequately captured by the parametric form given by equations (S32), (S33). Instead, we use this parametric form to capture the conditional belief distribution, conditional on the lineage having survived until the current sampling time point *t*_*k*_. We do so using the same approach as above, i.e., we obtain an MCMC sample (*n*_*j*_, *s*_*j*_), *j* = 1, 2, …, *M* from the distribution (S62) and then use equations (S58)–(S60) to estimate the mean and the variance of the lineage size *n* and the covariance between lineage size and selection coefficient, except we use only those MCMC samples with *n*_*j*_ *>* 0. We then apply equations (S48), (S49), (S55)–(S57) to estimate the parameters of the conditional belief distribution (S32), (S33). Furthermore, if the lineage was not observed at two consecutive time points, we assume that it has gone extinct and stop tracking it.

#### 3.3 Estimating populations mean fitness and the noise parameter

As mentioned above, our model of evolution and consequently our procedure for updating the belief distribution at the “current” time point *t*_*k*_ depends on the population’s mean fitness 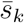 during the interval (*t*_*k*−1_, *t*_*k*_), *k ≥* 1 as a parameter (see equation (S34)). Similarly, our measurement distribution (equation (S30)) depends on the noise parameter *ϵ*_*k*_ during the interval (*t*_*k*−1_, *t*_*k*_). So far, we have assumed that both 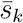 and *ϵ*_*k*_ are fixed and known. However, in reality the true values of 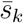 and *ϵ*_*k*_ are of course unknown and must be estimated before we update the lineage belief distributions at *t*_*k*_.

We estimate 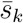 and *ϵ*_*k*_ using the maximum likelihood approach. To this end, we choose 3000 random lineages for which *r*_*ik*−1_ *>* 0 and classify each of them as either putatively neutral or putatively adapted (at the current time interval (*t*_*k*−1_, *t*_*k*_)) as described below. We denote the subsets of currently putatively neutral and putatively adapted lineages as *N*_*k*_ and *A*_*k*_, respectively. For each putatively adapted lineage *i* ∈ *A*_*k*_, we can treat the normalization constant 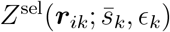 given by equation (S47) as the likelihood of observing the number of reads *r*_*ik*_ for that lineage at time *t*_*k*_ under the model with selection. Similarly, for each putatively neutral lineage *i* ∈ *N*_*k*_, we can treat the normalization constant 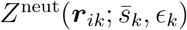 under the neutral model described below (see equation (S66)) as the likelihood of observing the number of reads *r*_*ik*_ for that lineage at time *t*_*k*_ under the neutral model. The procedures for estimating 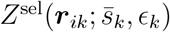 and 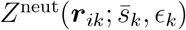 are described below. Since both 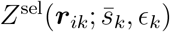 and 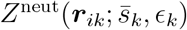 depend on the unknown parameters 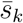 and *ϵ*_*k*,_ we can write down the log-likelihood function for 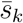 and *ϵ*_*k*_ as

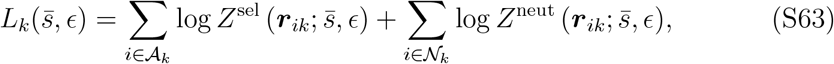

and estimate

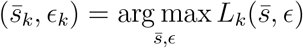

using standard optimization algorithms.

**Classification of lineages for estimating mean fitness.** We classify lineages into putatively neutral and putatively adapted to minimize estimation biases that may arise and be amplified over time due to the following positive feedback loop. Evolutionary dynamics in our model depend only on the difference between the fitness of a lineage and the mean fitness (see equations (S42),(S43)), but not on these two quantities individually. Thus, all selection coefficients and the mean fitness can be identified from data only up to a shared constant. We eliminate this ambiguity by assuming that the expected fitness of all lineages is zero at *t*_0_ (see Section **??**). In other words, we estimate selection coefficients with respect to population’s initial mean fitness. While this condition should be sufficient to accurately estimate the mean fitness and lineage selection coefficients on average, the large uncertainty in our priors can lead to large uncertainties in the initial estimates of lineage fitness (Figure S4) and mean fitness. These initial deviations can over time be “baked in” into the belief distributions because, as mentioned above, additional data can only correct any inaccuracies in the difference between lineage fitness and mean fitness but not in their individual values.

To make this reasoning more concrete, suppose that we do not classify lineages into putatively neutral and adapted and instead use lineage belief distributions to estimate mean fitness, i.e., use only *Z*^sel^ terms for all lineages in equation (S63). Imagine that to estimate mean fitness 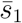 at the first time interval (*t*_0_, *t*_1_), by chance we pick 3,000 lineages whose frequency declines (slightly) more than expected under neutrality. Then, we will (slightly) over-estimate 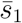. As a result, the projected distributions would under-estimate lineage sizes at *t*_1_, thus generating an overly strong surprise once the read counts are observed. This would push us to update the belief distribution towards overly high *s* for many lineages, which would in turn cause an additional over-estimate of the mean fitness at the next time interval (*t*_1_, *t*_2_), etc.

To dampen this positive feedback loop at the mean-fitness estimation step, we assume that all lineages are neutral by default (for the purposes of mean-fitness estimation) unless we have a high degree of confidence that a given lineage is adapted. Specifically, we classify lineage *i* as putatively adapted for the purposes of estimating 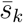 if

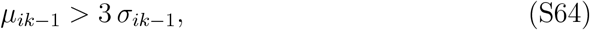

where

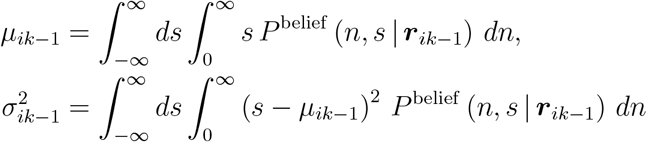

are the mean and variance of the marginal belief distribution for *s* for lineage *i* at time *t*_*k*−1_ estimated using equations (S48), (S49). The factor 3 in equation (S64) was chosen so that our confidence that the lineage is in fact adapted (i.e., has a positive *s*) exceeds 99%.

**Neutral model of lineage evolution.** To compute the marginal likelihood 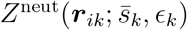 that *r*_*ik*_ reads are observed for lineage *i* at time point *t*_*k*_ if the lineage is neutral during the time interval (*t*_*k*−1_, *t*_*k*_), we employ a neutral model that is analogous to the model with selection described above in Sections 3.2.1 and 3.2.2.

As in the model with selection, we start with the belief distribution *P* ^belief^ (*n, s* | ***r***^′^), where ***r***^′^ = (*r*_0_, …, *r*_*k*−1_) is the observation vector up to and including the previous time point *t*_*k*−1_. Since the neutral model is formulated only in terms of the lineage size *n*, we first need to obtain the marginal belief distribution *P* ^belief^ (*n* | ***r***^′^). This distribution has no analytical expression. Therefore, to facilitate further calculations, we approximate it with a Gamma distribution with the scale parameter 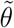 and shape parameter 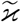 chosen to match the marginal mean and variance given by equations (S52), (S53) i.e., we set

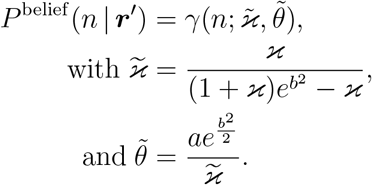

We then obtain the prior distribution for the lineage size at the current time point *t*_*k*_ analogously to equation (S44),

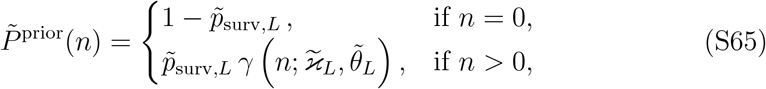

where the *L*-cycle survival probability 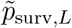 is given by equation (S45) with pa-rameters 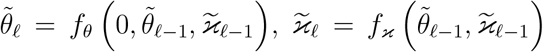 for *f_* = 1, 2, … *L* that are obtained recursively using equations (S42) and (S43) with 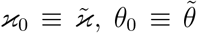. The marginal likelihood under the neutral model is then given by

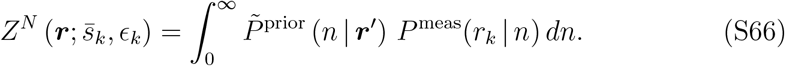

**Numerical estimation of the marginal likelihoods *Z*^sel^ and *Z*^neut^.** For every putatively adapted lineage, we estimate the marginal likelihood *Z*^sel^ given by equation (S47) by first obtaining a random sample (*s*_*j*_, *n*_*j*_), *j* = 1, 2, …, *M* with *M* = 5, 000 from the projected distribution 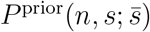 for that lineage, which is given by equation (S44), using a standard random-number generator, keeping only those samples with *n*_*j*_ *>* 0, i.e., conditional on lineage survival. We then calculate

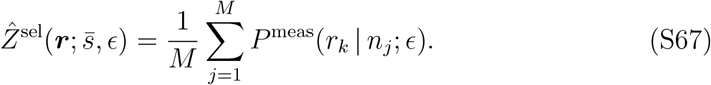

For every putatively neutral lineage, we estimate the marginal likelihood *Z*^neut^ given by equation (S66) analogously, by first obtaining a random sample *n*_*j*_, *j* = 1, 2, …, *M* from the projected distribution 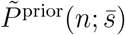 given by equation (S65), keeping only those samples with *n*_*j*_ *>* 0. We then estimate 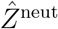 as per equation (S67).

#### 3.4 Identification of adapted lineages and estimation of their selection coefficients

As a result of applying BASIL, we obtain a time-varying belief distribution 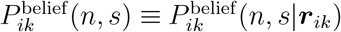 for each lineage *i* at each sampling time point *t*_*k*_, *k* = 1, 2, Our primary interest is in the marginal belief distribution for the selection coefficient 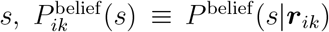 whose mean and variance are *µ*_*ik*_ and 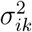, estimated using equations (S48), (S49). At early time points, we expect these belief distributions to have high variance because they are insufficiently constrained by the data. As data accumulates, the uncertainty in *s* should decline, particularly for lineages that actually acquired a single adaptive mutation (Figure S4). However, if we wait long enough, secondary adaptive mutations might appear, which could again increase the uncertainty in our belief of *s*. Thus, for each lineage *i*, we find the time point 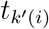 with minimal variance, 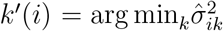. We call lineage *i* adapted if 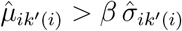 where *β* is the “confidence factor”, a hyper-parameter that controls the precision and recall of our inference. As described in the main text, we empirically determine that the confidence factor *β* = 3.3 maximizes the F1-score (the harmonic mean of precision and recall) in our simulated data, and use this value for all our analyses. For each lineage *i* called as adapted, we estimate its selection coefficient as 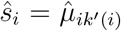, and we can calculate the credible interval for it using the standard deviation 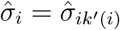.

##### Algorithm 1 BASIL

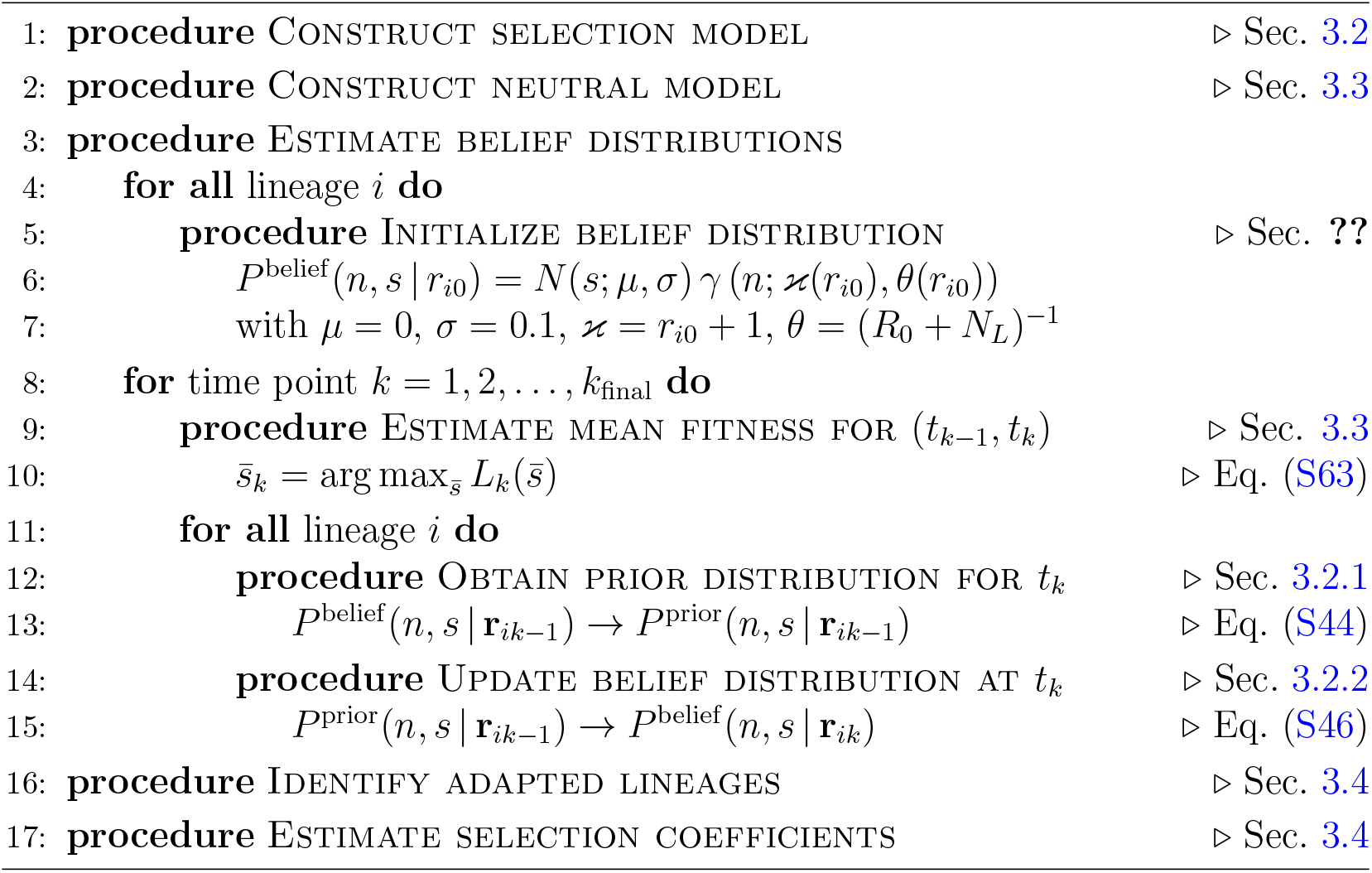

#### 3.5 Implementation

Figure 3 in the main text and Algorithm 1 show the overall BASIL workflow. We implement BASIL in a Python-based software package. For MCMC sampling, this package uses the C library *stan* and the Python package *pystan 2* as an interface between Python and *stan*. MCMC sampling is implemented using the No-U-Turn Sampler (NUTS) sampler, which efficiently generates proposals based on the posterior distribution [5]. For each belief distribution, we obtain *M* = 3500 samples, with a burn-in of 1,000 steps. For the maximum likelihood estimation of mean fitness, we use the Python package *Noisyopt* for the optimization of noisy functions. Noise arise from the fact that we estimate 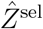 and 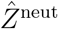 for every lineage using a Monte Carlo method as described in Section 3.3. The *Noisyopt* package repeatedly evaluates the stochastic target function, then averages over the stochasticity to ensure convergence.

The BASIL analysis of a single BLT experiment with 5 *×* 10^5^ lineages sampled at 10 time points took us about 50 hours to complete on AMD Ryzen 5 7600X 6-Core Processor, and we found that the runtime increases linearly in the number of lineages and sampling time points. However, since MCMC samples for different lineages are independent at each time point, our algorithm is easily parallelizable, and a multiprocessing capability has been implemented in BASIL. The number of processors can be set manually depending on the user’s environment. In our analyses, we found it convenient to use 12 to 32 processor cores.

BASIL code, example data and installation instructions are available at https://github.com/HuanyuKuo/BASIL-public

When barcode read count data are provided, the algorithm automatically detects the total number of barcodes and the cycle duration between sampling times. Other parameters can be adjusted in myConstant.py. These include:

- Biological settings: dilution factor, carrying capacity, and the number of randomly chosen reference lineages for mean fitness estimation.
- System settings: number of processors (CPU cores) used for parallel computation.

For adapted lineage calling, we use a confidence factor of *β* = 3.3 as the default setting in this work. However, users may adjust *β* to explore the sensitivity of lineage calls (see Discussion in the main text and Figure S12). Since *β* only affects the final lineage-calling step, changing its value does not require re-running the time-consuming MCMC estimation.

After completing a run, BASIL generates the following output files:

- Bayesian_global_parameters_XXX: inferred mean fitness trajectory and inferred *ϵ* trajectory.
- BASIL_Selection_Coefficient_for_called_Adapted_XXX: list of adapted lineages, including barcode indices, estimated selection coefficients (*s*) with mean and standard deviation, and the calling time.
- posterior_XXX_SModel_S_T1: the parametric belief distributions of all lineages at a particular sampling time point.
- glob_XXX_T1: information of maximizing the likelihood of the mean fitness and *ϵ* at a particular sampling time point.

We recommend that users first run BASIL on the provided example BLT dataset. This test run helps verify that the installation is working properly and familiarizes users with the workflow before applying the algorithm to their own data.

### 4 Application of BASIL to simulated data

#### 4.1 Belief distribution and lineage identification

Figure S4 shows how the belief distribution 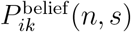 changes over time as information about the lineage frequency is accrued. Next, as discussed in the main text, we used simulated data to determine the optimal value of the confidence factor *β*, setting *β* = 3.3. The confidence factor has a simple interpretation. We use a linear classifier to classify lineages into adapted and neutral. Specifically, if we plot the inferred selection coefficient of each lineage *ŝ*_*i*_ and the estimated standard deviation 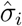 of the credible interval around it, then 1*/β* is the slope of the line that separates adapted from non-adapted lineages (see Figures S5). We find that lineages form two large clusters (with some finer clustering structure visible in Figures S5B) that contain overwhelmingly adapted or non-adapted lineages, respectively. The clas-sification line 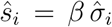 with *β* = 3.3 separates these two clusters very well (see Table 1 in the main text).

#### 4.2 Performance on simulated data

To analyze the performance of BASIL on lineage identification, we calculate the following standard statistics:

- The number of true positives (TP), i.e., adapted lineages identified as such;
- The number of false positives (FP), i.e., neutral lineages incorrectly identified as adapted;
- The number of true negatives (TN), i.e., neutral lineages identified as such;
- The number of false negatives (FN), i.e., adapted lineages incorrectly identified as neutral;
- Precision, i.e., the fraction of all positives that are true, TP/(TP+FP);
- Recall, or the true positive rate, i.e., the fraction of all adapted lineages that are identified as such, TP/(TP+FN);
- The F1-score, which is the harmonic mean of precision and recall.

All numbers and statistics are reported in Table 1 in the main text. In particular, in the weak selection region, we obtain 31 false positives (precision 98.8%) and 430 false negatives (recall 85.7%), and in the strong selection region, we obtain 7 false positives (precision 99.7%) and 519 false negatives (recall 82.7%). Figure S5 reveals that all false positives, i.e., neutral lineages that are incorrectly identified as adapted, are very close to the classification line, suggesting that in principle their rate could be further reduced by increasing *β* (albeit at the expense of increasing the rate of false negatives). In contrast, false negatives, i.e., adapted lineages that are incorrectly classified as neutral, are distributed broadly within the non-adapted cluster, indicating that no linear classifier that is based solely on *ŝ*_*i*_ and 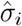 can achieve a substantially higher recall. This observation suggests that false negative lineages may be statistically indistinguishable from true neutral ones.

To investigate this conjecture, we first plotted the trajectories of all lineages stratified by their predicted and actual class labels. Figures S6 and S7 confirm that the trajectories of false negative lineages are visually indistinguishable from those of truly neutral lineages. One possible explanation for why some adapted lineages behave is if they are neutral is that these lineages have such small population sizes that their dynamics are governed largely by genetic drift, that is, they fail to “establish” [6]. An adapted lineage establishes approximately when its size *n*_*i*_ exceeds the inverse of its selection coefficient 1*/s*_*i*_, or equivalently, *n*_*i*_*s*_*i*_ *>* 1 [6]. Thus, lineages with smaller selection coefficients have a smaller chance of successfully establishing. Consistent with this prediction, we find that false negative lineages have significantly smaller selection coefficients than true positive lineages, with *P <* 0.001 in both cases (Welch’s t-test; see Figures S8). Next, for each lineage *i*, we estimated the maximum size it has achieved during the course of the simulation as 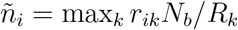 and plotted the lineage’s true selection coefficient *s*_*i*_ against *ñ*_*i*_. Figure S8 show that for the majority of false negative lineages, we have 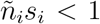, whereas for the vast majority of true positive lineages, 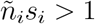, as expected. These observations support our hypothesis that some adapted lineages fail to establish. As a result, their dynamics are highly stochastic and essentially indistinguishable from neutral.

## Appendix A Expectation of the noise variance estimator and bias correction

Since the estimate (S4) we calculate its expectation to determine whether it is biased.

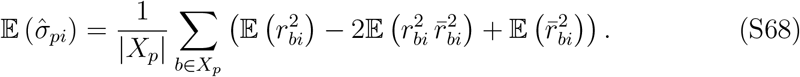

To calculate the expectations in the sum of equation (S68), note that

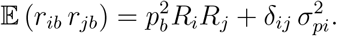

Now, we have

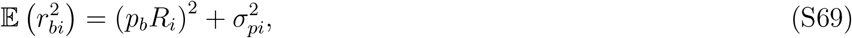

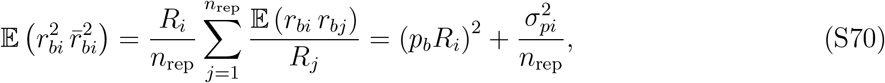

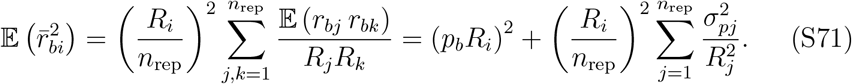

Substituting equations (S69)–(S71) into equation (S68), we obtain

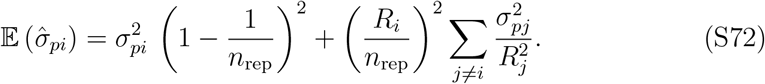

Thus, the estimate (S4) is a biased estimate of 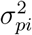. Note that if all replicates have the same coverage, *R*_*i*_ = *R*, and 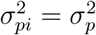, equation (S72) simplifies to

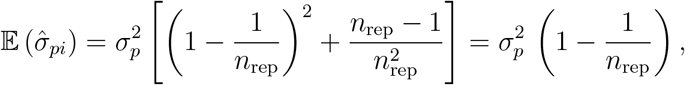

and correcting the bias is easy. To correct the bias in the general case, we rewrite equation (S72) in the matrix form as

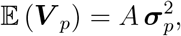

where 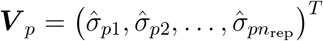 and 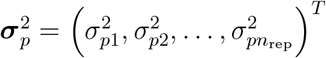 and

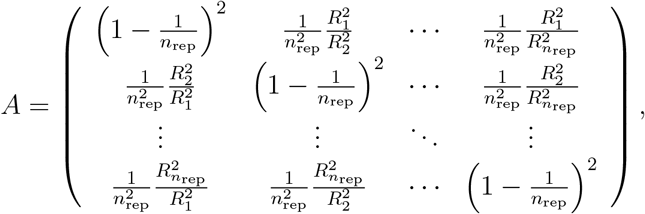

and obtain the bias-corrected estimate

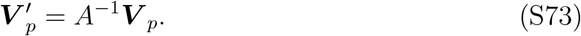

## Appendix B Expected frequency given the number of reads

Here, we derive the expressions (S22)–(S28) for the conditional expected frequency of a lineage *E* (*x*|*r*), given the number of reads *r*.

**Example 1. Poisson measurement noise and a uniform frequency distribution.** We first consider the case when the measurement process is modeled by the Poisson distribution (S21) and the frequency distribution is uniform. Then, the conditional distribution for the frequency *x* is given by

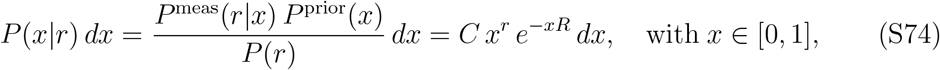

where *C* is the normalization constant. As long as *r/R* ≪ 1, this distribution is well approximated by the gamma distribution with the shape parameter *k* = *r* + 1 and scale parameter *θ* = 1*/R*. Thus, *C* = *R*^*r*+1^*/r*! whose mean is *kθ* = (*r* + 1)*/R* confirming (S22).

**Example 2. Poisson measurement noise and an exponential frequency distribution.** If the frequency distribution is exponential with mean 1*/N*_*L*_ (see equation (S23)), then we have

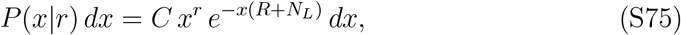

which is again is very well approximated by the gamma distribution with the shape parameter *k* = *r* + 1 and scale parameter *θ* = 1*/*(*R* + *N*_*L*_), as long as *r* ≪ *R*. Expression (S24) follows immediately.

**Example 3. Measurement noise with an increasing variance to mean ratio and a Gamma frequency distribution.** We next consider the case when the frequency distribution is Gamma with shape parameter *α* and scale parameter 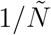 (see equation (S26)) and the read count is a negative binomial random variable with mean *xR* and the variance to the mean ratio 1 + *ϵxR* (see (S25)). When *xR* 1*/ϵ*, the negative binomial distribution (S25) converges to the Poisson distribution with mean *xR*, which implies that the conditional distribution for *x* is a Gamma generally, the conditional distribution for *x* given *r* is given by

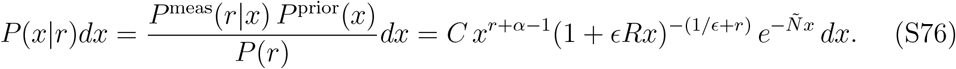

It is possible to use Laplace’s method to obtain approximate expressions for the normalization constant *C* and the moments of this distribution. Specifically, whenever *ϵ ≪* 1, we obtain

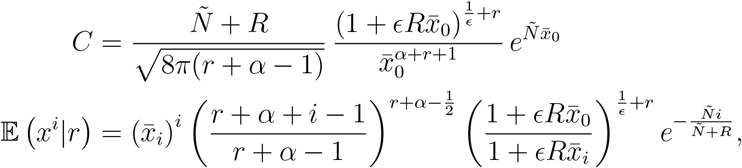

for *i, r* = 1, 2, …, where *x*_*i*_ = ^*r*+*α*+*i*−1^.

**Example 4. Measurement noise with a constant variance to mean ratio and an exponential frequency distribution.** Finally, we consider the case when the measurement process is over-dispersed but with a constant variance to the mean ratio *2κ* (see equation (S25)) and the frequency distribution is exponential (see equation (S23)). To facilitate analytical tractability, instead of tracking frequency *x* of the lineage in the population, we will track its lineage size *n*. We assume that *n* take values 1, 2, … and that its prior distribution is geometric with parameter *q*. The expected lineage size is *N/N*_*L*_, which implies that *q* = *N*_*L*_*/N*. Thus, for the conditional distribution for *n*, given *r* is

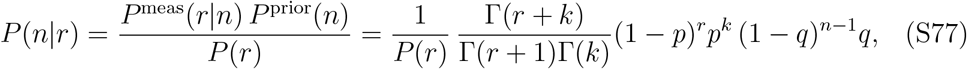

where *p* = 1/*2κ, k* = *αn, q* = *N*_*L*_*/N* and *α* = *R/N* (*2κ* − 1)^−1^. Expression (S77) suggests that the quantity 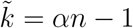 is a negative binomial random variable with parameters

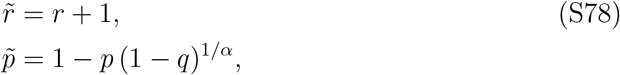

that is,

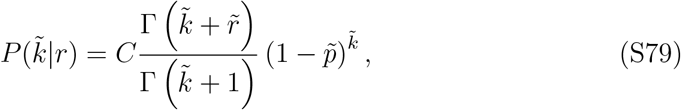

Where 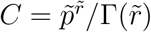 is the normalization constant. Thus, we have

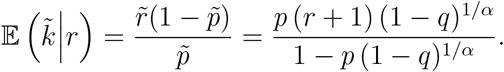

Since 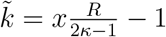, we have

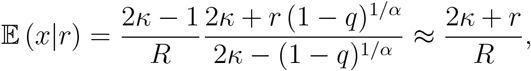

where the last approximation holds because *q* = *N*_*L*_*/N* ∼ 0.01 ≪ 1.

It will also be useful to calculate the expected number of cells in a lineage whose measured number of reads is *r*,

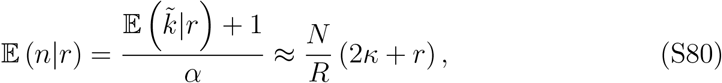

and its variance

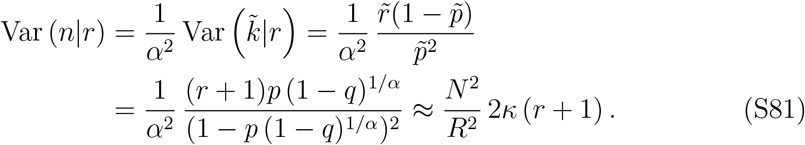

## Appendix C Expectation of the logarithm of a gamma-distributed random variable

Consider a random variable *X* that is Gamma-distributed with scale parameter *θ* and shape parameter *x*, such that its probability density is

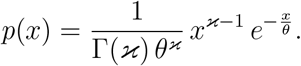

Therefore, we have

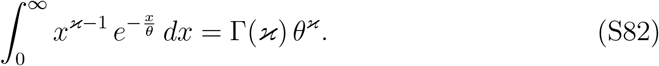

Differentiating both sides of equation (S82) with respect to *x*, we obtain

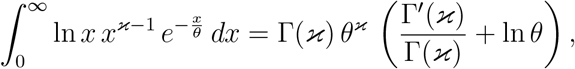

which can be rewritten as

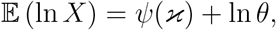

where 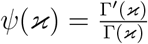 is the digamma function.

## 5 Supplementary Figures

**Figure S1.**
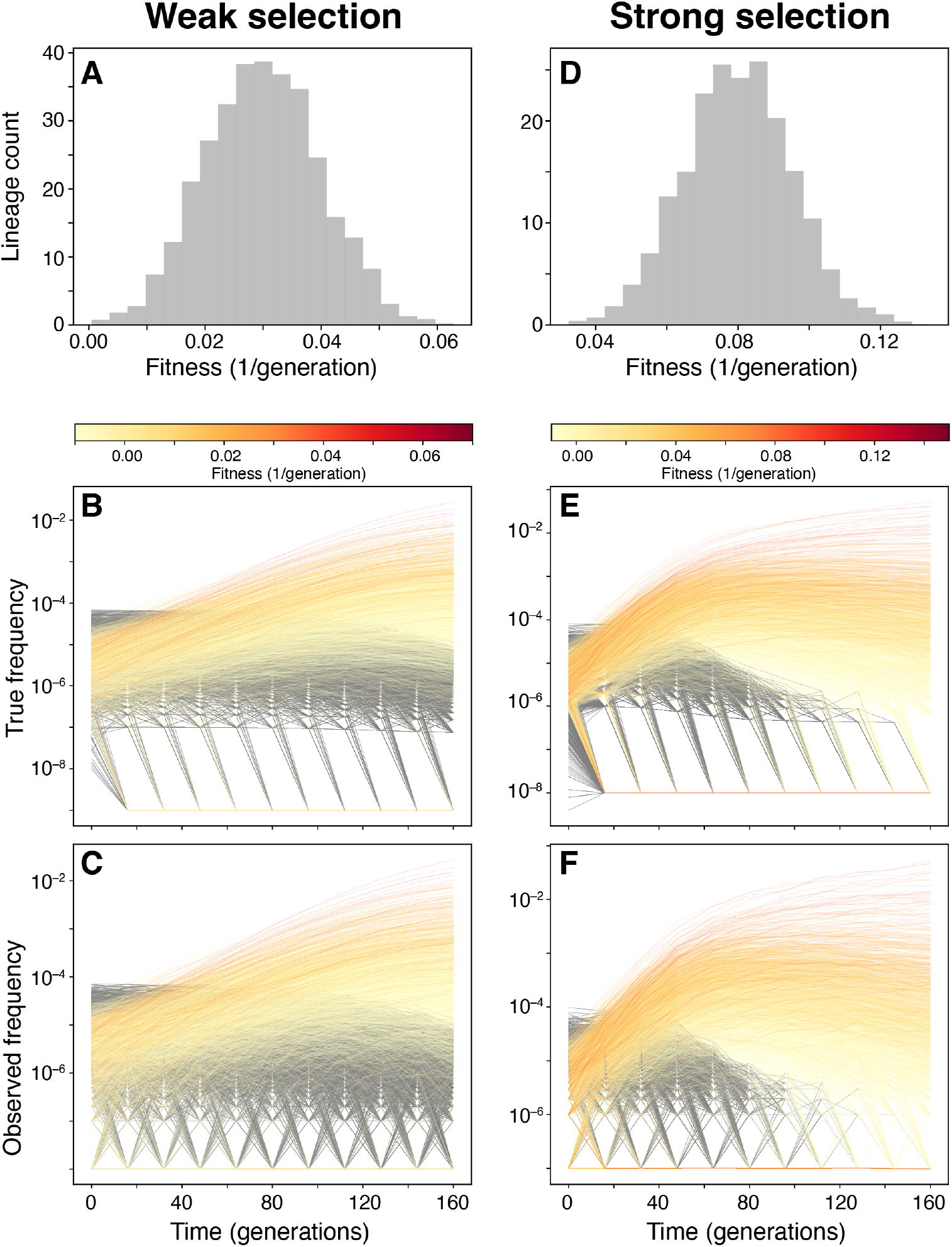
Simulated data. A–C. Weak selection regime. **D–F**. Strong selection regime. Distributions of lineage fitness (A,D). True lineage frequency trajectories (B,E). Observed barcode frequency trajectories (C,F). Adapted lineages are colored by their fitness, neutral lineages are gray.

**Figure S2.**
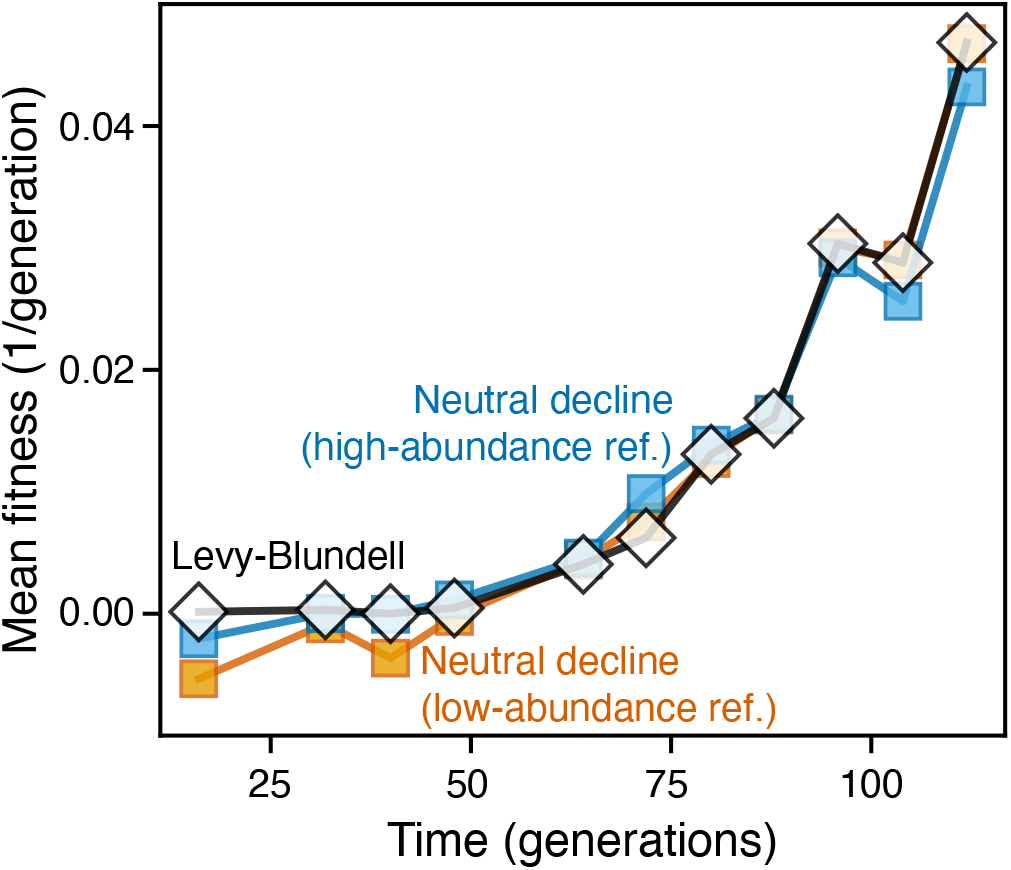
Correspondence between the full Levy-Blundell inference method and the simplified neutral decline method. Mean-fitness trajectories in Replicate 1 in the Levy 2015 dataset as reported in Ref. [3] (black line and white diamonds), and inferred by our the neutral decline method using either high-abundance lineages (blue line and squares) or low-abundance lineages (orange line and squares) as reference.

**Figure S3.**
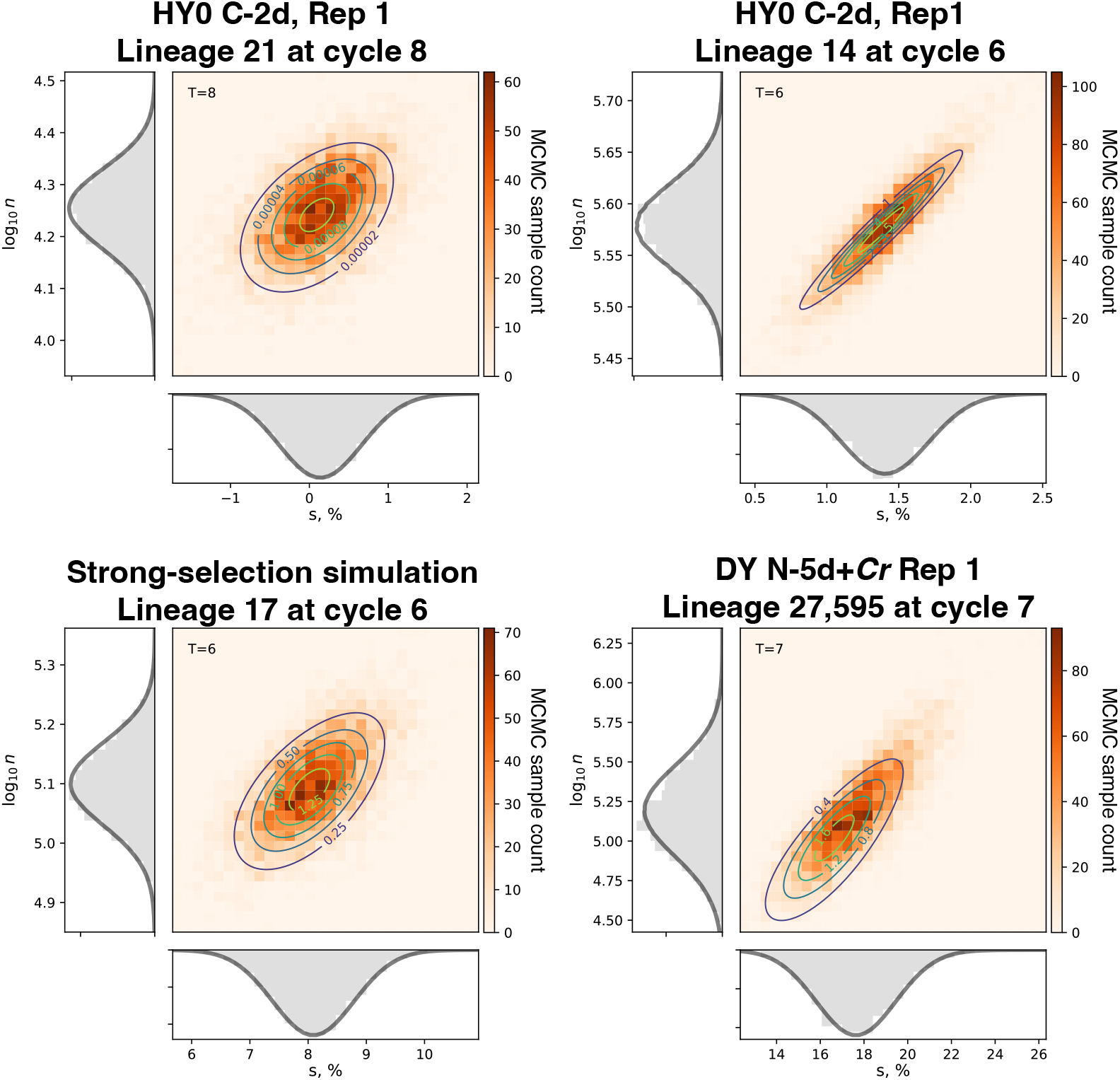
Estimation of the belief distribution *P* ^b^(*n, s*). Each panel corresponds to a single lineage in a particular dataset at a certain time point, as indicated in the panel title. The heatmap in each panel shows the number of MCMC samples in each (*s*, log_10_ *n*) bin. The contour lines correspond to the analytical function *P* ^b^(*n, s*) given by equations (S31)–(S33) with parameters fitted to the MCMC sample. Histograms show the marginal distributions for *s* (bottom) and log_10_ *n* (left) and the corresponding parametric curves. See Section 3.2.2 for details.

**Figure S4.**
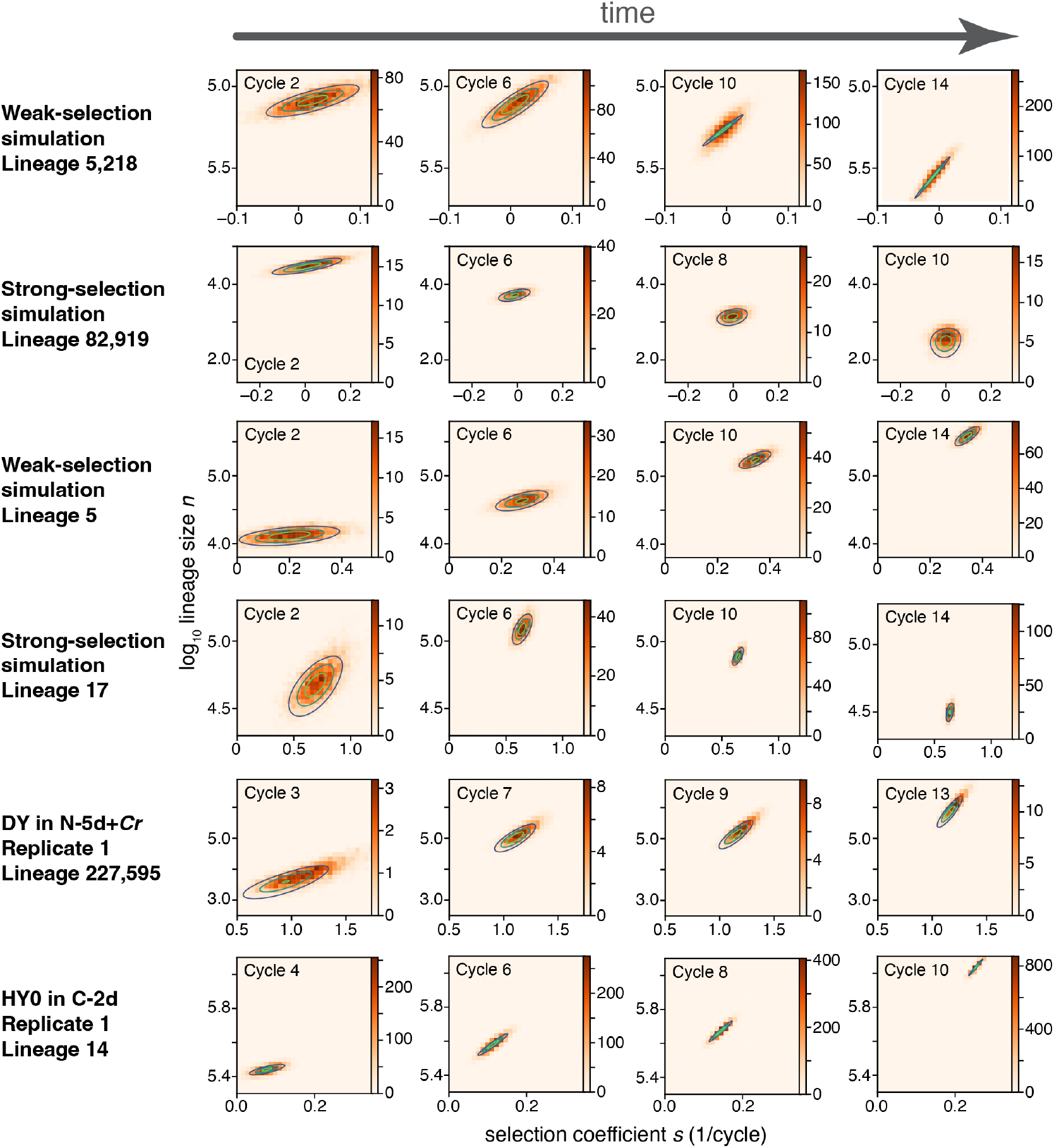
Changes in the belief distribution as data is accumulated. Each row corresponds to a single lineage in a particular dataset, as indicated on the left. Note that the selection coefficient here is shown on the per cycle basis. Notations are as in Figure S3.

**Figure S5.**
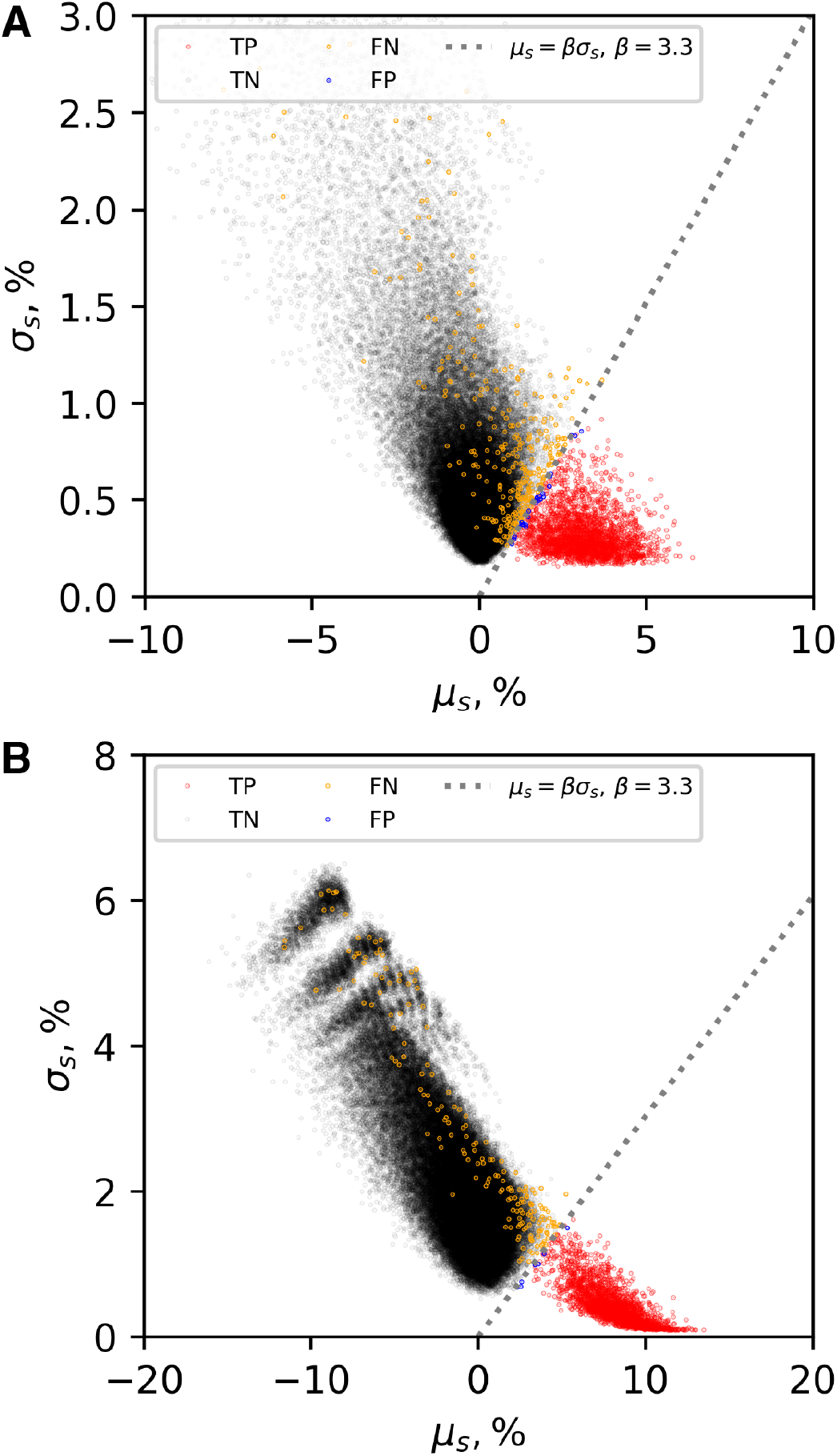
Classification of lineages in simulated data. **A**. Mean (*x* axis) versus the standard deviation (*y* axis) of the estimated marginal belief distribution for the lineage selection coefficient for simulated data in the weak selection regime. **B**. Same for the strong-selection regime. Each point represents a lineage. Colors represent different lineage classes: red = true positives, black = true negatives, orange = false negatives, blue = false positives. Dashed line is the classification line with *β* = 3.3.

**Figure S6.**
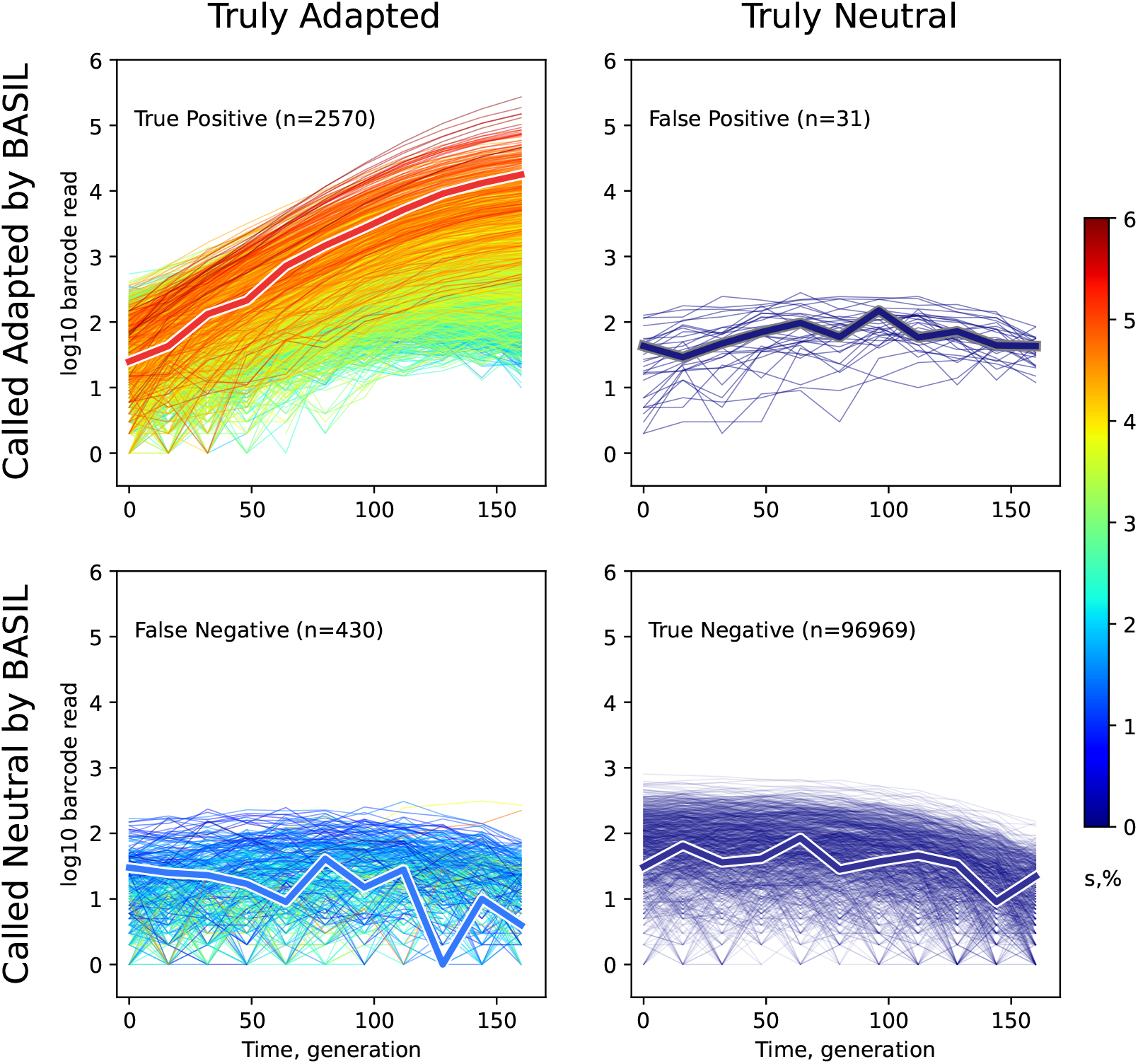
Lineage trajectories in the weak-selection simulation stratified by class. Data is the same as in Figure S1B. Only 970 random true negative lineages are shown to improve visualization. Trajectories are colored by lineage fitness.

**Figure S7.**
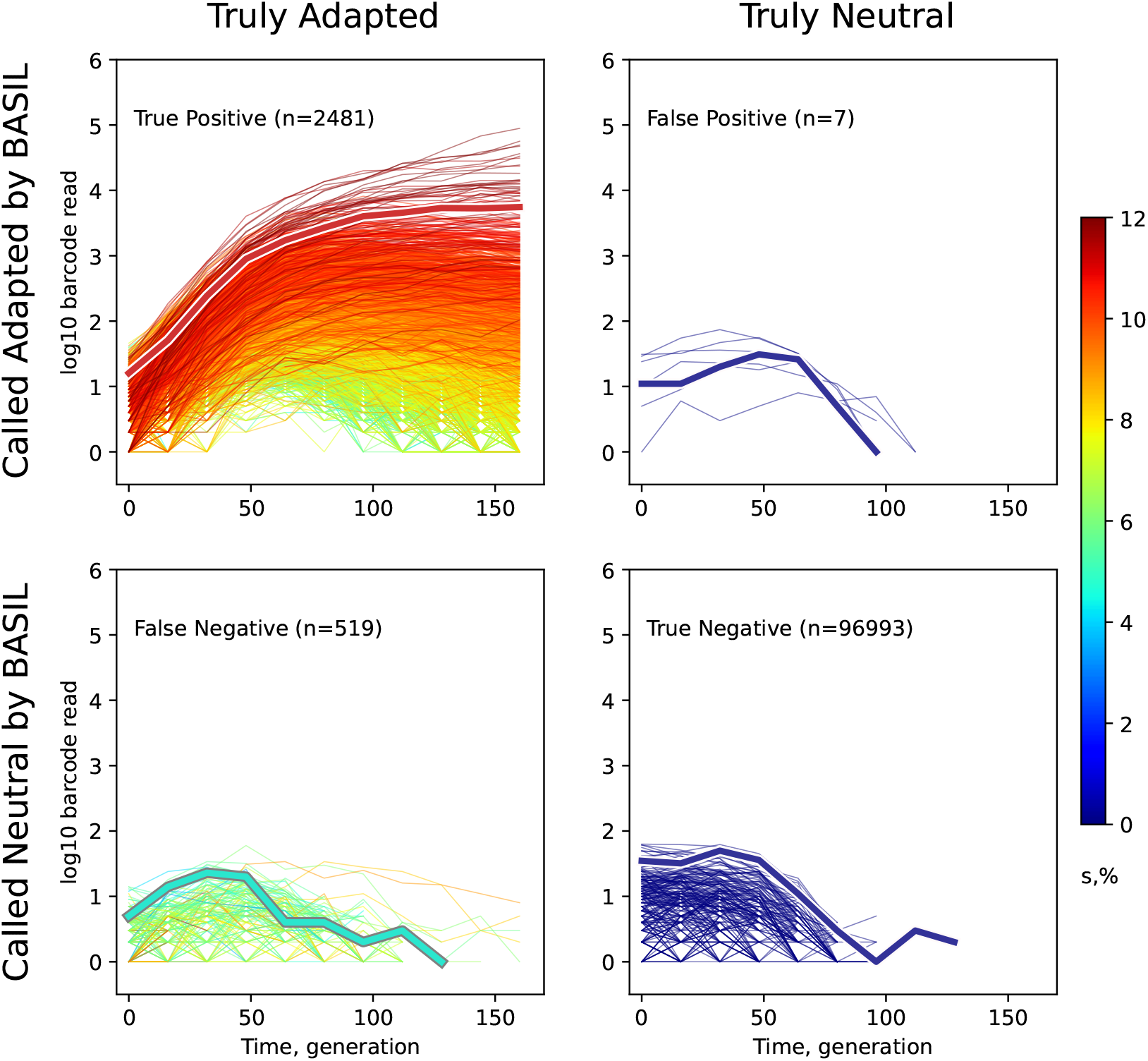
Lineage trajectories in the strong-selection simulation stratified by class. Data is the same as in Figure S1E. Only 243 random true negative lineages are shown to improve visualization. Trajectories are colored by lineage fitness.

**Figure S8.**
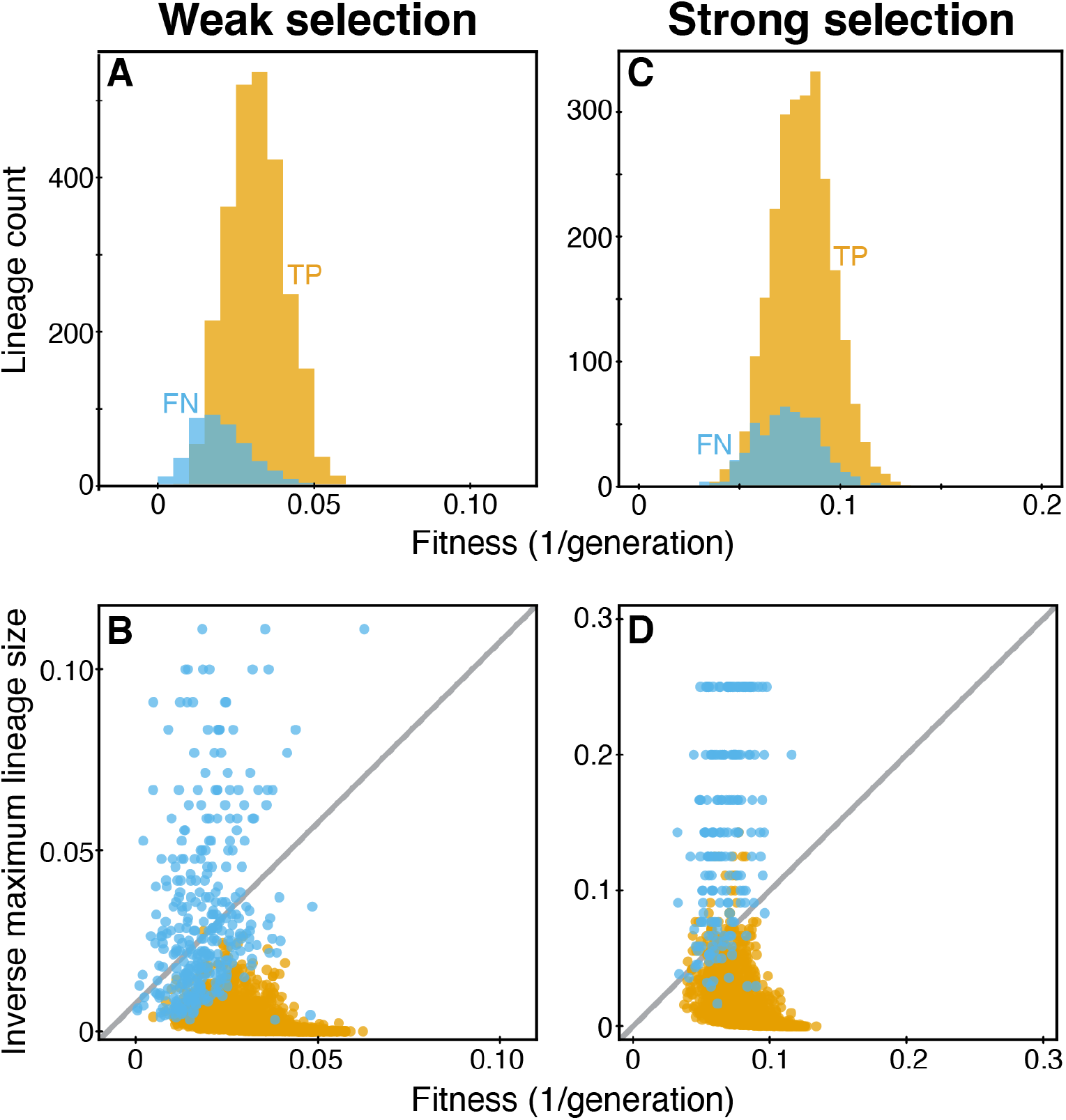
True positive and false negative lineages have distinct features in simulated data. **A**. Distribution of selection coefficients of true positives (yellow) and false negative (blue) lineages in the weak-selection simulation. **B**. The relationship between lineage fitness *s*_*i*_ and its maximum size *ñ*_*i*_ in the simulation (see Section 4.2 for details). Lineages above the diagonal (gray line) are those with *ñ*_*i*_*s*_*i*_ *<* 1, and lineages below the diagonal are those with *ñ*_*i*_*s*_*i*_ *>* 1. **C**. Same as panel A but for the strong-selection simulation. **D**. Same as panel B but for the strong-selection simulation.

**Figure S9.**
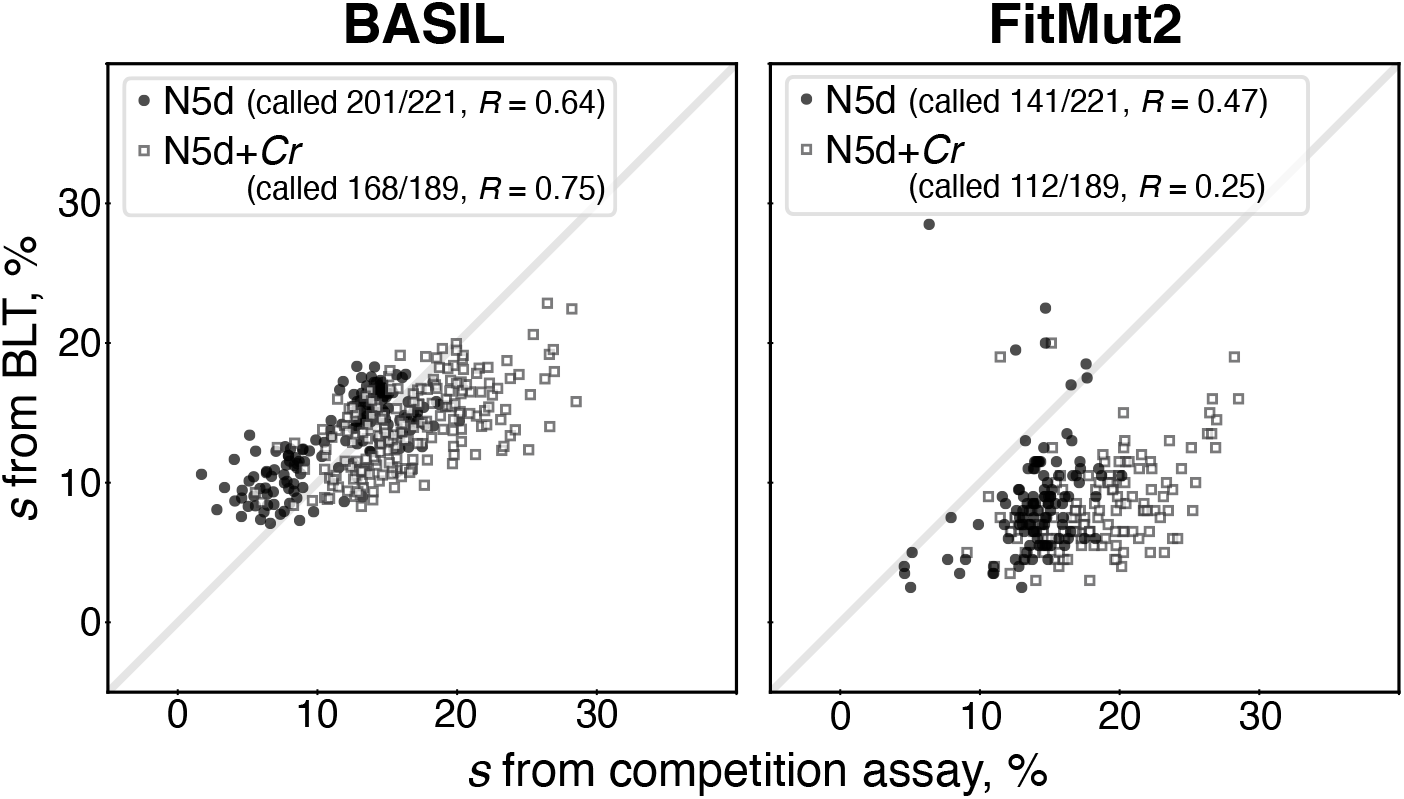
Comparison of BASIL and FitMut2 performance on Venkataram 2023 data. Lineage fitness inferred from BLT experiments by either BASIL or FitMut2 is plotted against the fitness of respective isolated clones measured in competition assays by Venkataram et al [7]. Filled circles represent lineages/clones measured in the N5d condition, which Venkataram et al refer to as “Alone”. Empty squares represent lineages/clones measured in the N5d+*Cr* condition, which Venkataram et al refer to as “Community”. The number of lineages called as adapted in the respective BLT experiment as well as Pearson correlation coefficient *R* are shown in parenthesis.

**Figure S10.**
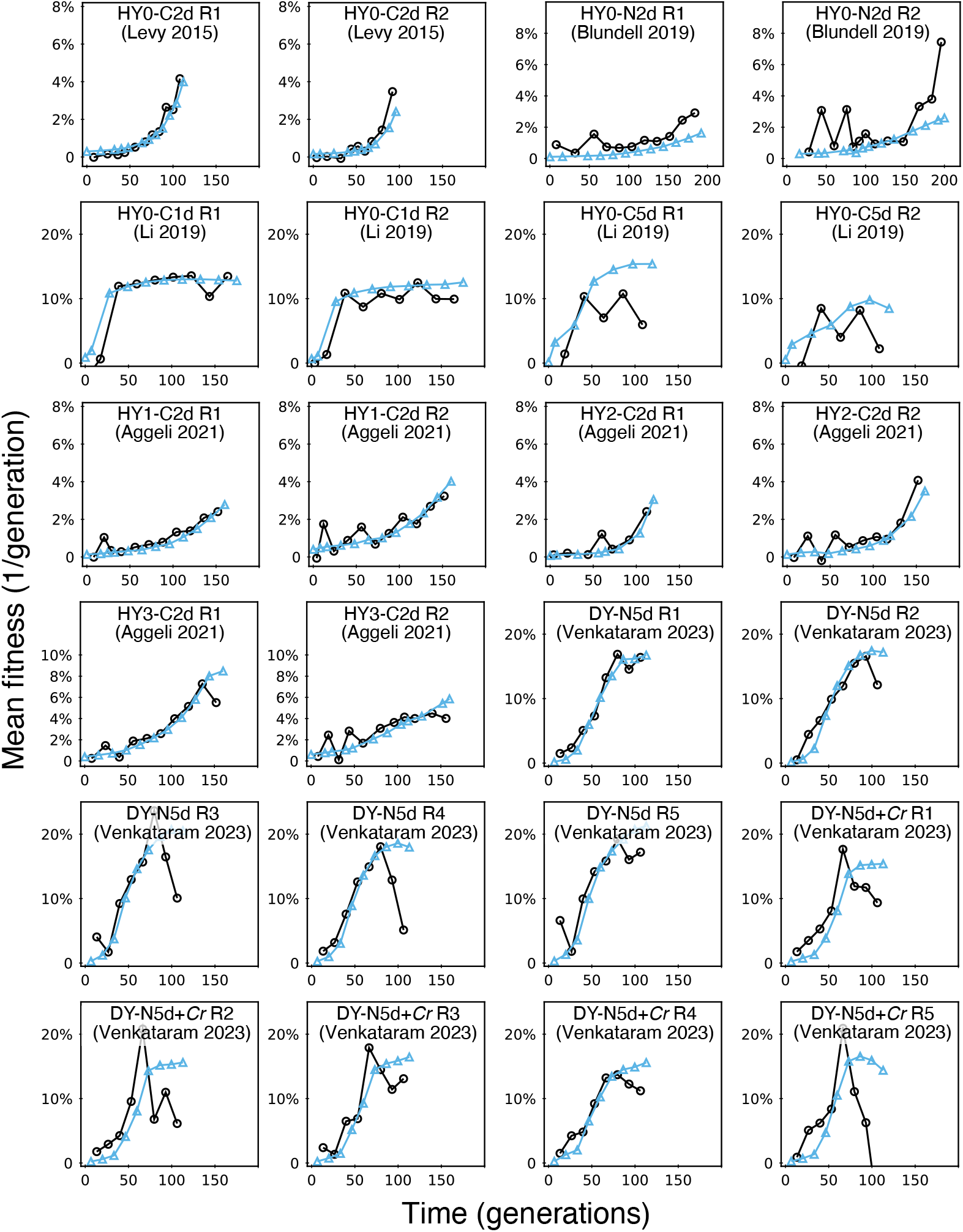
Mean fitness trajectories in published BLT datasets. Each panel represents a replicate of a BLT experiment, as indicated. Black lines show trajectories inferred by BASIL; blue lines show trajectories calculated from lineages identified as adapted.

**Figure S11.**
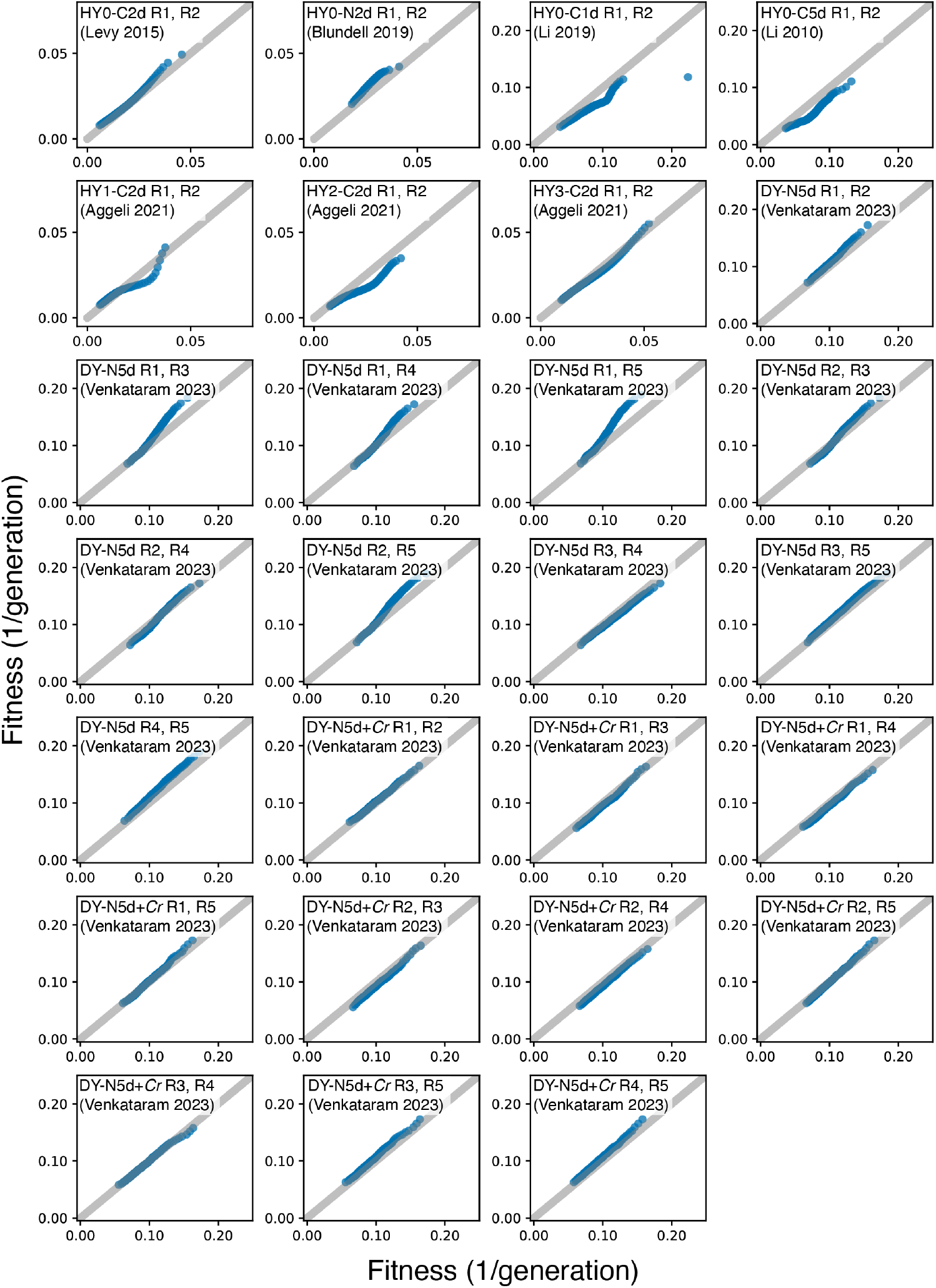
Quantile-quantile plots for the measured distributions of fitness effects (mDFEs) inferred across replicates. Each panel represents a pair of replicates from the same BLT study, as indicated. 100 quantiles are shown in each panel.

**Figure S12.**
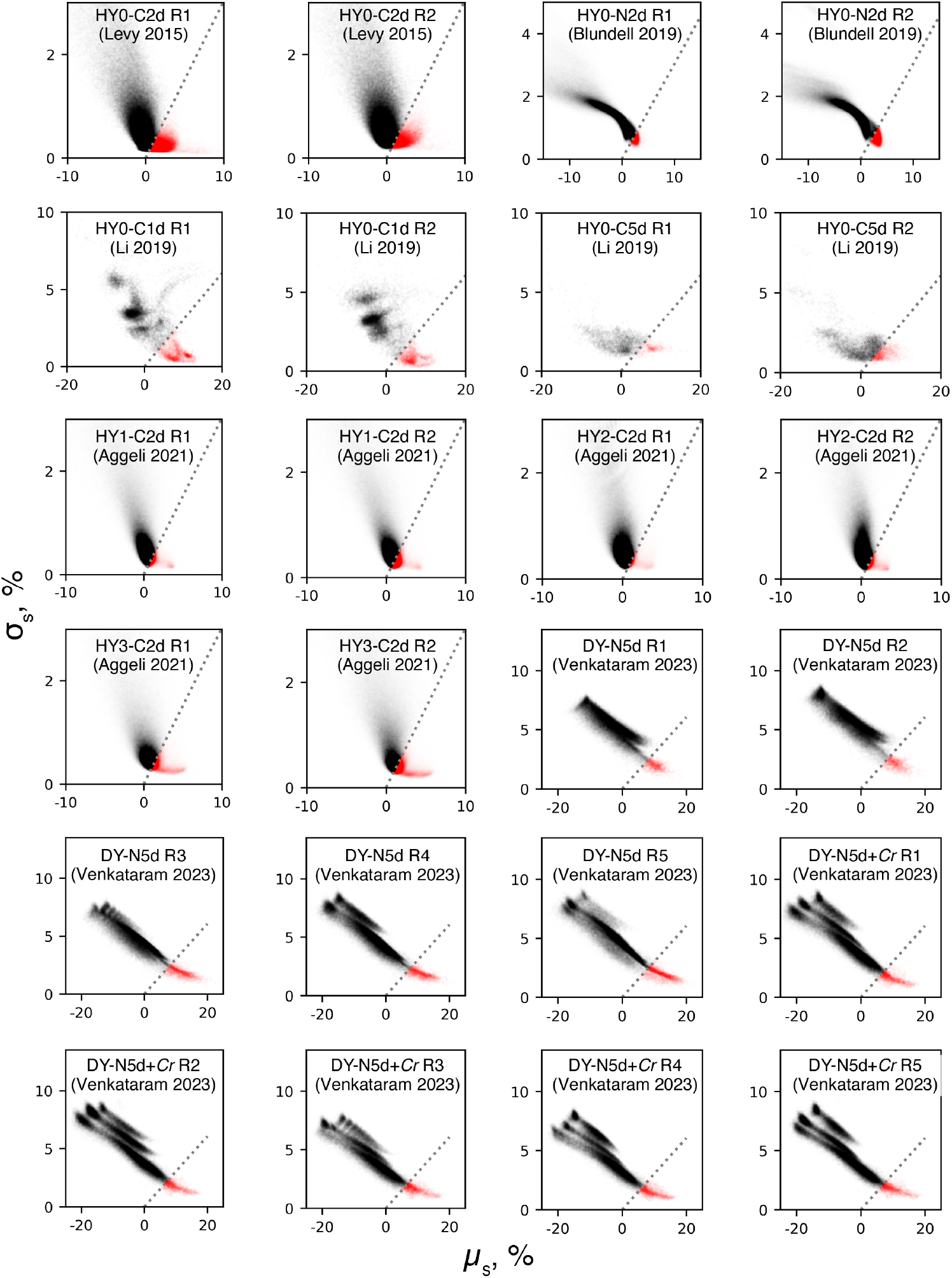
Classification of lineages in real BLT data. Each panel represents a replicate of a BLT experiment, as indicated. Notations are as in Figure S5.

## 6 Supplementary Tables

**Table S1.** Model fitting for the relationship between read-count mean and variance. ^1^Residual sum of squares.

| | Linear ($\epsilon = 0$) | | Quadratic ($\epsilon \neq 0$) | |
| --- | --- | --- | --- | --- |
| | Poisson<br>($a = 1$) | General<br>– | 1-param<br>($a = 1$) | General<br>– |
| Fitted $\hat{a}$ | – | 4.34 | – | 1.97 |
| Fitted $\hat{\epsilon}$ | – | – | 0.0024 | 0.0016 |
| RSS<sup>1</sup> | 173.6 | 85.2 | 55.1 | 48.3 |
| $R^2$ | 0.77 | 0.89 | 0.93 | 0.94 |

**Table S2.** Goodness of fit comparison between models fitting the relationship between read-count mean and variance. In all cases *n* = 41.

| Null model | | | Alternative model | | | $F$ | $P$-value |
| --- | --- | --- | --- | --- | --- | --- | --- |
| Name | $p_{\text{null}}$ | RSS<sub>null</sub> | Name | $p_{\text{alt}}$ | RSS<sub>alt</sub> | | |
| Poisson | 0 | 173.6 | Lin. Gen. | 1 | 85.2 | 41.5 | $1.13 \times 10^{-7}$ |
| Poisson | 0 | 173.6 | Quad. 1-param. | 1 | 55.1 | 50.7 | $1.63 \times 10^{-11}$ |
| Poisson | 0 | 173.6 | Quad. Gen. | 2 | 48.3 | 86.0 | $1.44 \times 10^{-11}$ |
| Lin. Gen. | 1 | 85.2 | Quad. Gen. | 2 | 48.3 | 29.9 | $2.87 \times 10^{-6}$ |
| Quad. 1-param. | 1 | 55.1 | Quad. Gen. | 2 | 48.3 | 5.56 | 0.0235 |

**Table S3.** Simulation parameters. For the weak selection regime, we chose the relationship between mean read count ⟨*r*_*i*_⟩ and variance in read count 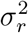 to be linear in order to make our simulations as similar to the work by Levy et al [3] as possible. For the strong selection regime, we chose the more realistic quadratic relationship, based on our results.

| Parameter | Selection |  |
| --- | --- | --- |
|  | Weak | Strong |
| $\langle s \rangle$ | 0.03 | 0.08 |
| $\sigma_s$ | 0.01 | 0.015 |
| $\langle N \rangle$ | $2.56 \times 10^9$ | $2.56 \times 10^8$ |
| $\langle N_b \rangle$ | $10^7$ | $10^6$ |
| $D$ | 256 | 256 |
| $R$ | $10^7$ | $10^6$ |
| $\sigma_r^2$ | $6 \langle r_i \rangle$ | $\langle r_i \rangle + 0.01 \langle r_i \rangle^2$ |

**Table S4.** Summary of BLT methods.

|  | **Levy-Blundell**<br>**(LB method)** | **FitMut2**<br>**(modified LB)** | **BASIL**<br>**(this work)** |
| --- | --- | --- | --- |
| Evolution model | Branching process of population size from a single cell | same as the LB method | Exponential growth and dilution in batch culture |
| Barcode measuring model | Noise model with a linear meanvariance. parameter: $\kappa$ | same as the LB method | Negative binomial with a quadratic meanvariance. parameter: $\epsilon$ |
| Probability of $n$ trajectory | Not applied | Not applied | The (Bayesian) belief distribution Sec.3.2 |
| Probability of $r$ trajectory | Noise model description Eq. (S8) | Noise model description with both adapted and neutral models Eq. (S12) | Marginal probability of read count Eq. (S47) |
| Mean-fitness estimation, $\bar{s}$ | Maximizing the likelihood function from the reference lineages | Direct calculation from the adapted lineages | Maximizing the likelihood function from the reference lineages |
| Choice of reference lineage | All small lineages with $20 < r < 40$, assuming they are putatively neutral | Recursively select all putatively adapted lineages | Randomly select 3000 lineages, then assign them to putatively neutral or adapted group |
| Neutral decline assumption | Yes | No | No |
| Lineage identification | Ratio of posterior probabilities of adapted and neutral models Eq. (S11) | Bayesian probability of model choice Eq. (S13) | Uncertainty of $s$ in belief distribution Sec.3.4 |
| Fitness effect inference, $s$ | Maximizing the likelihood function for $r$ trajectory Eq. (S10) | Maximizing the likelihood function for $r$ trajectory | The mean of $s$ in belief distribution Sec.3.4 |

**Table S5.** Notations used in BASIL.

| Notation | Description | Eqns |
| --- | --- | --- |
| $k$ | sampling time index, $k \in [0, k_f]$ | |
| $t_k$ | time (in generation) of the $k$th sample | |
| $R_k$ | total read depth at $t_k$ | |
| $D$ | dilution ratio, $D > 1$ | |
| $N$ | population size before dilution | |
| $N_b$ | population size after dilution, $N_b = N/D$ | |
| $\epsilon$ | measurement error parameter | (2), (??) |
| $s_i$ | selection coefficient (fitness) of the $i$th lineage | |
| $\bar{s}_k$ | mean fitness of the population in time interval $(t_{k-1}, t_k)$ for all $k \geq 1$ | |
| $n_{ik}$ | size of $i$th lineage at $t_k$ before dilution | |
| $r_{ik}$ | barcode read count of $i$th lineage at $t_k$ | |
| $\mathbf{r}_{ik}$ | observation vector for lineage $i$ prior to and including $t_k$, $\mathbf{r}_{ik} = (r_{i0}, \dots, r_{ik})$ | |
| $P^{\text{belief}}(n, s\vert \mathbf{r})$ | belief probability that a lineage with the observation vector $\mathbf{r}$ has size $n$ and fitness $s$ | (3), (S46) |
| $P_{ik}^{\text{belief}}(n, s)$ | shorthand for $P^{\text{belief}}(n, s\vert \mathbf{r}_{ik})$ | |
| $P^{\text{meas}}(r\vert n)$ | probability of observing $r$ reads for a lineage with size $n$. $P^{\text{meas}}$ depends on the total read depth $R$ and the measurement error parameter $\epsilon$ | (S30) |
| $P^{\text{prior}}(n, s\vert \mathbf{r}')$ | prior probability that a lineage with the observation vector $\mathbf{r}'$ has size $n$ and selection coefficient $s$ at the *next* sampling time point. $P^{\text{prior}}$ depends on the current mean fitness $\bar{s}$ | (S44) |
| $Z^{\text{neut}}(r_{ik})$ | marginal likelihood of $r_{ik}$ in the neutral model. $Z^{\text{neut}}$ depends on the current mean fitness $\bar{s}$ and measurement error parameter $\epsilon$ | (S66) |
| $Z^{\text{sel}}(r_{ik})$ | marginal likelihood of $r_{ik}$ in the selection model. $Z^{\text{sel}}$ depends on the current mean fitness $\bar{s}$ and measurement error parameter $\epsilon$ | (S47) |
| $L_k(\bar{s}, \epsilon)$ | log-likelihood function for the mean fitness $\bar{s}$ and measurement error parameter $\epsilon$ | (S63) |

**Table S6.**
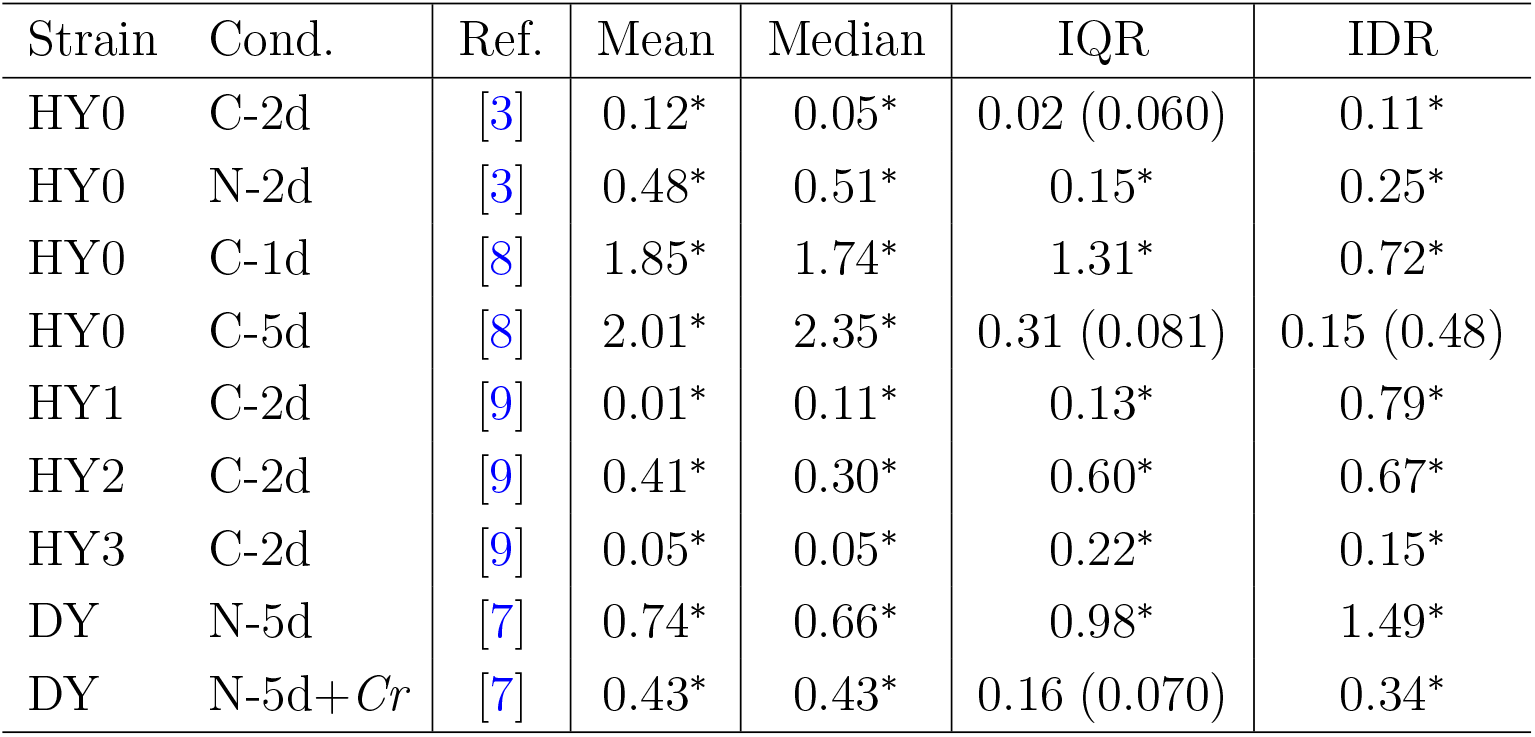
Differences of mDFE statistics across replicates. For each mDFE statistic, we report the absolute value of the pairwise difference in the statistic value, averaged across all pairs of replicates (see text for details). ^*^ indicates *P*-value *<* 0.01 (permutation test); *P*-values *≥* 0.01 are shown in parentheses.

**Table S7.** Effects of down-sampling. The “Full” dataset is Replicate 1 of the HY0 strain in C-1d environment [3]. See Materials and Methods in the main text for details. ^1^Pearson correlation coefficient. ^2^Averaged absolute difference of *s* inferred in Full and Reduced data.

|  |  | **BASIL** | **FitMut2** |
| --- | --- | --- | --- |
| Full data | # of adapt. lin. | 60920 | 15386 |
| Reduced coverage | # of adapt. lin. | 49330 | 1131 |
|  | Change in calling rate | −19.0% | −98.1% |
|  | Overlap | 48738 | 1130 |
|  | Relative overlap | 98.8% | 99.9% |
| | $s$ PCC<sup>1</sup> | 95.8% | 82.0% |
| | Error<sup>2</sup> in $s$ | 0.20% | 0.83% |
| Reduced sampling | # of adapt. lin. | 48206 | 55248 |
|  | Change in calling rate | −20.9% | +259.1% |
|  | Overlap | 48177 | 15381 |
|  | Relative overlap | 99.9% | 100, 0% |
| | $s$ PCC<sup>1</sup> | 97.9% | 62.0% |
| | Error<sup>2</sup> in $s$ | 0.18% | 1.29% |

